# Experimental Evolution of *Pseudomonas putida* under Silver Ion versus Nanoparticle Stress

**DOI:** 10.1101/2020.09.17.302794

**Authors:** Feng Dong, Ana C. Quevedo, Xiang Wang, Eugenia Valsami-Jones, Jan-Ulrich Kreft

## Abstract

Whether the antibacterial properties of silver nanoparticles (AgNPs) are simply due to the release of silver ions (Ag^+^) or, additionally, nanoparticle-specific effects, has been debated for over a decade. We used experimental evolution of the model environmental bacterium *Pseudomonas putida* to ask whether bacteria respond differently to Ag^+^ or AgNP treatment. We pre-evolved five cultures of strain KT2440 for 70 d without Ag to reduce confounding adaptations before dividing the fittest pre-evolved culture into five cultures each, evolving in the presence of low concentrations of Ag^+^, well-defined AgNPs or Ag-free controls for a further 75 d. The mutations in the Ag^+^ or AgNP evolved populations displayed different patterns that were statistically significant. The non-synonymous mutations in AgNP-treated populations were mostly associated with cell surface proteins, including cytoskeletal membrane protein (FtsZ), membrane sensor and regulator (EnvZ and GacS) and periplasmic protein (PP_2758). In contrast, Ag^+^ treatment selected for mutations linked to cytoplasmic proteins, including metal ion transporter (TauB) and those with metal binding domains (ThiL and PP_2397). These results suggest the existence of AgNP-specific effects, either caused by sustained delivery of Ag^+^ from AgNP-dissolution, more proximate delivery from cell-surface bound AgNPs, or by direct AgNP action on the cell’s outer membrane.

**Originality-Significance Statement:** The increasing use of silver nanoparticles (AgNPs) and their release into the environment may affect environmental microorganisms and their communities and evolution. It has long been debated whether the toxicity of AgNPs towards microorganisms is solely due to their dissolution into toxic Ag^+^ or whether distinct nanoparticle related toxicity exists. We set up an evolution experiment to explore the adaptation of the environmental model bacterium *Pseudomonas putida* to Ag^+^ versus AgNP stress in order to elucidate the potentially different toxicity mechanisms of ionic and nanoparticulate Ag. We found novel mutations and distinct mutation patterns under Ag^+^ and AgNP treatment by whole genome sequencing. Our work highlights the association of the mutations selected by Ag^+^ stress with metal ion metabolism inside the cells and the mutations specific to AgNP stress with the cell’s surface. The finding that *P. putida* cells evolved in different directions under selection by Ag^+^ and AgNPs demonstrates a need for assessing the toxicity of nanomaterials separately in any environmental risk assessments.

## Introduction

Silver (Ag) is not known to be an essential element for any organism. Instead, it is toxic to microorganisms, viruses, plants and animals and has been used for ∼3,000 years as an antimicrobial agent to preserve food and control infections (Russell and Hugo, 1994). The antimicrobial activities of Ag^+^ result primarily from its high affinity to a variety of biomolecules (Eckhardt *et al*., 2013). Silver nanoparticles (AgNPs) have received increasing interest for antimicrobial applications in many fields, such as water disinfection (Sankar *et al*., 2013), textiles (Janković and Plata, 2019) and infection control in medical settings (Chaloupka *et al*., 2010; Cui *et al*., 2015). Given their toxicity, AgNPs pose potential ecological risks once they are discharged into the environments (Nowack *et al*., 2011; Pourzahedi and Eckelman, 2015).

It has been debated over a decade whether AgNPs have the same bactericidal effects as Ag^+^ (Stabryla *et al*., 2018). As AgNPs hardly penetrate bacterial cells due to the physical barrier of the peptidoglycan sacculus and the lack of endocytosis mechanisms enabling nanoparticle uptake by eukaryotes, several studies suggested that the release of Ag^+^ from AgNPs fully accounts for the toxicity of AgNPs in bacteria (Hsueh *et al*., 2015) and microbial communities (Gwin *et al*., 2018). Using anoxic conditions to minimize the dissolution of Ag^+^ from AgNPs, no *direct* effects of silver nanoparticles were observed in *Escherichia coli* (Xiu *et al*., 2012). However, it has been reported that silver nanoparticles enhance bactericidal effects through attaching to the bacterial cell wall, releasing Ag^+^ at the cell surface and delivering high concentrations of Ag^+^ into the cells (McQuillan *et al*., 2012; Bondarenko *et al*., 2013). In support of this, measuring intracellular Ag^+^ concentrations suggested that it is the intracellular Ag^+^ that ultimately causes the toxicity of AgNPs (Ivask *et al*., 2009; Bondarenko *et al*., 2013). Moreover, using a genome-wide library of knockout mutants of *E. coli* (the Keio collection), it was demonstrated that Ag^+^ and AgNP treatments involve different toxicity pathways (Ivask *et al*., 2014). Similarly, transcriptomic and proteomic studies in *E. coli* and *Pseudomonas aeruginosa* identified differential gene regulation and protein production upon exposure to Ag^+^ or AgNPs (McQuillan and Shaw, 2014; Yan *et al*., 2018). The generation of reactive oxygen species (ROS) that is enhanced by AgNPs has also been reported as a nanoparticle-specific effect (He *et al*., 2012; Xu *et al*., 2012). Together, these studies suggest that AgNPs have distinct effects in the presence of oxygen. We suggest that the continuing debate can be resolved by distinguishing *direct* nanoparticle effects from nanoparticle-*specific* effects and realizing that anoxic conditions can only rule out direct effects. Indeed, oxic conditions, which lead to dissolution of AgNPs, are required for the manifestation of any differences due to the way in which Ag^+^ is delivered to the cells. For example, the concentration of free silver ions, added to cultures at the start, will decline as they bind to the cells and media components, an effect that will depend critically on cell density and environmental chemistry (Dong *et al*., 2017; Faiz *et al*., 2018). In contrast, AgNPs will continuously release new Ag^+^ and the nanoparticle-specific kinetics of Ag^+^ release and delivery can potentially alter the way in which cells are affected (Dong *et al*., 2017). Considering that Ag binds non-selectively to a broad range of biomolecules, it is generally accepted that evolution of Ag resistance in bacteria should be difficult (Medici *et al*., 2019), yet Ag resistance has been reported under prolonged Ag exposure such as in Ag-treated burn wounds (Larkin Mchugh *et al*., 1975) and in silver mines (Klaus *et al*., 1999), where bacteria can tolerate Ag^+^ at concentrations up to 1,080–5,400 mg/L. These minimum inhibitory concentrations (MICs) are much higher than those for wild type strains (6.5–32 mg/L) (Harrison *et al*., 2004), demonstrating that bacteria are capable of evolving Ag resistance. Prolonged exposure of *Bacillus* sp. to either metallic silver or silver oxide nanoparticles selected Ag resistance of up to 10 mg/L (Gunawan *et al*., 2013), but it is not clear whether the bacteria adapted to Ag^+^ or AgNPs and by which mutations. Known Ag resistance mechanisms target Ag^+^, including efflux (Gupta *et al*., 1999; Stoyanov *et al*., 2003), reduction to metallic Ag (Klaus *et al*., 1999), or chelation (Sedlak *et al*., 2012; Asiani *et al*., 2016). An evolution experiment with *E. coli* under AgNP stress showed that resistance can readily evolve (Graves *et al*., 2015). In another study, two strains of *E. coli* and one strain of *Pseudomonas aeruginosa* evolved AgNP-specific resistance by flagellin facilitated aggregation of AgNPs; this resistance by avoidance was stable over many passages of the cultures though no changes in coding sequences were observed (Panáček *et al*., 2018). Both studies did not compare evolution under AgNP stress with evolution under Ag^+^ stress so it remains unclear whether nanoparticle-specific toxicity pathways were responsible.

In this study, we hypothesized that if Ag^+^ or AgNPs indeed have different toxicity mechanisms, possibly due to AgNPs acting as a reservoir of Ag^+^, bacteria will evolve differently in their presence. We chose the environmental model bacterium *Pseudomonas putida* for the evolution experiment because its metabolism and stress responses have been well studied and its genome has been carefully annotated (Nelson *et al*., 2002; Puchałka *et al*., 2008; Belda *et al*., 2016). We pre-evolved *P. putida* KT2440 without Ag stress to disentangle adaptation to the new conditions of the evolution experiment with repeated cycles of growth and stationary phase from adaptation to Ag stress. In the following main evolution experiment, the pre-evolved culture further evolved in the presence of Ag^+^, well-defined AgNPs or Ag-free controls. The concentrations of Ag^+^ or AgNPs were chosen to be one tenth of the MICs to enable the cells to grow and evolve and to better reflect the low concentrations found in most environments. Nevertheless, they were higher than typical in natural environments and wastewater treatment plants (0–few hundred ng/L) (Li *et al*., 2016; Peters *et al*., 2018) but were within the range (1–100 μg/L) used in AgNP applications (Sankar *et al*., 2013; Chen *et al*., 2019). Whole genome sequencing allowed us to identify genetic changes during both evolution experiments. We found different mutation patterns in the populations that evolved under Ag^+^ or AgNP stress suggesting that AgNPs do have nanoparticle-specific effects.

## Results

The experimental design was to pre-evolve *P. putida* without stress to disentangle adaptation to the conditions of the evolution experiment from adaptation to silver stress, followed by evolution under Ag^+^ or AgNP stress and a stress-free control (Fig. 1).

**Figure 1.**
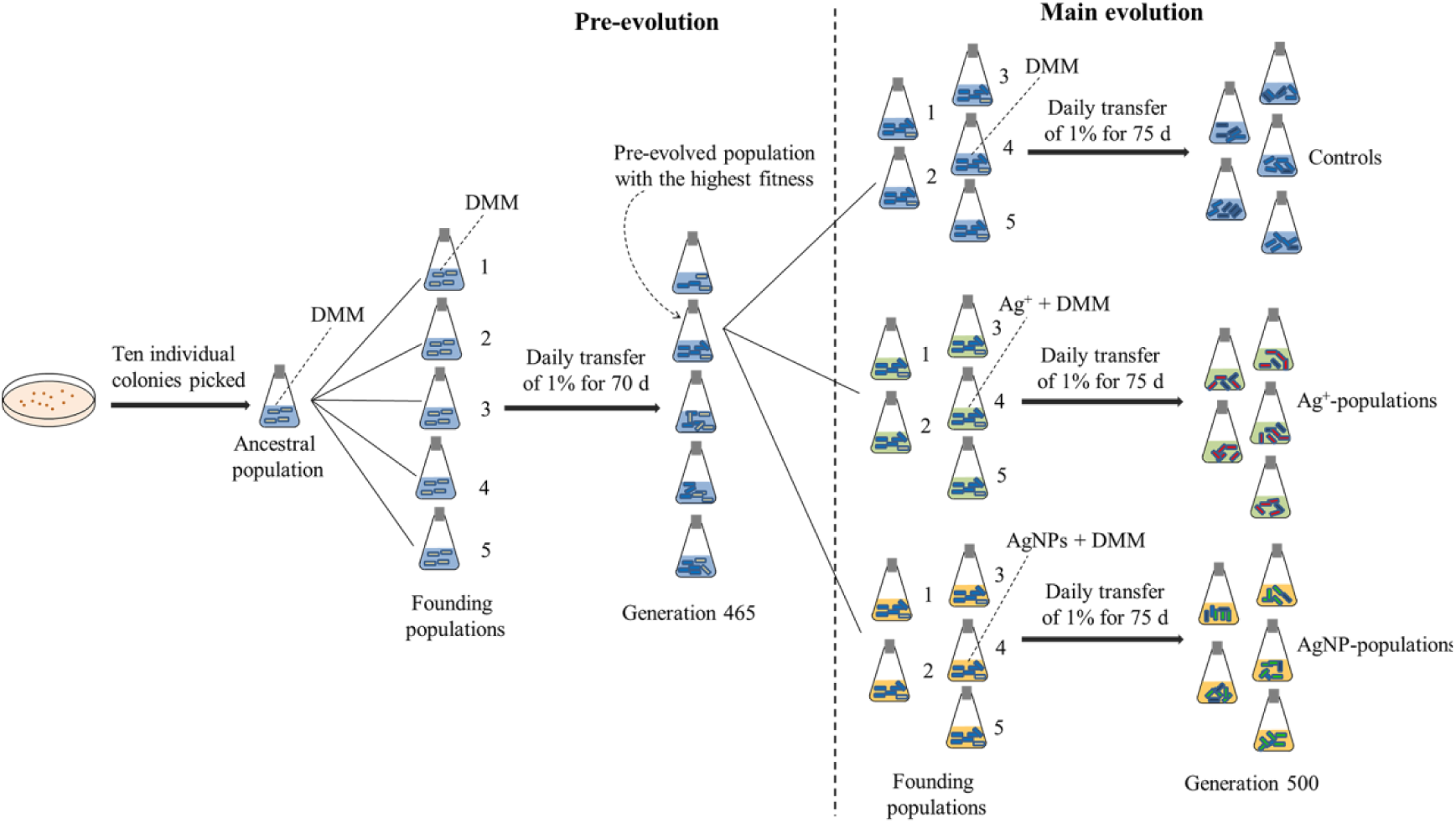
Design of the evolution experiment. In the pre-evolution stage, ten individual *Pseudomonas putida* KT2440 colonies were picked from LB plates and grown in liquid DMM into stationary phase to provide the ancestral population. Five parallel populations seeded from the ancestral population evolved independently for 70 d in DMM. In the main evolution experiment, the fittest population (# 2) amongst the five pre-evolved populations was divided into 15 populations that evolved 75 d either in the presence of Ag^+^ (5 μg/L), AgNPs (20 μg/L) or in Ag-free conditions. Five replicate cultures were set up for each condition.

### Synthesis and properties of AgNPs

We synthesized uncoated AgNPs to avoid confounding effects of the typically used capping agents such as citrate (El Badawy *et al*., 2011) and polyvinylpyrrolidone (Padmos *et al*., 2015). They were spherical and monodispersed (Fig. 2a). Their average diameter was 12.4 ± 4 nm and more than 95% were between 5 and 25 nm based on the transmission electron microscopy (TEM) images (Fig. 2b). A similar size distribution measured by differential centrifugal sedimentation (DCS) also confirmed the successful synthesis of uncoated AgNPs. Absence of detection of large particles by dynamic light scattering (DLS) indicated that the AgNPs had not aggregated. The AgNPs had a zeta potential of -23.0 ± 3.9 mV at a pH of 7.7 ± 0.6 in H_2_O. The pH of the AgNP stock suspension was 7.2 ± 0.4.

**Figure 2.**
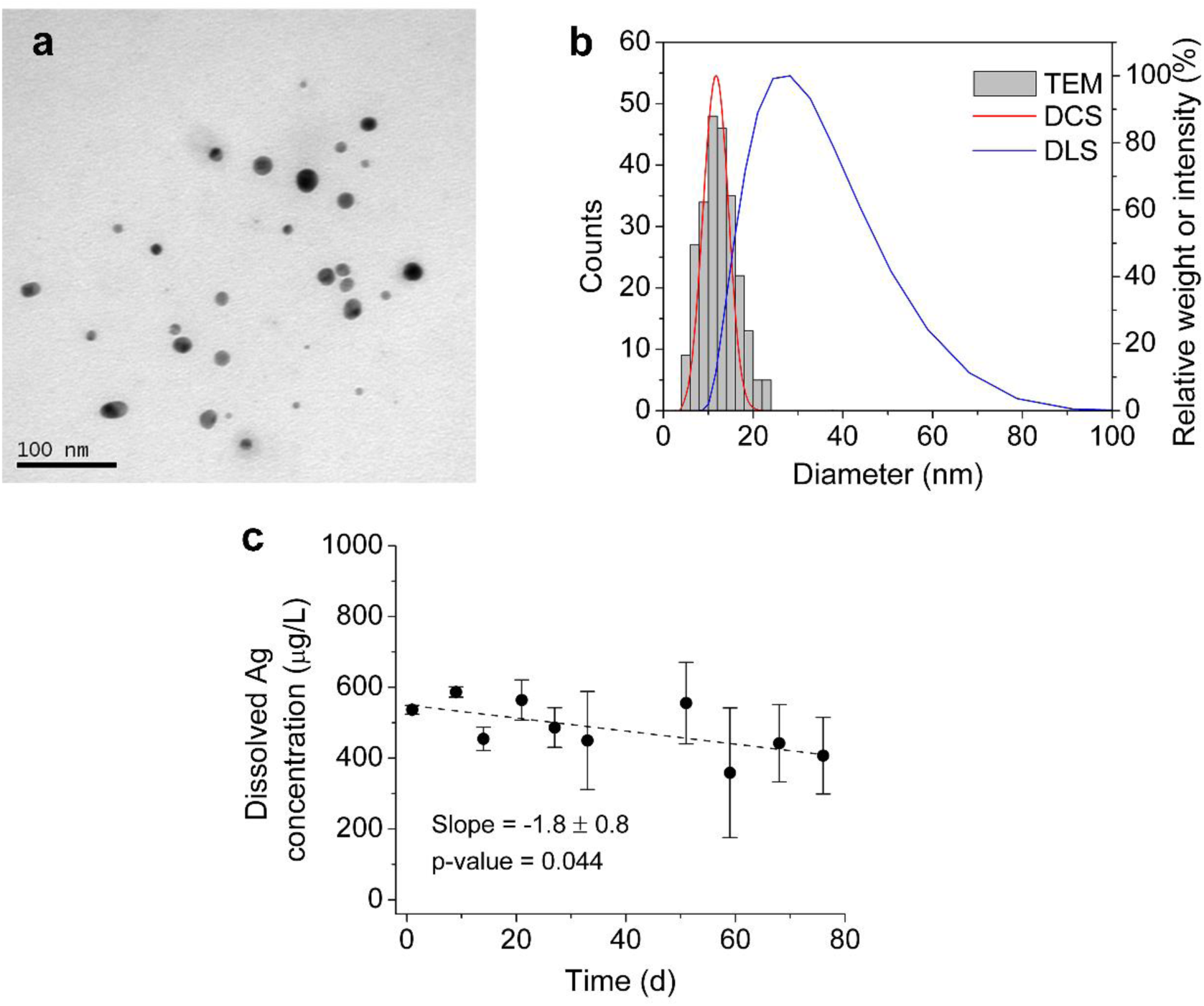
Properties of AgNPs. **(a)** Morphology obtained by TEM. **(b)** Diameter distribution based on TEM images analyzed by ImageJ (Schneider *et al*., 2012) and measured by DLS or DCS. **(c)** Dissolved Ag concentration in the AgNP stock over time during the main evolution experiment.

The stability of the AgNPs during storage was also examined. The concentration of total and dissolved Ag in the washed AgNP stock was 3,228 ± 469 and 537 ± 13 μg/L, respectively. The concentration of dissolved Ag in the AgNP stock was approximately constant over 76 d (Fig. 2c). The slight loss could be due to adsorption of Ag to the glass surface (Sekine *et al*., 2015; Malysheva *et al*., 2016). The stability was also confirmed by UV-Vis absorption spectra, which did not change over 44 d (Fig. S1a). Aggregation of AgNPs in the stock was not apparent in the Davis minimal medium (DMM) growth medium (Fig. S1b). Aggregation would be less likely at the lower concentration of AgNPs (161-fold dilution) that was used for the main evolution experiment since dilution of AgNPs substantially reduces aggregation (Dong *et al*., 2016). Thus, all data demonstrate stability of the AgNPs over the period of the evolution experiment.

### Mutations in the evolved populations

Contamination of cultures was not detected by sequencing as the GC content distribution of the contigs was the same and more than 99.9% of the reads mapped to Pseudomonadaceae family. To identify the mutations in the pre-evolved populations, the quality filtered reads from the ancestral and pre-evolved populations were aligned to the *Pseudomonas putida* KT2440 reference genome that has been well characterized (Supplementary Data S1) (Nelson *et al*., 2002; Belda *et al*., 2016). Twenty-four SNPs in total were identified in the pre-evolved population after 465 generations. Seven SNPs were located in intergenic regions, 13 SNPs were located in a putative surface adhesion protein gene (PP_0168), and 4 other SNPs were located in other coding regions (CDS) (*gacS, felQ, flgK* and PP_4920).

Mutations in all 15 populations of the main evolution experiment after 500 generations were identified (Supplementary Data S2). There were 198, 194 and 218 mutations in the populations that evolved without Ag stress, with Ag^+^ or AgNP stress, respectively. Most of these mutations did not occur in all the parallel cultures that had evolved under the same conditions. Half of the mutations were only found in one population. Populations from the three selective conditions had similar distributions of the number of parallel mutations (Fig. S2a). Multiple mutations in genes PP_0168 and PP_5662 were found in all populations, accounting for 81–93% of the mutations in CDS. These genes encode a putative surface adhesion protein and an apparent pseudogene, respectively. There was no significant difference in the total number of mutations or number of nonsynonymous mutations associated with the treatments (Fig. S2b,c).

### Mutations that differ between Ag^+^ and AgNP evolved populations

All mutations were listed and compared to determine whether they were associated with different Ag stresses (Supplementary Data S2). Figure 3a shows that the populations evolved under the three conditions shared approximately half of the mutations at the same positions, some of which were nonsynonymous (Fig. 3b). Despite most nonsynonymous mutations being unique to one of the 15 populations, many mutations appeared repeatedly in parallel populations (Supplementary Data S3). A modified Fisher’s exact test was used to determine the statistical significance of the association between mutation and condition (Table S2). Nine mutations were identified as statistically significant with p-values ≤ 0.05, including two mutations in intergenic regions, one mutation in one of the 16S ribosomal RNA genes, and six mutations in four protein coding sequences (*ftsZ, gacS, uvrY* and PP_2758) (Fig. 3c).

**Figure 3.**
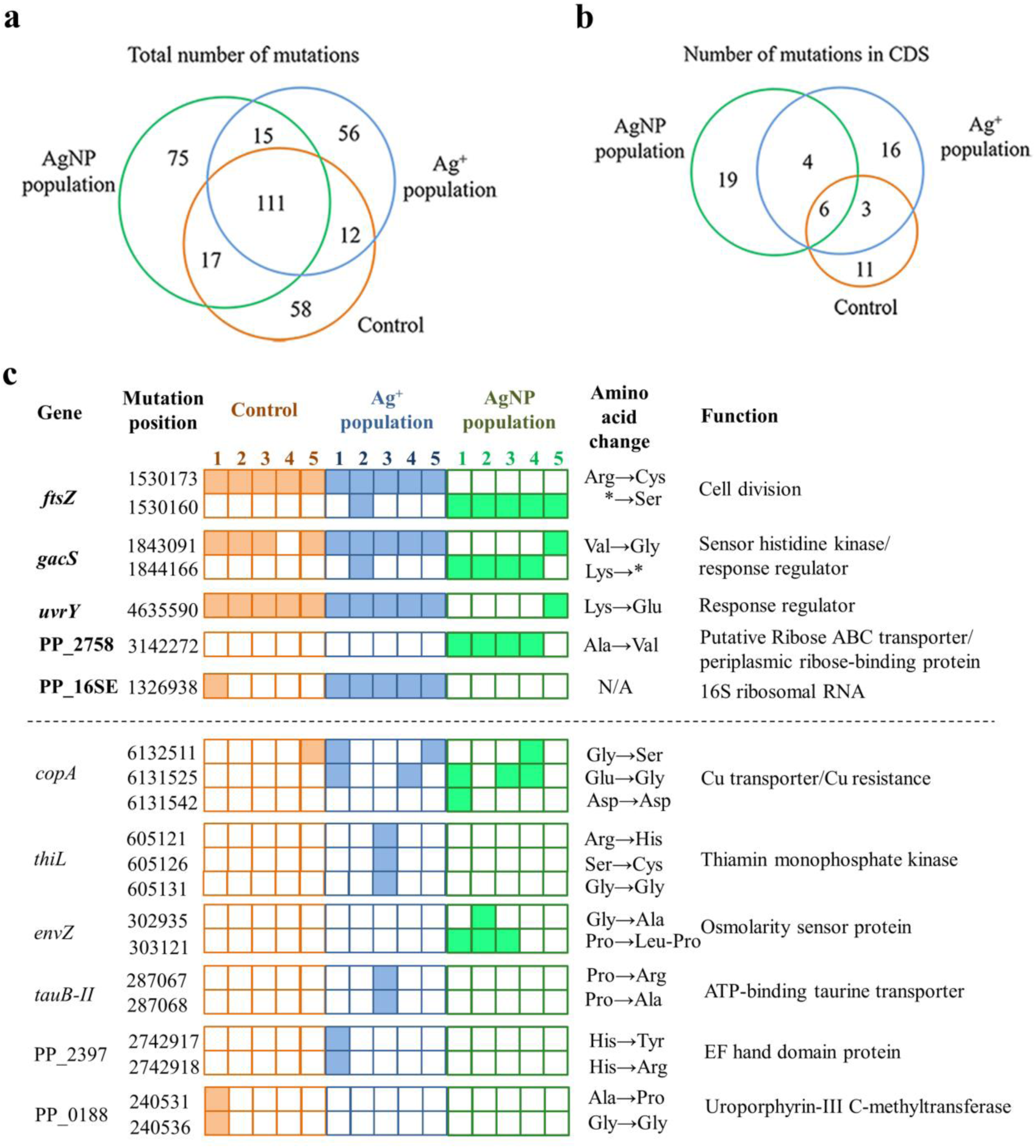
Mutations in the populations of the main evolution experiment. Venn diagram of counts of all mutations **(a)** and nonsynonymous mutations **(b)**. The numbers in the green, blue and orange circles refer to the counts of mutations in the AgNP-populations, Ag^+^-populations and controls, respectively. **(c)** List of genes with nonsynonymous mutations with mid-p-values ≤ 0.05 in Fisher’s exact test (gene names in bold and at the top) and genes with mutations in multiple positions (below). Five parallel populations had each evolved under three different conditions indicated by colors. The solid squares indicate mutation and the open squares indicate no mutation. For amino acid changes, “*” refers to a stop codon.

In addition, multiple mutations in the same genes were found in several genes in the populations treated with Ag^+^ or AgNPs but were mostly absent in the control populations (Fig. 3c): CopA is a P-type ATPase Cu-transporter associated with Cu/Ag resistance (Rensing *et al*., 2000; Stoyanov *et al*., 2003) so the mutations in three positions of *copA* in Ag^+^- and AgNP-populations might improve an Ag stress response. The occurrence of three mutations in *thiL* was only observed in one Ag^+^-population. The occurrence of two mutations in *tauB-II* and PP_2397 in Ag^+^-populations and two mutations in *envZ* in AgNP-populations further suggests that AgNPs and Ag^+^ had different toxicity mechanisms selecting for different mutations. Thus, Ag^+^ and AgNPs selected for different sets of mutations.

### Short-term fitness of evolved populations

Fitness is a phenotypic property that is dependent on the environment and as fitness differences drive evolution they can be used to quantify bacterial evolution (Orr, 2009). Here, relative fitness was measured over a single growth cycle by competing evolved populations against a reference *P. putida* KT2440 strain tagged with GFP to distinguish it from the evolved cells (Fig. S3). Since we observed large variations in fitness values in our preliminary competition assays, we examined the reproducibility of this assay (Fig. S4). The range in fitness values of 18 replicates was 1.38–1.64 and the coefficient of variation (standard deviation/mean) was 4.6%. This experiment with 18 replicates was repeated with similar results (Fig. S4). This shows that relative fitnesses are accurate (no systematic bias) but not precise (high variance) relative to the small changes in fitness values observed. Therefore, they need to be interpreted with caution.

The fitness of the GFP free KT2440 ancestor was 1.01 ± 0.03 relative to the GFP reference strain (Fig. 4a), indicating that the GFP tag did not affect fitness. After propagating the five parallel cultures in the pre-evolution experiment for 70 d (∼465 generations), their fitnesses ranged from 1.04 ± 0.02 to 1.16 ± 0.02 with a mean fitness of 1.10 ± 0.05 (Fig. 4a). The relative SD of fitness of 4.5% between the five parallel populations was similar to the value reported by Wiser and Lenski (Wiser and Lenski, 2015).

**Figure 4.**
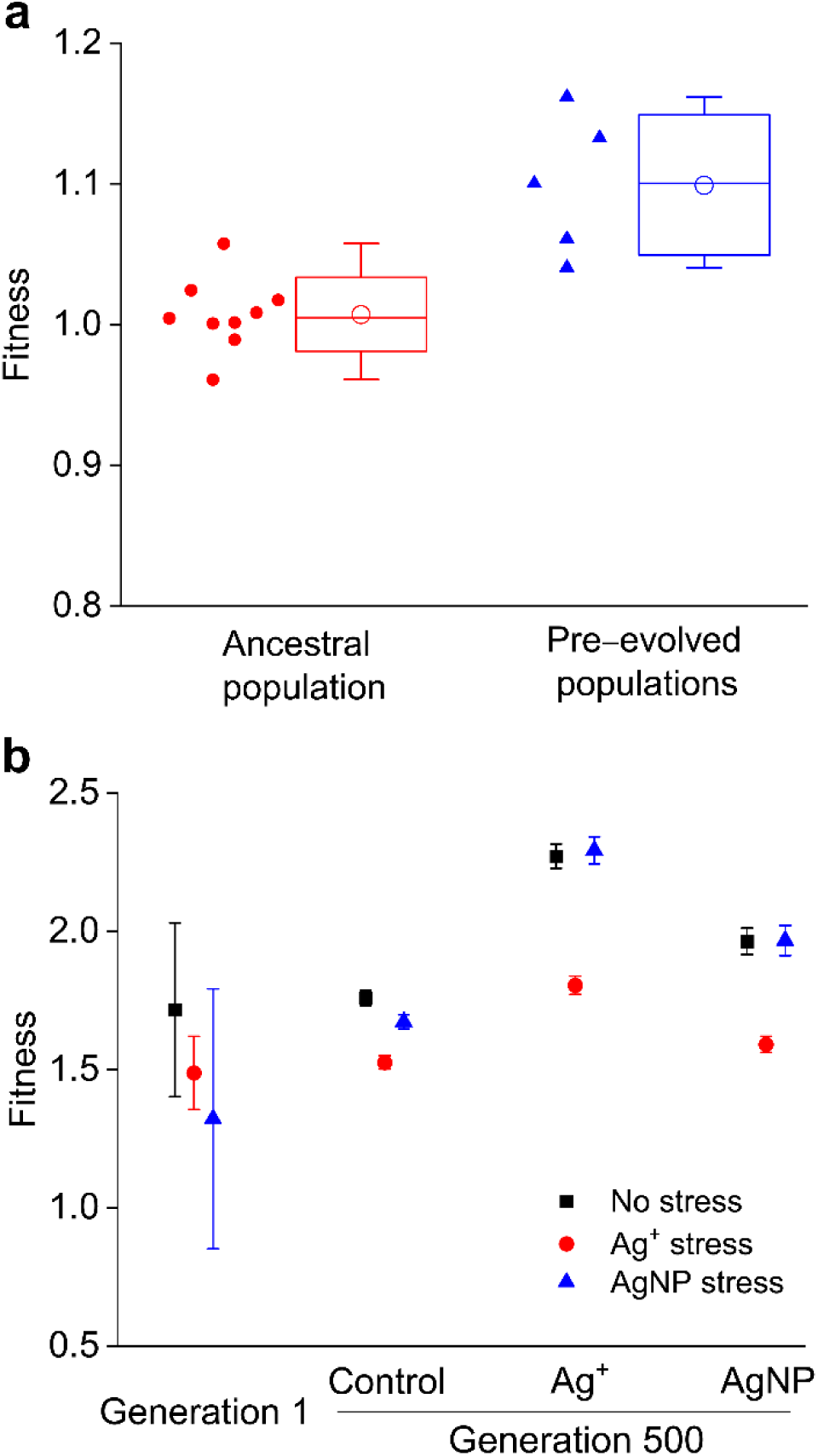
Fitnesses of evolved populations. **(a)** Fitnesses of the ancestral and pre-evolved populations. The data points for the ancestral culture are replicate assays of the same ancestral population competing against the GFP tagged reference strain, showing that the GFP tag had no significant effect (mean fitness ± SD, 1.01 ± 0.03) and also indicating variability of the assay (see also Fig. S4). The data points for the pre-evolved cultures are fitness values of the five parallel populations after 465 generations of pre-evolution. Boxes are SDs, bars medians, open circles means, and whiskers 5–95 percentiles. **(b)** Fitnesses of populations *evolved* in the presence of Ag^+^, AgNPs or Ag-free control medium and *competing* in the presence of Ag^+^, AgNP or Ag-free control medium. The five parallel cultures were pooled for this assay. Error bars for generation 1 represent the SDs of three replicate measurements. Error bars for generation 500 represent SDs derived from the viable counts assuming that the sampling of cells for the counts was the only source of variation and that cells were randomly sampled and counts therefore Poisson distributed (Clegg et al. unpublished).

Population 2 of the pre-evolution experiment with the largest fitness improvement was chosen as the founding culture for the main evolution experiment. The MICs for Ag^+^ and AgNPs for this founding culture were 50 and 200 μg/L by the standard broth dilution method, respectively. We chose 10% of these values, 5 μg/L Ag^+^ or 20 μg/L AgNPs, to select for adaptation to sub-lethal concentrations of silver in the main evolution experiment as these concentrations were not too high for the bacteria to grow and evolve and considering that concentrations in the environment are typically much lower than the MICs.

The concentration of dissolved Ag^+^ in the AgNP stock changed only slightly during the main evolution experiment (Fig. 2c). The concentration of dissolved Ag^+^ added to the cultures with the 20 μg/L AgNP dosage was calculated to be between 2.2–3.6 μg/L. Moreover, silver nanoparticles will release Ag^+^ upon addition to the bacterial culture at a rate of up to 0.03 μg Ag^+^/(L·h), calculated from the initial dissolution rate of AgNPs in DMM of (1.9 ± 0.4) × 10^−3^ μg Ag^+^/(μg AgNP·h), corresponding to 0.72 μg Ag^+^/L released in 24 h (Dong *et al*., 2017). This means that treatment by AgNPs was inevitably a combined Ag^+^ and AgNP stress, as in all other studies.

The five parallel populations were pooled to determine their fitness at the start and end of the main evolution experiment (generation 1 and 500) (Fig. 4b). Given that absolute differences in relative fitness values below about 0.2 may be in error, even if error bars happen to be smaller, one can only conclude that results are not in disagreement with general expectations: (i) Evolved populations had increased fitness under the conditions they evolved in or at least not decreased fitness in the case of the control. (ii) Fitness competing under Ag^+^ stress was lower than under AgNP stress. (iii) Fitness differences between AgNP stress and no stress were negligible at least for populations evolved in either Ag^+^ or AgNP. (iv) Increased fitness under Ag^+^ or AgNP stress did not compromise fitness in the absence of Ag (Fig. 4b).

We also measured the MICs of populations evolved under each treatment. The Ag^+^ MICs were in the same range of 5-7.5 μg/L for Ag-free, Ag^+^ and AgNP evolved populations (Fig. S5a), indicating that there was no clear difference between treatments. Likewise, the measured AgNP MICs of different cultures overlapped, with most replicates in the range for the ancestral population (160-240 μg/L) and one a little higher, suggesting again that there was no clear change of the MIC during evolution (Fig. S5b). It is to be noted that the Ag^+^ MICs measured by the agar dilution method were ∼5-fold lower than those determined by the broth dilution method. This discrepancy may be due to the different assays. In the broth assay, the population MIC is largely determined by the growth of cells with the highest MIC, which can proliferate more quickly and gradually increase in frequency in the population in silver-containing media. In contrast, the MIC obtained on agar plates largely depends on the fitness of the majority as each colony arises from a single cell of the original culture. Any rare cells with high MIC are likely to be missed as plating out takes a sample of the population. Finally, it is important to realize that MICs provide little information on the effects of sub-lethal concentrations that were used in the evolution experiments, so they do not mimic the evolution experiment as well as the short-term fitness assays, which correspond to a single growth cycle of the evolution experiment yet lack the repeated change from stationary phase through lag phase to next exponential phase.

## Discussion

We employed an evolution experiment to ask whether bacteria respond differently to Ag^+^ or AgNP stress. If Ag^+^ selects for different mutations than AgNPs, this would imply different toxicity mechanisms of ionic and nanoparticulate Ag. Consistent environmental conditions were kept throughout the whole evolution experiment, as the initially synthesized AgNPs were stably maintained. Also, reactants left after synthesis were removed to avoid any confounding effects on bacterial evolution. Overall, selective conditions in the Ag^+^ or AgNP treatment hardly changed from passage to passage, but the Ag^+^ or AgNP stress likely changed during each growth cycle as a result of the dissolution of AgNPs (less than 5% of the initially dosed AgNPs dissolved within one growth cycle according to calculations based on rates measured (Dong *et al*., 2017).

To our knowledge, this is the first study to have carried out a pre-evolution experiment to adapt the ancestor to the novel selective regime without the specific stress of interest in order to at least partially disentangle selection for confounding conditions from selection for conditions of interest. Indeed, the ancestor gained fitness by accumulating mutations in the pre-evolution experiment (Fig. 4 and Supplementary Data S1). As expected from the long-term evolution experiment in *Escherichia coli* showing that evolution never ceases (Wiser *et al*., 2013), the populations that continued to evolve under the same Ag/AgNP free conditions acquired further mutations. These mutations presumably increased fitness as they were selected for; however, fitness measurement errors were too large to confirm this. A thorough analysis of the increase of fitness values in the *Escherichia coli* long-term evolution experiment had similar precision (Wiser and Lenski, 2015). Such imprecision increases when the evolved populations become fitter, due to the increasing difficulty of enumerating the less-fit population and thereby magnifying measurement errors (Wiser and Lenski, 2015). In the pre-evolution experiment, the fitness of *P. putida* populations increased to 1.10 ± 0.05 after 465 generations, close to the calculated value of 1.12 for the long-term evolution experiment (LTEE) of *Escherichia coli* by Lenski and co-workers (Wiser *et al*., 2013; Lenski *et al*., 2015). The observed frequency of two nonsynonymous SNPs in our pre-evolution experiment were also similar to the LTEE (Barrick *et al*., 2009). Unexpectedly, some mutations reverted (Supplementary Data S1). Hence, we could not fully disentangle selection for the growth regime from selection for Ag/AgNP stress, as we could not continue the pre-evolution experiment indefinitely. Instead, we chose to run it for a similar number of generations as the main evolution experiment.

In the main evolution experiment, we chose a low dosage of Ag^+^ and AgNPs for two reasons. First, to be closer to the low concentrations found in most environments, see above. Second, to make sure that the bacteria were able to grow and evolve, avoiding the population crashes observed previously (Graves *et al*., 2015). Lower stress tends to enable bacteria to explore diverse mutation paths rather than the fewer and shorter paths leading to larger fitness gains, if they exist, under strong selection (Gullberg *et al*., 2011; Barrick and Lenski, 2013). The coexistence of Ag^+^ and AgNPs in the AgNP treatment complicates the comparison of Ag^+^ and AgNP treatments. This is unavoidable under aerobic growth conditions and indeed useful for elucidating nanoparticle-specific effects. Since the AgNP-populations had the opportunity to adapt to both Ag^+^ and AgNPs, one would expect these populations to have gained a higher fitness under Ag^+^ stress than the control populations. The results are consistent with this expectation (Fig. 4b), but the measurement errors do not allow firm conclusions. Further, AgNP-populations should have a higher fitness under AgNP stress than the Ag^+^ and control populations. However, the Ag^+^-populations had a (potentially significant) higher fitness under AgNP stress than the AgNP-populations, though the control populations had the lowest fitness. Overall, the fitness results alone have not answered the question whether AgNPs have specific effects on cells.

Unlike studies that used high or gradually increasing concentrations of an antimicrobial during evolution and obtained considerable increases of MICs (Gunawan *et al*., 2013; Panáček *et al*., 2018), we applied much lower levels of Ag^+^ or AgNPs that were more environmentally relevant to elucidate the microbial adaption under realistic conditions. It has been shown that antimicrobials are selecting for resistance in a ‘selective window’ below the MIC of the sensitive strain where the resistant mutant is still fitter than the sensitive wild type (Gullberg *et al*., 2011). Although the MICs of the evolved populations did not appear to have increased, and indeed did not have to increase given that changes in phenotype at sub-lethal concentrations may not change the MICs, subtler adaptations have to be presumed to have occurred. More obvious changes of phenotype may require prolonged accumulation of beneficial mutations (Blount *et al*., 2018).

While phenotypic changes were not obvious, the significant differences in the genetic profiles of Ag^+^ and AgNP evolved populations identified by whole-genome sequencing nevertheless answers the question whether bacteria evolved differently in response to ionic or nanoparticulate Ag stress in the affirmative. Notably, some mutations unique to one condition occurred in most (4-5) of the five parallel populations. Many of the mutations found in parallel populations have not previously been reported to be associated with the antibacterial actions of Ag^+^ or AgNPs. Understanding of the molecular mechanisms behind each mutation was out of scope for this work and requires detailed further investigation.

The populations that evolved under Ag^+^ or AgNP treatment shared many similar mutations with the control populations, but some mutations were specific for the selective conditions (Fig. 3a,b). The treatment with Ag^+^ had a significant association with the mutation of gene PP_16SE (Fig. 3c). In contrast, mutations in five genes (*ftsZ, gacS, uvrY*, PP_2758 and PP_16SE) were significantly associated with AgNP treatment. This suggests that AgNPs affected the cells differently from Ag^+^ in a way that different mutations were positively selected in response.

Moreover, multiple mutations in the same genes (*copA, thiL, tauB-II* and PP_2397) in the Ag^+^-populations that were absent in the control populations demonstrated that Ag^+^ did select for specific genetic changes (Fig. 3c). The AgNP selected populations were expected to have all the same mutations that appeared in the Ag^+^ selected populations since the AgNP suspension contained Ag^+^, but the genes with multiple mutations in Ag^+^ and AgNP selected populations were different, i.e. the mutations in PP_2758 and *envZ* were only observed in the AgNP-populations and the mutations in *thiL, tauB-II* and PP_2397 were only found in the Ag^+^-populations (Fig. 3c), further suggesting that AgNPs exerted different selective pressures. A recent study that evolved two strains of *E. coli* and one strain of *P. aeruginosa* under AgNP stress found that all three strains evolved AgNP-specific resistance due to flagellin facilitated aggregation and sedimentation of AgNPs (Panáček *et al*., 2018). This kind of resistance should be a public good as it would benefit other non-resistant bacteria nearby. Flagella-less mutants or immotile control bacteria were not included in this study. Curiously, the AgNP resistant phenotypes were stable in all strains although mutations in coding sequences were not detected (Panáček *et al*., 2018).

Previous studies have demonstrated that the CusCFBA efflux system (Randall *et al*., 2015), the plasmid-related SilCFBA system (Gupta *et al*., 1999), the outer membrane porins OmpF and OmpC (Li *et al*., 1997; Randall *et al*., 2015) and periplasmic proteins (SilE and an engineered Ag-binding protein) (Sedlak *et al*., 2012; Asiani *et al*., 2016) participate in bacterial detoxification of Ag^+^. A mutation in *cus* was not detected in any evolved populations. Instead, *copA*, which has a role in Ag efflux (Stoyanov *et al*., 2003), was mutated in multiple positions in Ag^+^ and AgNP stressed populations, providing direct evidence of adaptation. A previous study identified 68 open reading frames possibly contributing to metal homeostasis or resistance in *P. putida*, including two systems for monovalent cations (*pacS, cusCBA*), a cryptic *silP* gene, two operons for Cu chelation (*copAB*) and one metallothionein (Canovas *et al*., 2003). Surprisingly, there were no mutations in these genes apart from *copA* in the Ag^+^ or AgNP-evolved populations. The other mutations found in the evolved populations have not been previously reported to be linked with Ag resistance, suggesting that novel mechanisms are involved in the bacterial response to Ag stress.

Faster growth rate and larger cell size are phenotypic changes of evolved *E. coli* cells in the long-term evolution experiment (Lenski, 2010). FtsZ is recruited to the cell membrane to form the Z-ring that is required for cell division (Weart and Levin, 2003). All five parallel AgNP-populations acquired a mutation in *ftsZ* turning the stop codon into a codon for Ser, while in all five Ag^+^-populations, Arg was substituted with Cys. A similar phenomenon was observed in *gacS*, which encodes a transmembrane signal sensor protein that is coupled with the response regulator GacA, comprising a two-component system (Heeb and Haas, 2001) that can influence biofilm formation (López-Sánchez *et al*., 2016). A mutation of *uvrY*, which is homologous to *gacA*, occurred in the Ag^+^-populations and the control populations but was present in only one of the AgNP-populations, implying different regulatory strategies in response to the different Ag species.

The two AgNP-specific mutations in *envZ* and in PP_2758 (encoding a putative periplasmic protein) might indicate that AgNPs specifically affect the bacterial cell wall. The mutation in *envZ*, the inner membrane sensor of the envZ/ompR two-component regulatory system, may prevent the expression of OmpF/C porins (Slauch *et al*., 1988), which may reduce the internalization of Ag into the evolved bacterial cells or ROS production. The role of OmpF/C in bacterial Ag-resistance is well documented (Li *et al*., 1997; Randall *et al*., 2015). The genes *thiL* and PP_2397, mutated in Ag^+^-populations, encode a thiamin monophosphate kinase and an EF-hand domain protein, which are located in the cytosol and have magnesium (NISHINO, 1972) and calcium binding domains (Lewit-Bentley and Réty, 2000), respectively. The effect of the mutation in *tauB*-*II*, which codes for the taurine ABC transporter ATP-binding subunit (TauB) that is located at the cytoplasmic side (Eichhorn *et al*., 2000), is also unclear. Presumably, because AgNPs cannot penetrate the bacterial cell wall, the mutations specifically selected by AgNPs were associated with proteins in the periplasm and with proteins that regulate outer membrane proteins. In contrast, Ag^+^ mostly selected for mutations in cytoplasmic proteins, indicating a response to intracellular Ag^+^. Overall, *P. putida* evolved partially different adaptations upon exposure to low doses of Ag^+^ versus AgNPs.

In conclusion, *P. putida* populations evolved differently under prolonged exposure to low concentrations of Ag^+^ versus AgNPs. Mutations unique to AgNP-populations were mostly associated with the cell surface and two-component systems. In contrast, the mutations specific to Ag^+^-populations were in uptake systems and metal-binding proteins in the cytosol. The significantly different mutation patterns in populations that evolved under Ag^+^ or AgNP stress suggest different toxicity pathways for ionic and nanoparticulate Ag in bacteria. Since it has previously been demonstrated, at least in *E. coli*, that AgNPs are not toxic under anoxic conditions, which prevent the release of Ag^+^ ions (Xiu *et al*., 2012), we can rule out any *direct effects* and have to consider a more subtle mechanism for the AgNP effects to explain our results. The differences between Ag^+^ and AgNPs must be attributable to *nanoparticle-specific but indirect* effects, especially on the cell surface. These AgNP-specific effects are presumably due to differences in Ag^+^ delivery: while a Ag^+^ treatment will start with the highest concentration at the beginning, which is then titrated away, the AgNP treatment is accompanied by the binding of AgNPs to cell surfaces and continuously delivering Ag^+^, replenishing any Ag^+^ that has been titrated away (Dong *et al*., 2017). The existence of AgNP-specific effects has important implications for the application of AgNPs, toxicity testing, mitigation of environmental impacts and regulation of AgNPs.

## Experimental procedures

### Preparation of uncoated AgNPs

Uncoated AgNPs were made to avoid effects due to coating materials. They were produced by reducing AgNO_3_ with NaBH_4_ using an aqueous synthesis method (Dong *et al*., 2016). In order to remove any dissolved Ag and reactants remaining after the synthesis, the AgNP suspension was washed with deionized (DI) water by ultrafiltration through a membrane with pore size of 3 kDa (Amicon Ultracel 3 KDa Ultrafiltration Discs, Millipore, Billerica, USA) under nitrogen gas pressure (1 bar) using a stirrer cell (400 mL, Amicon, Millipore, Billerica, USA). After 10 washing cycles, any dissolved chemicals in the AgNP suspension would have been diluted at least 1,000 fold (2^10^). Any remaining trace amounts of B(OH)_4_^-^ (<1.5 μM), Na^+^ (<1.5 μM) and NO_3_^-^ (<0.06 μM) were assumed not to interfere with bacterial evolution. Some Ag^+^ unavoidably remained in the suspension due to the ongoing dissolution of AgNPs. The washed AgNP suspension was sterilized by filtration through a membrane with pore size of 0.2 μm and stored at nitrogen gas atmosphere.

### Characterization of uncoated AgNPs

The size distribution of AgNPs was measured by DLS (Zetasizer Nano, Malvern Instruments, Malvern, UK) and DCS (DC24000, CPS Instruments Europe, Oosterhout, Netherlands) (Krpetić *et al*., 2013). Localized surface plasmon resonance of uncoated AgNPs in suspension was measured by UV-Vis absorption spectrometry (UV-Vis 6800, Jenway, Stone, UK) (Peng *et al*., 2010). Zeta potential was also measured by the Zetasizer Nano (El Badawy *et al*., 2010). The morphology of AgNPs was examined by TEM (JEOL 1200EX, Tokyo, Japan).

To examine the aggregation state of AgNPs in the medium used for the evolution experiment, the AgNP stock (0.5 mL) was mixed with DMM (0.5 mL) in a cuvette using the pipette and the changes of z-average diameter were immediately recorded for 2 h (Zetasizer Nano). The concentration of dissolved Ag in the AgNP stock suspension during the evolution experiment was measured using a combined aggregation-centrifugation method (Dong *et al*., 2016), but a higher Ca(NO_3_)_2_ final concentration of 10 mM was needed to quickly aggregate washed AgNPs (Fig. S6).

### Bacterial strains and media

The strain for the evolution experiment was a wild type *Pseudomonas putida* KT2440 that was kindly donated by Víctor de Lorenzo (Centro Nacional de Biotecnología, CNB-CSIC, Madrid, Spain). A *Pseudomonas putida* KT2440 chromosomally tagged with GFP was used as the reference ancestor in competition assays and also kindly donated by Víctor de Lorenzo. In order to minimize precipitation and reactions of AgNPs during the exposure, the simple salts medium DMM rather than a complex medium was used to grow the cells as in the *E. coli* long-term evolution experiment (Lenski *et al*., 1991) and previous toxicological studies with *P. putida* (Fabrega *et al*., 2009). DMM contains (per liter) (pH 7.2): 7.0 g K_2_HPO_4_, 2.0 g KH_2_PO_4_, 1.0 g (NH_4_)_2_SO_4_, 0.1 g MgSO_4_, 1.530 g sodium citrate dihydrate and 1 mL SL10 trace elements solution (Atlas, 2010). The trace elements stock solution contains (per liter): 1500 mg FeCl_2_·4H_2_O, 190 mg CoCl_2_·6H_2_O, 100 mg MnCl_2_·4H_2_O, 70 mg ZnCl_2_, 6 mg H_3_BO_3_, 36 mg Na_2_MoO_4_·2H_2_O, 24 mg NiCl_2_·6H_2_O and 2 mg CuCl_2_·2H_2_O (Widdel *et al*., 1983). Precipitation of silver (20 μg/L) in the DMM salts solution can be neglected according to solubility calculations carried out with Visual MINTEQ (Version 3.0, The Royal Institute of Technology, Sweden) (Table S1).

### Evolution experiment

Ten individual *P. putida* colonies on LB plates were picked and grown in 20 mL DMM overnight. This culture was designated as the ancestral population and used to seed five parallel cultures that were transferred daily for 70 d. Every 24 h, 0.1 mL of the evolving cultures in stationary phase were transferred into 9.9 mL fresh DMM in 100 mL glass flasks and grown at 30 °C. Cultures were aerated by shaking at 150 rpm in a water bath shaker. The 100-fold dilution resulted in an initial cell density of ∼10^7^ CFU/mL and 6.64 (log_2_100) generations each day. Preserving the evolved populations in frozen condition (−80 °C) every 3 d at the early stage (1-27 d) and every 7 d later based on a previously reported method allowed us to store the ‘fossil record’ of their evolution.

Before the main evolution experiment, we assessed the MIC of the ancestral population following the standard broth dilution method (Andrews, 2001). The population with the highest fitness, assessed by short-term competition assays, amongst the five parallel cultures after ∼467 generations in the pre-evolution experiment was chosen as the founding population for the main evolution experiment. The frozen pre-evolved population was revived in DMM overnight. Five parallels each were set up by transferring 0.1 mL aliquots into 9.9 mL fresh DMM containing either Ag^+^ or AgNPs (5 μg/L Ag^+^, 20 μg/L AgNPs) at a concentration of one tenth of the respective MICs to start the main evolution. They were designated as the Ag^+^-population and AgNP-population, respectively. Another five parallel cultures evolved in DMM devoid of Ag^+^ and AgNPs, designated as the control. These 15 populations were propagated independently for 75 d (500 generations) in the same way as in the pre-evolution experiment, apart from the Ag^+^ or AgNP additions to the medium. To ensure consistency of the Ag^+^ or AgNP treatment during the main evolution experiment, the same AgNO_3_ or AgNP stock was used throughout.

### Short-term fitness assay

Fitness differences between evolved and ancestral populations, or relative fitness, was measured by competition in the same medium (Lenski *et al*., 1991). This is similar to the evolution experiment, but comprises only one growth cycle so is short-term and lacks the restart of growth in the presence or absence of silver after a passage. In this study, the evolved population was competed against a GFP tagged *P. putida* KT2440 reference strain. The constitutive expression of the chromosomally inserted GFP gene enabled us to distinguish evolved and reference strains on the same LB agar plates used for counting since LB is better for resuscitation and survival of the bacterial cells. The competitions were prepared by reviving frozen stock cultures of the two populations in DMM overnight. These were then mixed together in approximately equal numbers judged by OD_600_ and added into fresh DMM for growth (initial cell number 3-5 × 10^6^ CFU/mL). Viable cells were counted before and after the 24 h competition. The relative fitness *F* is defined as 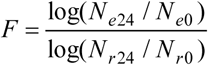 (equation 1), where *N*_*e0*_ and *N*_*e24*_ is the initial and final (24 h) cell density for the evolved population, respectively, and *N*_*r0*_ and *N*_*r24*_ is the initial and final (24 h) cell density for the reference population, respectively. That part of the standard error of this fitness measure that is solely due to sampling the cells was estimated by optimizing the fit of simulated sampling to the observed data with a program coded by our laboratory (Clegg et al. unpublished, code available on github: https://github.com/kreft/fitnessCalc/).

To assess the fitnesses of cultures from the main evolution experiment, the competitions were carried out as described above but in DMM containing either no silver, 5 μg/L Ag^+^ or 20 μg/L AgNPs. Due to the large number of combinations of three selection pressures and three assay conditions it was necessary to pool all five parallel cultures for the competition with the reference strain. After reviving the 15 frozen cultures in DMM overnight, the five populations from the same treatment were pooled with approximately same number of each population before competing against the reference culture.

To check the validity of the fitness measurement, the reproducibility of the fitness assay was examined by competing one evolved population (culture 1 from the control conditions of the main evolution experiment, generation 500) against the reference strain in six replicate competitions as described above. After 6 h of growth, when the two populations were expected to be at the end of the exponential phase, 1 mL of the competitor mix was collected, pelleted (centrifuged at 5,100 g for 10 min), resuspended in DMM salts without citrate or any other carbon and energy source, and stored at 4 °C. For the purpose of convenience, viable counting was carried out by spreading cells on LB plates the next day. The six replicate competitions were counted in triplicate to estimate the variance due to viable counting. Additionally, the whole procedure was repeated one week later.

### MIC assay for evolved populations

MICs by definition provide information on an antimicrobial concentration required for complete inhibition, but do not reveal effects of sub-lethal concentrations. We measured the MICs of evolved cultures on the level of individual cells giving rise to colonies on agar plates because reviving frozen cultures in liquid media may change the bacterial population structure as a result of potentially heterogeneous growth rates of subpopulations present in evolved cultures. In brief, the evolved frozen culture was thawed, diluted by DI H_2_O to a cell density of around 1.6 × 10^5^ CFU/mL and spread onto DMM agar that contained varying concentrations of Ag^+^ (0-12.5 µg/L) or AgNP (0-400 µg/L). To reduce any potential effect of changes of Ag^+^ or AgNP in the agar, Ag^+^ agar plates were used within three weeks and AgNP plates on the day of preparation. After inoculation, the bacterial cells were cultivated for 2 d at 30°C in an incubator. The MIC was determined as the concentration at which colony formation was inhibited (clear drop in colony number < 5).

### Whole genome sequencing

Reviving the frozen populations was performed by streaking the stock cultures on LB plates. This step was carried out to satisfy the standard operating procedure of the sequencing pipeline but may have introduced a bias against mutants that grow poorly on agar plates. Bacteria from these plates were harvested with a sterile loop and transferred into Microbank tubes (Microbank™, Pro-Lab Diagnostics, Canada) that contain beads binding the bacterial cells. For DNA extraction, three beads were washed with extraction buffer containing lysozyme and RNase A and incubated for 25 min at 37 °C. Proteinase K and RNaseA were then added and the mixture incubated for another 5 min at 65 °C. Genomic DNA was purified using SPRI beads and resuspended in an EB buffer.

DNA was quantified in triplicate with the Quantit dsDNA HS assay in an Eppendorff AF2200 plate reader. Genomic DNA libraries were prepared for Illumina sequencing using Nextera XT Library Prep Kit (Illumina, San Diego, USA) according to the manufacturer’s protocol with the following modifications: two nanograms of DNA instead of one were used as input, and PCR elongation time was increased to 1 min from 0.5 min. DNA quantification and library preparation were carried out on a Hamilton Microlab STAR automated liquid handling system. Pooled libraries were quantified using the Kapa Biosystems Library Quantification Kit on a Roche light cycler 96 qPCR machine. Libraries were sequenced by Illumina MiSeq using a 250 bp paired end protocol. The genome sequences of ancestral and evolved populations were deposited at the European Nucleotide Archive (ENA accession number: PRJEB16768). Stocks of all the sequenced cultures are maintained by MicrobesNG.

### Variant calling

The raw reads were trimmed with the tool Trimmomatic to remove adapters and bases with quality scores < 15. Reads were aligned to the *Pseudomonas putida* KT2440 reference genome (Accession number AE015451, GenBank) using Burrows-Wheeler Aligner (BWA-Mem 0.7.5) and processed using SAMtools 1.2. Variants were called using VarScan 2.3.9 with the following settings: Phred score ≥ 15, depth of variant-supporting bases ≥ 3, variant allele frequency ≥ 0.1, p-value for calling variants ≤ 0.05. Variants were further manually filtered according to strand bias (0.1 ≤ ratio of variant allele frequency of one strand over the other ≤ 0.9). Single nucleotide polymorphisms (SNPs) in homopolymer tracts of length ≥ 8 bp were then manually excluded. We annotated the mutations according to a revised *Pseudomonas putida* KT2440 genome (Belda *et al*., 2016) (Accession number AE015451.2, GenBank).

### Association of mutations with selective pressures and other statistics

Any association between mutations (wild type vs mutant) and selective pressures (Ag^+^, AgNP or control) was tested with a modified version of Fisher’s exact test that accounts for the discrete nature of the results by calculating mid-p-values (Choi *et al*., 2015). Such mid-p-values for the two-tailed test were calculated with the oddsratio.fisher function in R (part of the epitools package) (Table S2). OriginPro 8 Software was used to carry out statistical analysis of reproducibility of fitness assays and graphical visualization of data.

## Acknowledgements

We thank the Darwin Trust Edinburgh for Feng Dong’s PhD scholarship and the Mexican National Council for Science and Technology (CONACyT) for Ana Carrazco Quevedo’s PhD scholarship. We are indebted to Pablo Fuentes-Utrilla and Emily Richardson from MicrobesNG (https://www.microbesng.uk) at the Institute for Microbiology and Infection (IMI) at the University of Birmingham (funded by the BBSRC, grant number BB/L024209/1) for sequencing and bioinformatic analysis. We are grateful for the support from NERC via the Facility for Environmental Nanoscience Analysis and Characterisation (FENAC) at the University of Birmingham.

## Competing financial interests

The authors declare no competing financial interests.

## Notes

### Competing Interest Statement

The authors have declared no competing interest.

### Summary of Updates

author name changed

