## Supporting Information for "Experimental Evolution of *Pseudomonas putida* under Silver Ion versus Nanoparticle Stress"

**Table S1** Prediction of potential precipitation of $Ag^+$ (20 μg/L) in Davis minimal medium.

| Mineral | log(IAP) | Saturation index (log(IAP) – log($K_{sp}$)) |
|---|---|---|
| $Ag_2SO_{4(s)}$ | -16.824 | -12.053 |
| $Ag_3PO_{4(s)}$ | -28.315 | -10.725 |
| Cerargyrite ($AgCl_{(s)}$) | -11.931 | -2.370 |

IAP refers to ion activity product, $K_{sp}$ refers to solubility product constant. A saturation index < 0 indicates that $Ag^+$ precipitation is not expected to occur. Here, the index is well below zero so precipitation is very unlikely, even if activity and solubility parameters were somewhat different in our experimental conditions.

**Table S2** Mid-p-values calculated with a modified Fisher's exact test for association between mutation and condition.

| Outcome: WT, Mutant | | Condition 2 | | | | | |
|---|---|---|---|---|---|---|---|
| | | 0, 5 | 1, 4 | 2, 3 | 3, 2 | 4, 1 | 5, 0 |
| Condition 1 | 5, 0 | 0.0040 | 0.0238 | 0.0833 | 0.2222 | 0.5 | 1 |
| | 4, 1 | 0.0238 | 0.1071 | 0.2857 | 0.5833 | 1 | 0.5 |
| | 3, 2 | 0.0833 | 0.2857 | 0.6032 | 1 | 0.5833 | 0.2222 |
| | 2, 3 | 0.2222 | 0.5833 | 1 | 0.6032 | 0.2857 | 0.0833 |
| | 1, 4 | 0.5 | 1 | 0.5833 | 0.2857 | 0.1071 | 0.0238 |
| | 0, 5 | 1 | 0.5 | 0.2222 | 0.0833 | 0.0238 | 0.0040 |

Mid-p-values for all possible combinations of mutant frequencies in the five parallel cultures evolved under different conditions calculated with a modified Fisher's exact test for association between mutation and condition (Choi *et al.*, 2015). WT indicates the number of parallel cultures that have no mutation (from 0 to 5). Likewise, mutant indicates the number of parallel cultures that have a mutation in the same locus (from 0 to 5). The matrix is symmetric. If all five parallel cultures acquired the same unique mutation that was not present in more than one culture evolved under the other condition (row 0,5 and column 4,1), the association between mutation and condition is statistically significant ($p \leqslant 0.05$). Likewise, the association is significant if four parallel cultures acquired the same unique mutation that was not present in any culture evolved under the other conditions (row 1,4 and column 5,0). It is likely that more mutations would have shown significant associations if the sample size would have been higher than five.

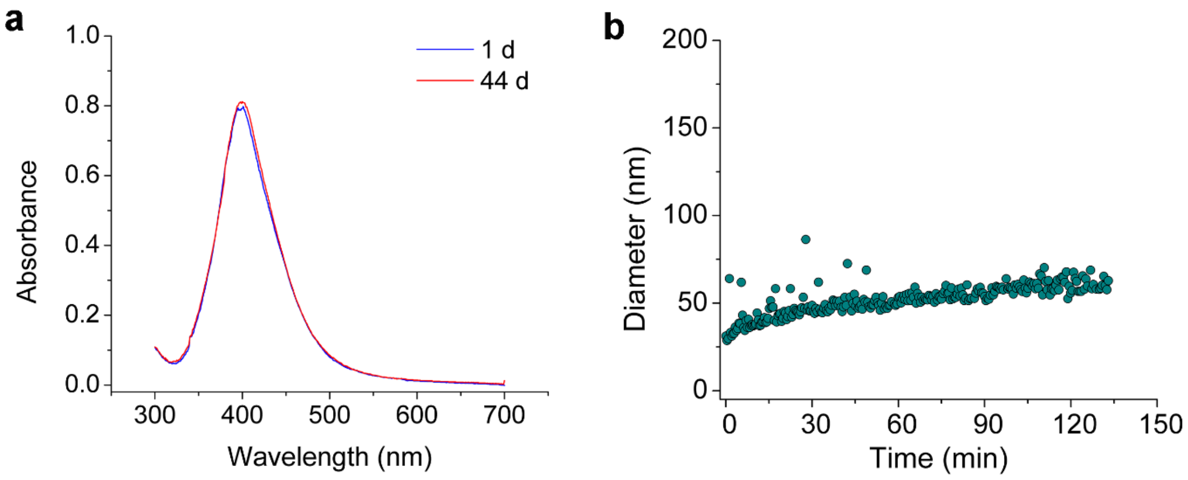


**Figure S1.** Stability of AgNPs. **(a)** UV-Vis absorption spectra of the AgNP stock suspension. **(b)** Hydrodynamic diameter of the AgNP stock suspension in DMM measured by time-resolved DLS, which was recorded at intervals of approximately 0.4 min. Only a slight increase of the hydrodynamic diameter was observed over 133 min (from around 30 to around 60 nm), suggesting that the AgNPs in the stock were relatively stable in DMM.

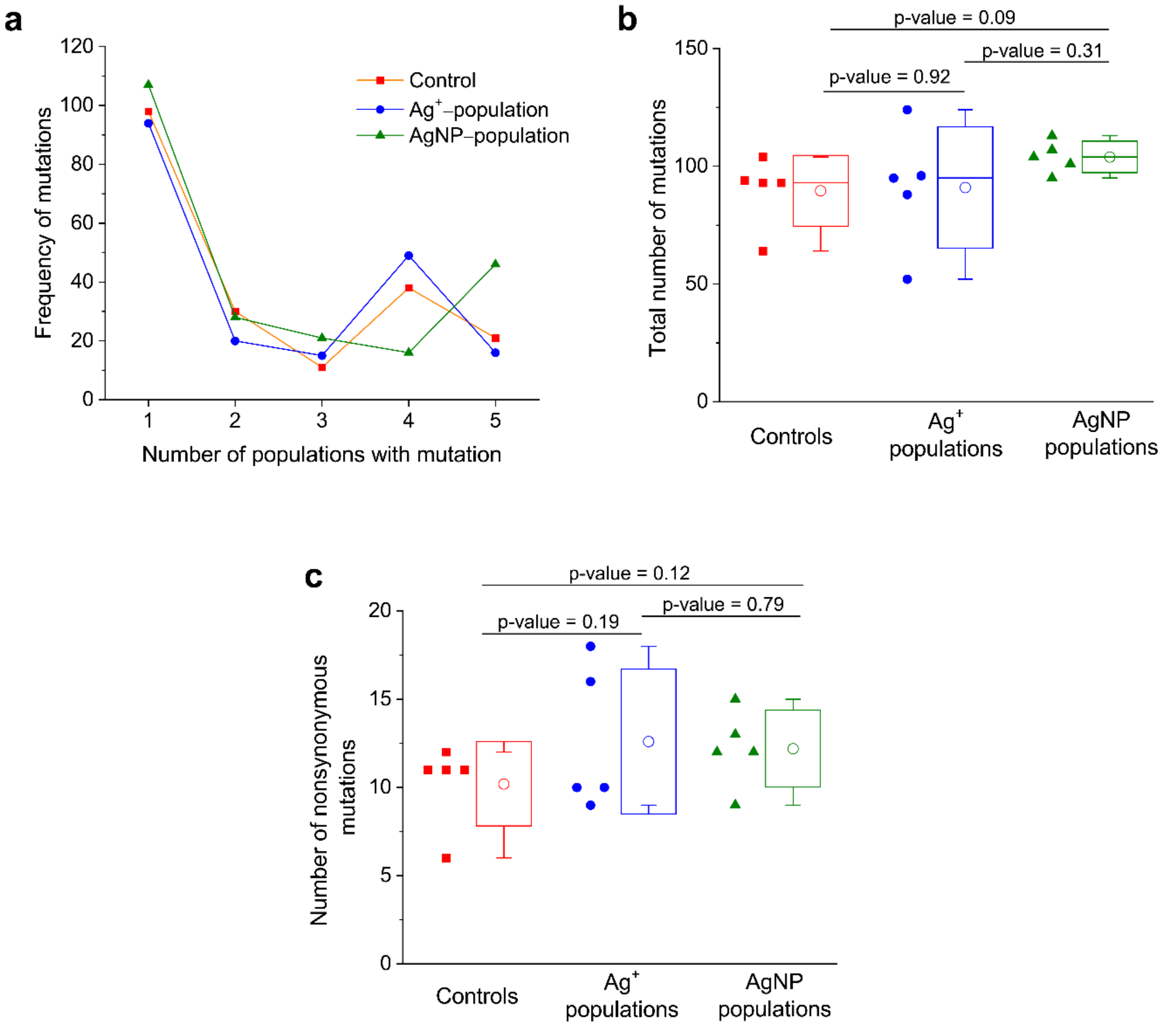


**Figure S2.** Number of mutations in populations that had evolved for 500 generations under $Ag^{+}$ or AgNP stress or without stress (control). **(a)** The frequency of mutations that occurred 1, 2, 3, 4 or 5 times in the five parallel cultures. **(b)** Total number of mutations in the 15 populations. **(c)** Number of nonsynonymous mutations in the coding regions (CDS) in the 15 populations. Open circles are means, boxes SDs, bars medians, and whiskers 5–95 percentiles.

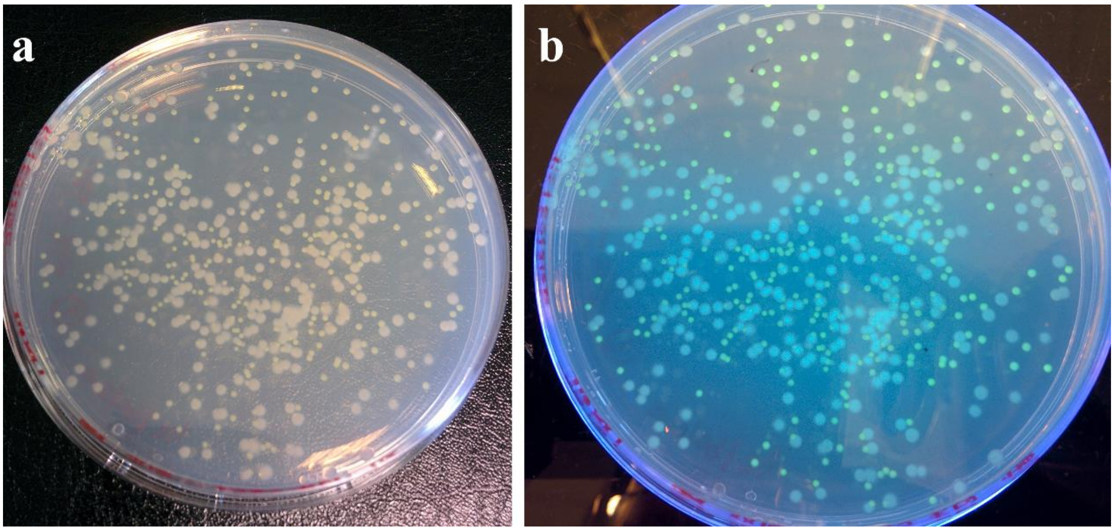


**Figure S3.** Colonies of evolved and reference cells on the same LB plate. The larger, white colonies were formed by evolved cells. Green colonies were formed by the GFP tagged reference cells. **(a)** Under natural light. **(b)** Illuminated by UV light.

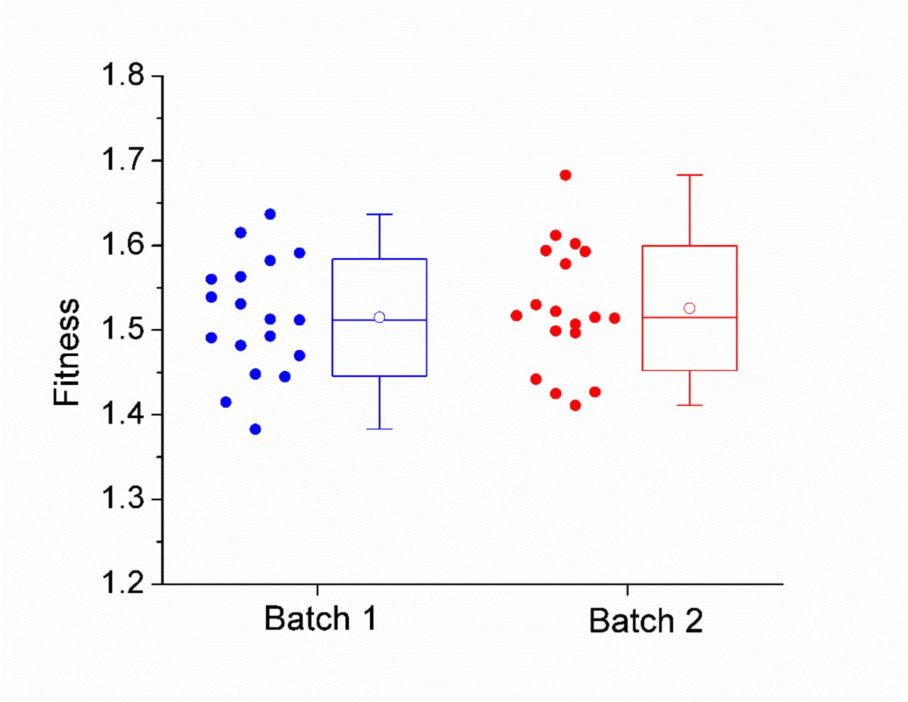


**Figure S4.** Reproducibility of the fitness assay, demonstrating that fitness estimates are not very precise. A population that had evolved without stress (control) in the main evolution experiment (culture 1, generation 500) was competed with the GFP reference strain in Davis minimal medium. Two batches of fitness tests with 18 replicates each were carried out in separate weeks. Data points are individual replicates. Boxes represent SDs, bars medians, open circles means, whiskers 5–95 percentiles. There was no significant difference in the mean and variance of the two batches of fitness values (p-value of 0.636 for mean (paired t-test, two-tailed), p-value of 0.798 for variance (F-test, two-tailed).

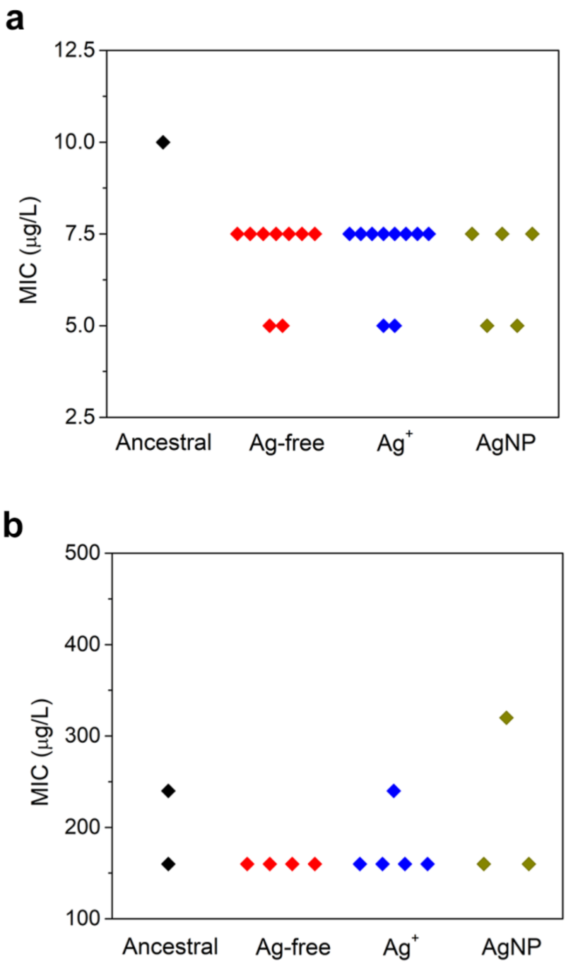


**Figure S5.** Minimum inhibitory concentrations (MICs) of ancestral and parallel Ag-free, $Ag^+$ and AgNP evolved populations with respect to (a) $Ag^+$ or (b) AgNPs, measured on different days. The AgNP MIC refers to the total Ag concentration in the dosed AgNP suspension. MICs were measured by the agar dilution method. No statistically significant differences in MICs were identified between the evolved populations when the $Ag^+$ or AgNP MICs of any two of ancestral, Ag-free, $Ag^+$-evolved and AgNP-evolved population were compared (Mann-Whitney U test, all p-values > 0.28).

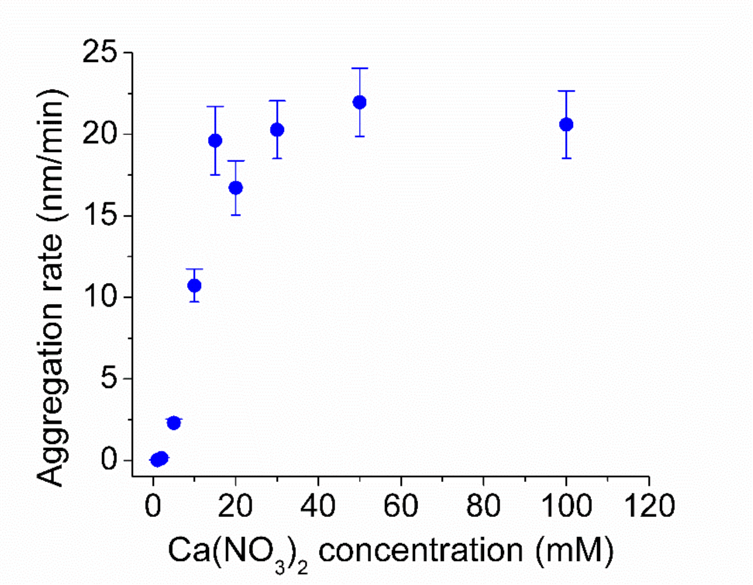


**Figure S6.** Aggregation rates of AgNP stock suspensions in different concentrations of $Ca(NO_3)_2$. To measure dissolved Ag, $Ca(NO_3)_2$ was added to aggregate the washed AgNPs so they could be spun down and removed from the solution as described (Dong *et al.*, 2016).
