## Supplementary Data S1 for "Experimental Evolution of *Pseudomonas putida* under Silver Ion versus Nanoparticle Stress"

SI for Feng Dong, Ana C. Quevedo, Xiang Wang, Eugenia Valsami-Jones & Jan-Ulrich Kreft (2020).Experimental Evolution of Pseudomonas putida under Silver Ion versus Nanoparticle Stress.

List of mutations in the pre-evolved population. The population with the highest fitness after evolving for 465 generations in the absence of AgNO3 or AgNP (culture 2) was compared with the ancestral Pseudomonas putida KT2440 population.

| **Gene name** | **Gene position** | **Gene function or product** | **Mutation Position** | **Effect** | **Functional class** |
|---|---|---|---|---|---|
| Anonymous | | | 178,396 | Intergenic | |
| PP_0168 | 194495-220543 | Putative surface adhesion protein | 195,505 | Synonymous | Silent |
| PP_0168 | 194495-220543 | Putative surface adhesion protein | 195,559 | Synonymous | Silent |
| PP_0168 | 194495-220543 | Putative surface adhesion protein | 195,634 | Synonymous | Silent |
| PP_0168 | 194495-220543 | Putative surface adhesion protein | 196,459 | Synonymous | Silent |
| PP_0168 | 194495-220543 | Putative surface adhesion protein | 196,489 | Synonymous | Silent |
| PP_0168 | 194495-220543 | Putative surface adhesion protein | 196,672 | Synonymous | Silent |
| PP_0168 | 194495-220543 | Putative surface adhesion protein | 196,795 | Synonymous | Silent |
| PP_0168 | 194495-220543 | Putative surface adhesion protein | 196,927 | Synonymous | Silent |
| PP_0168 | 194495-220543 | Putative surface adhesion protein | 197,128 | Synonymous | Silent |
| PP_0168 | 194495-220543 | Putative surface adhesion protein | 197,572 | Synonymous | Silent |
| PP_0168 | 194495-220543 | Putative surface adhesion protein | 197,590 | Synonymous | Silent |
| PP_0168 | 194495-220543 | Putative surface adhesion protein | 197,632 | Synonymous | Silent |
| PP_0168 | 194495-220543 | Putative surface adhesion protein | 197,695 | Synonymous | Silent |
| gacS | 1842040-1844793 | Sensor protein GacS | 1,843,901 | Nonsynonymous | Missense |
| Anonymous | | | 2,069,557 | Intergenic | |
| Anonymous | | | 4,022,306 | Intergenic | |
| Anonymous | | | 4,022,307 | Intergenic | |
| felQ | 4963882-4965357 | Transcriptional regulator FleQ | 4,964,454 | Nonsynonymous | Missense |
| flgK | 4973561-4975603 | Flagellar hook-associated protein FlgK | 4,974,437 | Frame shift | |
| PP_4920 | 5592055-5592801 | Lipoprotein | 5,592,669 | Synonymous | Silent |
| Anonymous | | | 5,988,889 | Intergenic | |
| Anonymous | | | 5,988,905 | Intergenic | |
| Anonymous | | | 5,988,910 | Intergenic | |
