## Supplementary Data S2 for "Experimental Evolution of *Pseudomonas putida* under Silver Ion versus Nanoparticle Stress"

SI for Feng Dong, Ana C. Quevedo, Xiang Wang, Eugenia Valsami-Jones & Jan-Ulrich Kreft (2020).Experimental Evolution of Pseudomonas putida under Silver Ion versus Nanoparticle Stress.

List of all mutations (including synonomous) in the 15 populations after evolving for 500 generations in the main evolution experiment. The reads of the evolved populations were aligned with the *Pseudomonas putida* KT2440 reference genome. Gene names were obtained from the recently updated annotation of the reference genome. The populations evolved under three different conditions: controls (evolved without Ag stress); $Ag^{+}$-populations (evolved in the presence of $Ag^{+}$); AgNP-populations (evolved in the presence of AgNPs). Mutations were only called when they passed quality thresholds: 0 refers to no mutation; 1 refers to mutation. Under each condition (control, $Ag^{+}$ treatment and AgNP treatment), there were five parallel populations. A modified Fisher's exact test was used to calculate the mid-p-values for association of mutation with treatment.

| Gene name | Gene function or product | Mutation position | Reference genome | Mutation | Effect | Codon Change | Amino acid Change | Mutation call: Control C1 | Control C2 | Control C3 | Control C4 | Control C5 | $Ag^{+}$-population C1 | $Ag^{+}$-population C2 | $Ag^{+}$-population C3 | $Ag^{+}$-population C4 | $Ag^{+}$-population C5 | AgNP-population C1 | AgNP-population C2 | AgNP-population C3 | AgNP-population C4 | AgNP-population C5 | Total number of parallel mutations: Control | $Ag^{+}$-population | AgNP-population | p-values for association: $Ag^{+}$-population vs Control | AgNP-population vs Control | AgNP-population vs $Ag^{+}$-population |
|---|---|---|---|---|---|---|---|---|---|---|---|---|---|---|---|---|---|---|---|---|---|---|---|---|---|---|---|---|
| Anonymous | | 172,966 | A | G | Intergenic | | | 1 | 1 | 1 | 1 | 1 | 1 | 1 | 1 | 1 | 0 | 0 | 1 | 1 | 1 | 1 | 5 | 4 | 4 | 0.50 | 0.5 | 1 |
| Anonymous | | 173,074 | A | G | Intergenic | | | 0 | 0 | 0 | 0 | 0 | 1 | 0 | 0 | 0 | 0 | 1 | 1 | 1 | 1 | 0 | 0 | 1 | 4 | 0.5 | 0.02381 | 0.1071 |
| Anonymous | | 178,396 | A | G | Intergenic | | | 1 | 1 | 1 | 0 | 0 | 1 | 1 | 1 | 1 | 0 | 1 | 0 | 0 | 1 | 0 | 3 | 4 | 2 | 0.5833 | 0.6032 | 0.2857 |
| Anonymous | | 178,437 | C | T | Intergenic | | | 1 | 1 | 1 | 1 | 1 | 1 | 1 | 1 | 1 | 0 | 1 | 1 | 0 | 1 | 0 | 5 | 4 | 3 | 0.5 | 0.2222 | 0.5833 |
| Anonymous | | 178,504 | A | G | Intergenic | | | 1 | 0 | 0 | 0 | 0 | 0 | 0 | 1 | 1 | 0 | 1 | 1 | 1 | 1 | 1 | 1 | 2 | 5 | 0.5833 | 0.02381 | 0.08333 |
| PP_0168 | Putative surface adhesion protein | 195,406 | C | T | Synonymous_Coding | aaC/aaT | N304 | 0 | 0 | 0 | 0 | 1 | 0 | 0 | 0 | 0 | 0 | 0 | 1 | 0 | 1 | 0 | 1 | 0 | 2 | 0.5 | 0.5833 | 0.2222 |
| PP_0168 | Putative surface adhesion protein | 195,424 | G | A | Synonymous_Coding | gtG/gtA | V310 | 0 | 1 | 1 | 0 | 1 | 1 | 0 | 0 | 1 | 0 | 1 | 0 | 1 | 1 | 0 | 3 | 2 | 3 | 0.6032 | 1 | 0.6032 |
| PP_0168 | Putative surface adhesion protein | 195,449 | G | A | Nonsynonymous_Coding | Gtc/Atc | V319I | 0 | 0 | 0 | 0 | 0 | 1 | 0 | 0 | 0 | 0 | 0 | 0 | 1 | 0 | 0 | 0 | 1 | 1 | 0.5 | 0.5 | 1 |
| PP_0168 | Putative surface adhesion protein | 19,450 | T | C | Nonsynonymous_Coding | gTc/gCc | V319A | 0 | 0 | 0 | 0 | 0 | 1 | 0 | 1 | 0 | 1 | 0 | 0 | 0 | 0 | 0 | 0 | 3 | 0 | 0.08333 | 1 | 0.08333 |
| PP_0168 | Putative surface adhesion protein | 195,457 | T | C | Synonymous_Coding | acT/acC | T321 | 0 | 0 | 0 | 0 | 1 | 1 | 0 | 0 | 0 | 0 | 0 | 0 | 0 | 0 | 0 | 1 | 1 | 0 | 1 | 0.5 | 0.5 |
| PP_0168 | Putative surface adhesion protein | 195,472 | A | G | Synonymous_Coding | aaA/aaG | K326 | 0 | 1 | 1 | 0 | 0 | 0 | 0 | 0 | 0 | 0 | 0 | 0 | 0 | 0 | 0 | 2 | 0 | 0 | 0.2222 | 0.2222 | 1 |
| PP_0168 | Putative surface adhesion protein | 195,505 | C | G | Synonymous_Coding | gcC/gcG | A337 | 1 | 0 | 1 | 1 | 1 | 1 | 0 | 0 | 1 | 1 | 1 | 0 | 1 | 1 | 1 | 4 | 3 | 4 | 0.5833 | 1 | 0.5833 |
| PP_0168 | Putative surface adhesion protein | 195,559 | C | A | Synonymous_Coding | acC/acA | T355 | 1 | 1 | 1 | 1 | 1 | 1 | 1 | 1 | 1 | 1 | 1 | 0 | 1 | 0 | 1 | 5 | 5 | 3 | 1 | 0.2222 | 0.2222 |
| PP_0168 | Putative surface adhesion protein | 195,568 | T | C | Synonymous_Coding | aaT/aaC | N358 | 0 | 0 | 0 | 0 | 0 | 0 | 0 | 0 | 1 | 0 | 0 | 0 | 0 | 0 | 0 | 0 | 1 | 0 | 0.5 | 1 | 0.5 |
| PP_0168 | Putative surface adhesion protein | 195,589 | T | C | Synonymous_Coding | aaT/aaC | N365 | 0 | 0 | 1 | 0 | 1 | 0 | 1 | 1 | 1 | 1 | 1 | 1 | 1 | 1 | 1 | 2 | 4 | 5 | 0.2857 | 0.08333 | 1 |
| PP_0168 | Putative surface adhesion protein | 195,628 | C | T | Synonymous_Coding | gaC/gaT | D378 | 1 | 1 | 1 | 1 | 1 | 1 | 0 | 1 | 0 | 0 | 0 | 1 | 0 | 1 | 0 | 5 | 2 | 2 | 0.08333 | 0.08333 | 1 |
| PP_0168 | Putative surface adhesion protein | 195,629 | A | G | Non_Synonymous_Coding | Acc/Gcc | T379A | 1 | 1 | 1 | 1 | 1 | 1 | 0 | 1 | 0 | 1 | 0 | 1 | 0 | 1 | 0 | 5 | 3 | 2 | 0.2222 | 0.08333 | 0.6032 |
| PP_0168 | Putative surface adhesion protein | 195,634 | C | T | Synonymous_Coding | acC/acT | T380 | 1 | 1 | 1 | 1 | 1 | 1 | 1 | 1 | 1 | 1 | 1 | 0 | 1 | 1 | 1 | 5 | 5 | 4 | 1 | 0.5 | 0.5 |
| PP_0168 | Putative surface adhesion protein | 195,685 | T | C | Synonymous_Coding | acT/acC | T397 | 1 | 0 | 1 | 0 | 0 | 0 | 1 | 1 | 1 | 1 | 1 | 1 | 0 | 1 | 1 | 2 | 4 | 4 | 0.2857 | 0.2857 | 1 |
| PP_0168 | Putative surface adhesion protein | 195,724 | A | G | Synonymous_Coding | gtA/gtG | V410 | 0 | 0 | 0 | 0 | 0 | 1 | 0 | 1 | 0 | 0 | 1 | 0 | 0 | 0 | 0 | 0 | 2 | 1 | 0.2222 | 0.5 | 0.5833 |
| PP_0168 | Putative surface adhesion protein | 195,732 | A | C | Non_Synonymous_Coding | aAc/aCc | N413T | 1 | 1 | 1 | 0 | 1 | 0 | 1 | 1 | 1 | 1 | 1 | 1 | 1 | 1 | 1 | 4 | 4 | 5 | 1 | 1 | 1 |
| PP_0168 | Putative surface adhesion protein | 195,766 | G | C | Synonymous_Coding | gcG/gcC | A424 | 0 | 1 | 0 | 0 | 0 | 1 | 0 | 0 | 0 | 0 | 0 | 0 | 0 | 0 | 1 | 1 | 1 | 1 | 1 | 1 | 1 |
| PP_0168 | Putative surface adhesion protein | 195,769 | T | C | Synonymous_Coding | ggT/ggC | G425 | 0 | 1 | 1 | 0 | 1 | 1 | 0 | 0 | 0 | 0 | 0 | 0 | 1 | 0 | 1 | 3 | 1 | 2 | 0.2857 | 0.6032 | 0.5833 |
| PP_0168 | Putative surface adhesion protein | 195,805 | C | G | Synonymous_Coding | gcC/gcG | A437 | 0 | 1 | 0 | 0 | 0 | 0 | 0 | 0 | 0 | 0 | 0 | 0 | 0 | 0 | 0 | 1 | 0 | 0 | 0.5 | 0.5 | 1 |
| PP_0168 | Putative surface adhesion protein | 195,826 | C | T | Synonymous_Coding | ggC/ggT | G444 | 0 | 0 | 0 | 0 | 1 | 0 | 0 | 0 | 1 | 0 | 0 | 0 | 0 | 0 | 1 | 1 | 1 | 1 | 1 | 1 | 1 |
| PP_0168 | Putative surface adhesion protein | 195,852 | G | A | Nonsynonymous_Coding | aGc/aAc | S453N | 0 | 1 | 1 | 1 | 1 | 1 | 0 | 0 | 0 | 0 | 0 | 1 | 0 | 0 | 0 | 4 | 1 | 1 | 0.1071 | 0.1071 | 1 |
| PP_0168 | Putative surface adhesion protein | 195,877 | G | A | Synonymous_Coding | caG/caA | Q461 | 0 | 0 | 0 | 1 | 0 | 0 | 0 | 0 | 0 | 0 | 0 | 0 | 0 | 0 | 0 | 1 | 0 | 0 | 0.5 | 0.5 | 1 |
| PP_0168 | Putative surface adhesion protein | 195,929 | A | G | Nonsynonymous_Coding | Acc/Gcc | T479A | 0 | 0 | 1 | 0 | 0 | 0 | 0 | 0 | 0 | 0 | 0 | 0 | 0 | 0 | 0 | 1 | 0 | 0 | 0.5 | 0.5 | 1 |
| PP_0168 | Putative surface adhesion protein | 195,934 | C | T | Synonymous_Coding | acC/acT | T480 | 0 | 0 | 0 | 0 | 0 | 0 | 0 | 0 | 0 | 0 | 0 | 0 | 0 | 0 | 1 | 0 | 0 | 1 | 1 | 0.5 | 0.5 |
| PP_0168 | Putative surface adhesion protein | 195,979 | G | A | Synonymous_Coding | gtG/gtA | V495 | 0 | 0 | 0 | 0 | 0 | 0 | 0 | 1 | 0 | 0 | 0 | 0 | 0 | 0 | 0 | 0 | 1 | 0 | 0.5 | 1 | 0.5 |
| PP_0168 | Putative surface adhesion protein | 195,985 | C | T | Synonymous_Coding | acC/acT | T497 | 0 | 0 | 0 | 0 | 0 | 0 | 0 | 0 | 0 | 0 | 0 | 0 | 0 | 0 | 1 | 0 | 0 | 1 | 1 | 0.5 | 0.5 |
| PP_0168 | Putative surface adhesion protein | 196,027 | T | C | Synonymous_Coding | acT/acC | T511 | 1 | 1 | 1 | 0 | 0 | 1 | 0 | 1 | 1 | 1 | 1 | 1 | 1 | 1 | 1 | 3 | 4 | 5 | 0.5833 | 0.2222 | 1 |
| PP_0168 | Putative surface adhesion protein | 196,051 | G | C | Synonymous_Coding | acG/acC | T519 | 0 | 1 | 0 | 0 | 0 | 0 | 0 | 0 | 0 | 0 | 0 | 0 | 0 | 0 | 0 | 1 | 0 | 0 | 0.5 | 0.5 | 1 |
| PP_0168 | Putative surface adhesion protein | 196,066 | G | C | Synonymous_Coding | gcG/gcC | A524 | 0 | 0 | 0 | 0 | 0 | 0 | 0 | 0 | 0 | 0 | 1 | 1 | 0 | 0 | 0 | 0 | 0 | 2 | 1 | 0.2222 | 0.2222 |
| PP_0168 | Putative surface adhesion protein | 196,069 | T | C | Synonymous_Coding | ggT/ggC | G525 | 0 | 0 | 1 | 0 | 0 | 0 | 0 | 0 | 1 | 0 | 0 | 1 | 0 | 0 | 0 | 1 | 1 | 1 | 1 | 1 | 1 |
| PP_0168 | Putative surface adhesion protein | 196,132 | G | C | Synonymous_Coding | acG/acC | T546 | 0 | 0 | 1 | 0 | 1 | 1 | 1 | 0 | 1 | 1 | 1 | 1 | 1 | 1 | 1 | 2 | 4 | 5 | 0.2857 | 0.08333 | 1 |
| PP_0168 | Putative surface adhesion protein | 196,159 | G | A | Synonymous_Coding | acG/acA | T555 | 1 | 0 | 1 | 0 | 1 | 1 | 0 | 0 | 1 | 1 | 0 | 0 | 1 | 1 | 1 | 3 | 3 | 3 | 1 | 1 | 1 |
| PP_0168 | Putative surface adhesion protein | 196,195 | G | C | Synonymous_Coding | acG/acC | T567 | 1 | 0 | 1 | 0 | 0 | 0 | 1 | 0 | 0 | 0 | 0 | 0 | 0 | 1 | 1 | 2 | 1 | 2 | 0.5833 | 1 | 0.5833 |
| PP_0168 | Putative surface adhesion protein | 196,234 | T | C | Synonymous_Coding | acT/acC | T580 | 1 | 0 | 1 | 1 | 1 | 1 | 1 | 1 | 1 | 0 | 1 | 1 | 1 | 1 | 1 | 4 | 4 | 5 | 1 | 1 | 1 |
| PP_0168 | Putative surface adhesion protein | 196,285 | T | C | Synonymous_Coding | acT/acC | T597 | 1 | 0 | 0 | 0 | 1 | 1 | 1 | 1 | 1 | 0 | 0 | 1 | 0 | 1 | 1 | 2 | 4 | 3 | 0.2857 | 0.6032 | 0.5833 |
| PP_0168 | Putative surface adhesion protein | 196,306 | T | C | Synonymous_Coding | aaT/aaC | N604 | 1 | 1 | 1 | 1 | 1 | 1 | 1 | 1 | 0 | 1 | 1 | 1 | 1 | 1 | 1 | 5 | 4 | 5 | 0.5 | 1 | 1 |
| PP_0168 | Putative surface adhesion protein | 196,324 | G | A | Synonymous_Coding | gtG/gtA | V610 | 0 | 0 | 0 | 0 | 0 | 0 | 0 | 0 | 0 | 0 | 0 | 1 | 0 | 1 | 1 | 0 | 0 | 3 | 1 | 0.08333 | 0.08333 |
| PP_0168 | Putative surface adhesion protein | 196,327 | A | C | Synonymous_Coding | acA/acC | T611 | 1 | 1 | 1 | 1 | 1 | 1 | 1 | 1 | 1 | 0 | 1 | 1 | 1 | 1 | 1 | 5 | 4 | 5 | 0.5 | 1 | 1 |
| PP_0168 | Putative surface adhesion protein | 196,351 | C | G | Synonymous_Coding | acC/acG | T619 | 1 | 1 | 1 | 1 | 1 | 1 | 1 | 1 | 0 | 1 | 1 | 1 | 1 | 1 | 1 | 5 | 4 | 5 | 0.5 | 1 | 1 |
| PP_0168 | Putative surface adhesion protein | 196,366 | C | G | Synonymous_Coding | gcC/gcG | A624 | 0 | 0 | 0 | 0 | 0 | 0 | 0 | 0 | 0 | 0 | 0 | 1 | 0 | 0 | 0 | 0 | 0 | 1 | 1 | 0.5 | 0.5 |
| PP_0168 | Putative surface adhesion protein | 196,369 | C | T | Synonymous_Coding | ggC/ggT | G625 | 0 | 0 | 0 | 0 | 0 | 0 | 0 | 0 | 0 | 0 | 0 | 1 | 0 | 0 | 0 | 0 | 0 | 1 | 1 | 0.5 | 0.5 |
| PP_0168 | Putative surface adhesion protein | 196,372 | A | G | Synonymous_Coding | aaA/aaG | K626 | 1 | 1 | 1 | 1 | 1 | 1 | 1 | 1 | 1 | 1 | 1 | 1 | 1 | 1 | 1 | 5 | 5 | 5 | 1 | 1 | 1 |
| PP_0168 | Putative surface adhesion protein | 196,390 | T | C | Synonymous_Coding | gaT/gaC | D632 | 1 | 1 | 1 | 0 | 1 | 0 | 1 | 1 | 0 | 1 | 1 | 0 | 1 | 1 | 1 | 4 | 3 | 4 | 0.5833 | 1 | 0.5833 |
| PP_0168 | Putative surface adhesion protein | 196,405 | G | C | Synonymous_Coding | gcG/gcC | A637 | 0 | 0 | 0 | 0 | 0 | 0 | 0 | 0 | 0 | 0 | 0 | 0 | 0 | 1 | 0 | 0 | 0 | 1 | 1 | 0.5 | 0.5 |
| PP_0168 | Putative surface adhesion protein | 196,432 | T | C | Synonymous_Coding | acT/acC | T646 | 0 | 1 | 0 | 1 | 0 | 0 | 0 | 0 | 0 | 0 | 0 | 0 | 0 | 0 | 1 | 2 | 0 | 1 | 0.2222 | 0.5833 | 0.5 |
| PP_0168 | Putative surface adhesion protein | 196,459 | C | A | Synonymous_Coding | acC/acA | T655 | 0 | 1 | 0 | 1 | 1 | 1 | 1 | 1 | 0 | 1 | 1 | 1 | 1 | 1 | 1 | 3 | 4 | 5 | 0.5833 | 0.2222 | 1 |
| PP_0168 | Putative surface adhesion protein | 196,489 | T | C | Synonymous_Coding | aaT/aaC | N665 | 1 | 1 | 1 | 0 | 1 | 1 | 1 | 1 | 1 | 0 | 1 | 1 | 1 | 1 | 1 | 4 | 4 | 5 | 1 | 1 | 1 |
| PP_0168 | Putative surface adhesion protein | 196,495 | G | C | Synonymous_Coding | acG/acC | T667 | 1 | 1 | 1 | 1 | 1 | 1 | 1 | 1 | 1 | 0 | 1 | 1 | 1 | 1 | 1 | 5 | 4 | 5 | 0.5 | 1 | 1 |
| PP_0168 | Putative surface adhesion protein | 196,528 | T | C | Synonymous_Coding | gaT/gaC | D678 | 0 | 0 | 0 | 0 | 0 | 0 | 0 | 0 | 0 | 0 | 0 | 1 | 0 | 0 | 0 | 0 | 0 | 1 | 1 | 0.5 | 0.5 |
| PP_0168 | Putative surface adhesion protein | 196,579 | G | A | Synonymous_Coding | gtG/gtA | V695 | 1 | 1 | 1 | 1 | 1 | 1 | 1 | 1 | 1 | 1 | 1 | 1 | 1 | 1 | 1 | 5 | 5 | 5 | 1 | 1 | 1 |

| | | | | | | | | | | | | | | | | | | | | | | | | | | | | |
|---|---|---|---|---|---|---|---|---|---|---|---|---|---|---|---|---|---|---|---|---|---|---|---|---|---|---|---|---|
| PP_0168 | Putative surface adhesion protein | 196,585 | C | T | Synonymous_Coding | acC/acT | T697 | 0 | 0 | 1 | 1 | 0 | 1 | 1 | 0 | 1 | 0 | 0 | 0 | 1 | 0 | 0 | 2 | 3 | 1 | 0.6032 | 0.5833 | 0.2857 |
| PP_0168 | Putative surface adhesion protein | 196,606 | T | C | Synonymous_Coding | aaT/aaC | N704 | 0 | 0 | 0 | 1 | 0 | 0 | 0 | 0 | 1 | 0 | 0 | 0 | 0 | 0 | 0 | 1 | 1 | 0 | 1 | 0.5 | 0.5 |
| PP_0168 | Putative surface adhesion protein | 196,624 | G | A | Synonymous_Coding | gtG/gtA | V710 | 0 | 0 | 0 | 1 | 1 | 0 | 0 | 0 | 1 | 0 | 0 | 0 | 0 | 0 | 0 | 2 | 1 | 0 | 0.5833 | 0.2222 | 0.5 |
| PP_0168 | Putative surface adhesion protein | 196,627 | A | C | Synonymous_Coding | acA/acC | T711 | 1 | 1 | 1 | 1 | 1 | 1 | 1 | 1 | 1 | 1 | 0 | 1 | 1 | 1 | 1 | 5 | 5 | 4 | 1 | 0.5 | 0.5 |
| PP_0168 | Putative surface adhesion protein | 196,672 | G | A | Synonymous_Coding | aaG/aaA | K726 | 1 | 0 | 0 | 0 | 1 | 0 | 1 | 0 | 0 | 0 | 1 | 0 | 0 | 0 | 0 | 2 | 1 | 1 | 0.5833 | 0.5833 | 1 |
| PP_0168 | Putative surface adhesion protein | 196,705 | G | C | Synonymous_Coding | gcG/gcC | A737 | 1 | 1 | 1 | 0 | 0 | 0 | 1 | 1 | 0 | 0 | 0 | 0 | 0 | 1 | 1 | 3 | 2 | 2 | 0.6032 | 0.6032 | 1 |
| PP_0168 | Putative surface adhesion protein | 196,752 | A | G | Nonsynonymous_Coding | aAc/aGc | N753S | 0 | 0 | 0 | 0 | 0 | 0 | 0 | 0 | 0 | 1 | 0 | 0 | 0 | 0 | 0 | 0 | 1 | 0 | 0.5 | 1 | 0.5 |
| PP_0168 | Putative surface adhesion protein | 196,759 | T | C,A | Synonymous_Coding | acT/acC | T755 | 1 | 1 | 1 | 0 | 1 | 1 | 1 | 1 | 0 | 1 | 1 | 1 | 0 | 1 | 1 | 4 | 4 | 4 | 1 | 1 | 1 |
| PP_0168 | Putative surface adhesion protein | 196,795 | G | C | Synonymous_Coding | acG/acC | T767 | 0 | 0 | 0 | 0 | 1 | 0 | 0 | 1 | 0 | 0 | 0 | 1 | 0 | 0 | 0 | 1 | 1 | 1 | 1 | 1 | 1 |
| PP_0168 | Putative surface adhesion protein | 196,828 | T | C | Synonymous_Coding | gaT/gaC | D778 | 0 | 1 | 0 | 0 | 0 | 0 | 0 | 0 | 0 | 0 | 0 | 1 | 0 | 0 | 0 | 1 | 0 | 1 | 0.5 | 1 | 0.5 |
| PP_0168 | Putative surface adhesion protein | 196,829 | G | A | Nonsynonymous_Coding | Gcc/Acc | A779T | 0 | 1 | 0 | 0 | 0 | 0 | 0 | 0 | 0 | 0 | 0 | 0 | 0 | 0 | 0 | 1 | 0 | 0 | 0.5 | 0.5 | 1 |
| PP_0168 | Putative surface adhesion protein | 196,885 | C | T | Synonymous_Coding | acC/acT | T797 | 1 | 1 | 1 | 1 | 0 | 0 | 1 | 1 | 1 | 1 | 1 | 1 | 1 | 0 | 0 | 4 | 4 | 3 | 1 | 0.5833 | 0.5833 |
| PP_0168 | Putative surface adhesion protein | 196,906 | T | C | Synonymous_Coding | aaT/aaC | N804 | 0 | 0 | 1 | 0 | 0 | 0 | 0 | 0 | 0 | 0 | 0 | 0 | 0 | 0 | 0 | 1 | 0 | 0 | 0.5 | 0.5 | 1 |
| PP_0168 | Putative surface adhesion protein | 196,924 | G | A | Synonymous_Coding | gtG/gtA | V810 | 0 | 0 | 0 | 1 | 0 | 0 | 0 | 0 | 0 | 0 | 0 | 0 | 0 | 0 | 0 | 1 | 0 | 0 | 0.5 | 0.5 | 1 |
| PP_0168 | Putative surface adhesion protein | 196,927 | A | C | Synonymous_Coding | acA/acC | T811 | 1 | 1 | 1 | 1 | 1 | 1 | 1 | 1 | 1 | 0 | 1 | 1 | 1 | 1 | 1 | 5 | 4 | 5 | 0.5 | 1 | 1 |
| PP_0168 | Putative surface adhesion protein | 196,972 | G | A | Synonymous_Coding | aaG/aaA | K826 | 1 | 1 | 1 | 1 | 1 | 1 | 1 | 0 | 1 | 0 | 1 | 1 | 1 | 1 | 0 | 5 | 3 | 4 | 0.2222 | 0.5 | 0.5833 |
| PP_0168 | Putative surface adhesion protein | 197,128 | T | C | Synonymous_Coding | gaT/gaC | D878 | 0 | 0 | 0 | 0 | 0 | 0 | 0 | 0 | 0 | 0 | 1 | 1 | 0 | 0 | 0 | 0 | 0 | 2 | 1 | 0.2222 | 0.2222 |
| PP_0168 | Putative surface adhesion protein | 197,129 | G | A | Non_Synonymous_Coding | Gcc/Acc | A879T | 0 | 0 | 0 | 0 | 0 | 0 | 0 | 0 | 0 | 0 | 1 | 1 | 0 | 0 | 0 | 0 | 0 | 2 | 1 | 0.2222 | 0.2222 |
| PP_0168 | Putative surface adhesion protein | 197,179 | A | G | Synonymous_Coding | gtA/gtG | V895 | 1 | 1 | 1 | 1 | 0 | 1 | 1 | 1 | 1 | 1 | 1 | 1 | 1 | 1 | 1 | 4 | 5 | 5 | 1 | 1 | 1 |
| PP_0168 | Putative surface adhesion protein | 197,185 | C | T | Synonymous_Coding | acC/acT | T897 | | | | | | | | | | | | | | | | 0 | 0 | 0 | 1 | 1 | 1 |
| PP_0168 | Putative surface adhesion protein | 197,206 | T | C | Synonymous_Coding | aaT/aaC | N904 | 0 | 0 | 0 | 0 | 0 | 0 | 0 | 0 | 1 | 0 | 0 | 0 | 0 | 0 | 0 | 0 | 1 | 0 | 0.5 | 1 | 0.5 |
| PP_0168 | Putative surface adhesion protein | 197,224 | A | G | Synonymous_Coding | gtA/gtG | V910 | 1 | 0 | 0 | 0 | 0 | 1 | 0 | 1 | 1 | 1 | 1 | 0 | 0 | 0 | 1 | 1 | 4 | 2 | 0.1071 | 0.5833 | 0.2857 |
| PP_0168 | Putative surface adhesion protein | 197,227 | C | A | Synonymous_Coding | acC/acA | T911 | 0 | 0 | 0 | 0 | 0 | 0 | 0 | 0 | 0 | 0 | 0 | 0 | 0 | 0 | 1 | 0 | 0 | 1 | 1 | 0.5 | 0.5 |
| PP_0168 | Putative surface adhesion protein | 197,250 | T | C | Nonsynonymous_Coding | gTc/gCc | V919A | 0 | 0 | 0 | 0 | 1 | 0 | 0 | 0 | 0 | 0 | 0 | 0 | 0 | 0 | 0 | 1 | 0 | 0 | 0.5 | 0.5 | 1 |
| PP_0168 | Putative surface adhesion protein | 197,257 | T | C | Synonymous_Coding | acT/acC | T921 | 1 | 0 | 0 | 0 | 0 | 0 | 0 | 0 | 0 | 1 | 1 | 0 | 0 | 0 | 0 | 1 | 1 | 1 | 1 | 1 | 1 |
| PP_0168 | Putative surface adhesion protein | 197,326 | T | C | Synonymous_Coding | ggT/ggC | G944 | 0 | 0 | 0 | 0 | 0 | 0 | 0 | 0 | 0 | 1 | 0 | 0 | 0 | 0 | 0 | 0 | 1 | 0 | 0.5 | 1 | 0.5 |
| PP_0168 | Putative surface adhesion protein | 197,395 | G | C | Synonymous_Coding | acG/acC | T967 | 0 | 0 | 1 | 0 | 0 | 1 | 0 | 0 | 0 | 0 | 0 | 0 | 0 | 0 | 0 | 1 | 1 | 0 | 1 | 0.5 | 0.5 |
| PP_0168 | Putative surface adhesion protein | 197,479 | G | A | Synonymous_Coding | gtG/gtA | V995 | 0 | 1 | 1 | 0 | 0 | 0 | 0 | 0 | 0 | 0 | 0 | 0 | 0 | 0 | 0 | 2 | 0 | 0 | 0.2222 | 0.2222 | 1 |
| PP_0168 | Putative surface adhesion protein | 197,496 | T | C | Non_Synonymous_Coding | aTc/aCc | I1001T | 0 | 1 | 1 | 1 | 1 | 1 | 1 | 1 | 1 | 1 | 1 | 1 | 0 | 1 | 1 | 4 | 5 | 4 | 1 | 1 | 0.5 |
| PP_0168 | Putative surface adhesion protein | 197,524 | A | G | Synonymous_Coding | gtA/gtG | V1010 | 0 | 0 | 0 | 1 | 0 | 0 | 0 | 0 | 0 | 1 | 0 | 0 | 0 | 0 | 0 | 1 | 1 | 0 | 1 | 0.5 | 0.5 |
| PP_0168 | Putative surface adhesion protein | 197,551 | C | G | Synonymous_Coding | acC/acG | T1019 | 0 | 0 | 0 | 0 | 0 | 0 | 0 | 0 | 1 | 0 | 0 | 0 | 0 | 1 | 0 | 0 | 1 | 1 | 0.5 | 0.5 | 1 |
| PP_0168 | Putative surface adhesion protein | 197,572 | G | A | Synonymous_Coding | aaG/aaA | K1026 | 1 | 1 | 0 | 1 | 1 | 1 | 1 | 1 | 1 | 1 | 1 | 1 | 1 | 1 | 0 | 4 | 5 | 4 | 1 | 1 | 0.5 |
| PP_0168 | Putative surface adhesion protein | 197,590 | T | C | Synonymous_Coding | gaT/gaC | D1032 | 1 | 0 | 0 | 1 | 1 | 1 | 1 | 1 | 1 | 1 | 1 | 1 | 0 | 1 | 0 | 3 | 5 | 3 | 0.2222 | 1 | 0.2222 |
| PP_0168 | Putative surface adhesion protein | 197,605 | G | C | Synonymous_Coding | gcG/gcC | A1037 | 0 | 0 | 0 | 0 | 0 | 0 | 0 | 1 | 0 | 0 | 0 | 0 | 0 | 0 | 0 | 0 | 1 | 0 | 0.5 | 1 | 0.5 |
| PP_0168 | Putative surface adhesion protein | 197,632 | T | C | Synonymous_Coding | acT/acC | T1046 | 1 | 1 | 1 | 1 | 1 | 1 | 1 | 1 | 1 | 1 | 1 | 1 | 1 | 1 | 1 | 5 | 5 | 5 | 1 | 1 | 1 |
| PP_0168 | Putative surface adhesion protein | 197,665 | T | C | Synonymous_Coding | ggT/ggC | G1057 | 0 | 0 | 0 | 0 | 0 | 0 | 0 | 1 | 0 | 0 | 0 | 0 | 0 | 0 | 0 | 0 | 1 | 0 | 0.5 | 1 | 0.5 |
| PP_0168 | Putative surface adhesion protein | 197,689 | C | T | Synonymous_Coding | aaC/aaT | N1065 | 1 | 0 | 0 | 0 | 0 | 0 | 0 | 0 | 0 | 0 | 0 | 0 | 0 | 0 | 0 | 1 | 0 | 0 | 0.5 | 0.5 | 1 |
| PP_0168 | Putative surface adhesion protein | 197,695 | G | C | Synonymous_Coding | acG/acC | T1067 | 1 | 1 | 1 | 0 | 0 | 0 | 1 | 0 | 1 | 0 | 1 | 1 | 0 | 1 | 1 | 3 | 2 | 4 | 0.6032 | 0.5833 | 0.2857 |
| PP_0168 | Putative surface adhesion protein | 197,715 | C | A | Nonsynonymous_Coding | aCc/aAc | T1074N | 1 | 1 | 0 | 0 | 0 | 0 | 1 | 0 | 0 | 0 | 0 | 1 | 0 | 1 | 0 | 2 | 1 | 2 | 0.5833 | 1 | 0.5833 |
| PP_0168 | Putative surface adhesion protein | 197,834 | T | C | Synonymous_Coding | Ttg/Ctg | L1114 | 1 | 1 | 1 | 1 | 1 | 1 | 1 | 1 | 0 | 0 | 1 | 1 | 1 | 1 | 1 | 5 | 3 | 5 | 0.2222 | 1 | 0.2222 |
| PP_0168 | Putative surface adhesion protein | 197,866 | G | C | Synonymous_Coding | gcG/gcC | A1124 | 0 | 1 | 1 | 1 | 0 | 0 | 0 | 0 | 0 | 1 | 1 | 0 | 1 | 0 | 1 | 3 | 1 | 3 | 0.2857 | 1 | 0.2857 |
| PP_0168 | Putative surface adhesion protein | 197,869 | T | C | Synonymous_Coding | ggT/ggC | G1125 | 0 | 1 | 1 | 1 | 0 | 1 | 1 | 0 | 0 | 1 | 1 | 1 | 1 | 0 | 0 | 3 | 3 | 3 | 1 | 1 | 1 |
| PP_0168 | Putative surface adhesion protein | 197,872 | A | G | Synonymous_Coding | aaA/aaG | K1126 | 0 | 0 | 0 | 0 | 0 | 0 | 0 | 0 | 0 | 0 | 0 | 1 | 0 | 0 | 0 | 0 | 0 | 1 | 1 | 0.5 | 0.5 |
| PP_0168 | Putative surface adhesion protein | 197,926 | C | T | Synonymous_Coding | ggC/ggT | G1144 | 0 | 0 | 0 | 1 | 0 | 0 | 0 | 0 | 0 | 1 | 0 | 0 | 0 | 0 | 0 | 1 | 1 | 0 | 1 | 0.5 | 0.5 |
| PP_0168 | Putative surface adhesion protein | 198,400 | C | G | Synonymous_Coding | acC/acG | T1302 | 0 | 0 | 0 | 1 | 0 | 0 | 0 | 0 | 0 | 0 | 0 | 0 | 0 | 0 | 0 | 1 | 0 | 0 | 0.5 | 0.5 | 1 |
| PP_0168 | Putative surface adhesion protein | 199,318 | T | G | Synonymous_Coding | acT/acG | T1608 | 0 | 0 | 0 | 1 | 0 | 1 | 0 | 0 | 0 | 0 | 0 | 0 | 0 | 0 | 0 | 1 | 1 | 0 | 1 | 0.5 | 0.5 |
| PP_0168 | Putative surface adhesion protein | 199,480 | T | C | Synonymous_Coding | ggT/ggC | G1662 | 0 | 0 | 0 | 0 | 0 | 0 | 0 | 0 | 0 | 0 | 1 | 0 | 0 | 0 | 0 | 0 | 0 | 1 | 1 | 0.5 | 0.5 |
| PP_0168 | Putative surface adhesion protein | 199,550 | T | C | Synonymous_Coding | Ttg/Ctg | L1686 | 0 | 0 | 0 | 0 | 0 | 0 | 0 | 0 | 0 | 0 | 1 | 0 | 0 | 0 | 1 | 0 | 0 | 2 | 1 | 0.2222 | 0.2222 |
| PP_0168 | Putative surface adhesion protein | 199,744 | T | C | Synonymous_Coding | aaT/aaC | N1750 | 0 | 0 | 0 | 0 | 0 | 0 | 0 | 0 | 0 | 0 | 0 | 0 | 1 | 0 | 0 | 0 | 0 | 1 | 1 | 0.5 | 0.5 |
| PP_0168 | Putative surface adhesion protein | 199,822 | G | C | Synonymous_Coding | gcG/gcC | A1776 | 1 | 0 | 0 | 0 | 0 | 0 | 0 | 0 | 0 | 0 | 0 | 0 | 0 | 0 | 0 | 1 | 0 | 0 | 0.5 | 0.5 | 1 |
| PP_0168 | Putative surface adhesion protein | 200,113 | T | C | Synonymous_Coding | aaT/aaC | N1873 | 0 | 0 | 0 | 0 | 0 | 1 | 1 | 0 | 0 | 0 | 0 | 0 | 0 | 0 | 1 | 0 | 2 | 1 | 0.2222 | 0.5 | 0.5833 |
| PP_0168 | Putative surface adhesion protein | 200,164 | T | C | Synonymous_Coding | gaT/gaC | D1890 | 0 | 0 | 1 | 1 | 0 | 0 | 0 | 0 | 0 | 0 | 0 | 0 | 0 | 0 | 1 | 2 | 0 | 1 | 0.2222 | 0.5833 | 0.5 |
| PP_0168 | Putative surface adhesion protein | 200,338 | G | C | Synonymous_Coding | gtG/gtC | V1948 | 0 | 0 | 0 | 0 | 0 | 0 | 0 | 0 | 0 | 1 | 0 | 0 | 0 | 0 | 0 | 0 | 1 | 0 | 0.5 | 1 | 0.5 |
| PP_0168 | Putative surface adhesion protein | 200,473 | T | C | Synonymous_Coding | ggT/ggC | G1993 | 0 | 0 | 1 | 0 | 0 | 0 | 1 | 0 | 1 | 0 | 0 | 0 | 0 | 0 | 0 | 1 | 2 | 0 | 0.5833 | 0.5 | 0.2222 |
| PP_0168 | Putative surface adhesion protein | 201,007 | T | C | Synonymous_Coding | ggT/ggC | G2171 | 0 | 0 | 0 | 0 | 0 | 0 | 0 | 0 | 0 | 0 | 0 | 0 | 0 | 1 | 0 | 0 | 0 | 1 | 1 | 0.5 | 0.5 |
| PP_0168 | Putative surface adhesion protein | 201,043 | T | C | Synonymous_Coding | gaT/gaC | D2183 | 0 | 0 | 0 | 1 | 0 | 0 | 0 | 0 | 0 | 0 | 0 | 0 | 0 | 0 | 1 | 1 | 0 | 1 | 0.5 | 1 | 0.5 |
| PP_0168 | Putative surface adhesion protein | 201,100 | T | C | Synonymous_Coding | acT/acC | T2202 | 0 | 0 | 0 | 0 | 0 | 0 | 0 | 0 | 0 | 0 | 0 | 1 | 0 | 0 | 0 | 0 | 0 | 1 | 1 | 0.5 | 0.5 |
| PP_0168 | Putative surface adhesion protein | 201,175 | T | C | Synonymous_Coding | aaT/aaC | N2227 | 1 | 0 | 1 | 0 | 0 | 0 | 0 | 0 | 0 | 0 | 0 | 0 | 0 | 0 | 0 | 2 | 0 | 0 | 0.2222 | 0.2222 | 1 |
| PP_0168 | Putative surface adhesion protein | 201,622 | C | G | Synonymous_Coding | gtC/gtG | V2376 | 0 | 0 | 0 | 0 | 0 | 0 | 0 | 0 | 0 | 0 | 0 | 1 | 0 | 0 | 0 | 0 | 0 | 1 | 1 | 0.5 | 0.5 |
| PP_0168 | Putative surface adhesion protein | 201,652 | G | C | Synonymous_Coding | gtG/gtC | V2386 | 0 | 0 | 0 | 0 | 0 | 0 | 0 | 0 | 0 | 0 | 0 | 1 | 0 | 0 | 1 | 0 | 0 | 2 | 1 | 0.2222 | 0.2222 |
| PP_0168 | Putative surface adhesion protein | 201,685 | G | C | Synonymous_Coding | acG/acC | T2397 | 0 | 0 | 0 | 1 | 0 | 0 | 0 | 0 | 0 | 0 | 1 | 0 | 1 | 0 | 0 | 1 | 0 | 2 | 0.5 | 0.5833 | 0.2222 |
| PP_0168 | Putative surface adhesion protein | 201,715 | T | C | Synonymous_Coding | aaT/aaC | N2407 | 0 | 0 | 0 | 0 | 0 | 1 | 0 | 0 | 0 | 0 | 0 | 0 | 0 | 0 | 0 | 0 | 1 | 0 | 0.5 | 1 | 0.5 |
| PP_0168 | Putative surface adhesion protein | 201,798 | T | C | Nonsynonymous_Coding | gTa/gCa | V2435A | 0 | 0 | 0 | 0 | 0 | 0 | 0 | 0 | 0 | 0 | 0 | 1 | 0 | 0 | 0 | 0 | 0 | 1 | 1 | 0.5 | 0.5 |
| PP_0168 | Putative surface adhesion protein | 201,859 | G | C | Synonymous_Coding | gcG/gcC | A2455 | 0 | 0 | 0 | 0 | 0 | 0 | 0 | 0 | 1 | 0 | 0 | 1 | 0 | 1 | 0 | 0 | 1 | 2 | 0.5 | 0.2222 | 0.5833 |
| PP_0168 | Putative surface adhesion protein | 201,997 | T | C | Synonymous_Coding | acT/acC | T2501 | 0 | 0 | 0 | 0 | 0 | 0 | 0 | 0 | 0 | 0 | 0 | 1 | 0 | 0 | 0 | 0 | 0 | 1 | 1 | 0.5 | 0.5 |
| PP_0168 | Putative surface adhesion protein | 202,111 | G | A | Synonymous_Coding | gaG/gaA | E2539 | 0 | 1 | 0 | 0 | 0 | 0 | 0 | 0 | 0 | 0 | 0 | 0 | 0 | 0 | 0 | 1 | 0 | 0 | 0.5 | 0.5 | 1 |
| PP_0168 | Putative surface adhesion protein | 202,117 | T | C | Synonymous_Coding | agT/agC | S2541 | 0 | 1 | 0 | 0 | 0 | 0 | 0 | 0 | 0 | 0 | 0 | 0 | 0 | 0 | 0 | 1 | 0 | 0 | 0.5 | 0.5 | 1 |
| PP_0168 | Putative surface adhesion protein | 202,321 | T | C | Synonymous_Coding | ggT/ggC | G2609 | 0 | 0 | 1 | 0 | 0 | 0 | 0 | 0 | 0 | 0 | 0 | 0 | 0 | 0 | 0 | 1 | 0 | 0 | 0.5 | 0.5 | 1 |
| PP_0168 | Putative surface adhesion protein | 203,227 | C | G | Synonymous_Coding | acC/acG | T2911 | 0 | 0 | 0 | 0 | 0 | 1 | 0 | 0 | 0 | 0 | 0 | 0 | 0 | 0 | 0 | 0 | 1 | 0 | 0.5 | 1 | 0.5 |
| PP_0168 | Putative surface adhesion protein | 203,422 | T | C | Synonymous_Coding | gcT/gcC | A2976 | 0 | 0 | 0 | 0 | 1 | 0 | 0 | 0 | 1 | 1 | 0 | 0 | 1 | 0 | 0 | 1 | 2 | 1 | 0.5833 | 1 | 0.5833 |
| PP_0168 | Putative surface adhesion protein | 203,647 | T | C | Synonymous_Coding | acT/acC | T3051 | 0 | 0 | 1 | 0 | 0 | 0 | 0 | 0 | 0 | 0 | 0 | 0 | 0 | 0 | 0 | 1 | 0 | 0 | 0.5 | 0.5 | 1 |
| PP_0168 | Putative surface adhesion protein | 203,674 | T | C | Synonymous_Coding | gaT/gaC | D3060 | 0 | 1 | 0 | 0 | 0 | 0 | 0 | 0 | 0 | 0 | 1 | 0 | 0 | 0 | 0 | 1 | 0 | 1 | 0.5 | 1 | 0.5 |
| PP_0168 | Putative surface adhesion protein | 204,180 | A | C | Nonsynonymous_Coding | aAc/aCc | N3229T | 0 | 0 | 0 | 0 | 0 | 0 | 0 | 0 | 0 | 0 | 0 | 1 | 0 | 0 | 0 | 0 | 0 | 1 | 1 | 0.5 | 0.5 |
| PP_0168 | Putative surface adhesion protein | 204,532 | T | C | Synonymous_Coding | ggT/ggC | G3346 | 0 | 0 | 1 | 0 | 0 | 0 | 0 | 0 | 0 | 0 | 0 | 0 | 0 | 0 | 0 | 1 | 0 | 0 | 0.5 | 0.5 | 1 |
| PP_0168 | Putative surface adhesion protein | 205,120 | A | T | Synonymous_Coding | gcA/gcT | A3542 | 0 | 0 | 0 | 0 | 0 | 0 | 0 | 0 | 0 | 0 | 0 | 0 | 1 | 0 | 0 | 0 | 0 | 1 | 1 | 0.5 | 0.5 |
| PP_0168 | Putative surface adhesion protein | 205,327 | C | G | Synonymous_Coding | acC/acG | T3611 | 0 | 0 | 1 | 0 | 0 | 0 | 0 | 0 | 0 | 0 | 0 | 1 | 0 | 0 | 0 | 1 | 0 | 1 | 0.5 | 1 | 0.5 |
| PP_0168 | Putative surface adhesion protein | 205,672 | T | C | Synonymous_Coding | acT/acC | T3726 | 0 | 1 | 0 | 0 | 0 | 0 | 0 | 0 | 0 | 0 | 0 | 0 | 0 | 0 | 0 | 1 | 0 | 0 | 0.5 | 0.5 | 1 |

| | | | | | | | | | | | | | | | | | | | | | | | | | | | | |
|---|---|---|---|---|---|---|---|---|---|---|---|---|---|---|---|---|---|---|---|---|---|---|---|---|---|---|---|---|
| PP_0168 | Putative surface adhesion protein | 205,789 | C | T | Synonymous_Coding | ggC/ggT | G3765 | 0 | 1 | 0 | 0 | 0 | 0 | 0 | 0 | 0 | 0 | 0 | 0 | 0 | 0 | 0 | 1 | 0 | 0 | 0.5 | 0.5 | 1 |
| PP_0168 | Putative surface adhesion protein | 205,912 | G | C | Synonymous_Coding | acG/acC | T3806 | 0 | 0 | 1 | 0 | 0 | 0 | 0 | 0 | 0 | 0 | 0 | 0 | 0 | 0 | 0 | 1 | 0 | 0 | 0.5 | 0.5 | 1 |
| PP_0168 | Putative surface adhesion protein | 206,143 | A | G | Synonymous_Coding | caA/caG | Q3883 | 0 | 1 | 0 | 0 | 0 | 0 | 0 | 0 | 0 | 0 | 0 | 0 | 0 | 0 | 1 | 1 | 0 | 1 | 0.5 | 1 | 0.5 |
| PP_0168 | Putative surface adhesion protein | 206,311 | G | C | Synonymous_Coding | gtG/gtC | V3939 | 0 | 0 | 0 | 0 | 0 | 0 | 0 | 0 | 0 | 0 | 0 | 1 | 0 | 0 | 0 | 0 | 0 | 1 | 1 | 0.5 | 0.5 |
| PP_0168 | Putative surface adhesion protein | 206,440 | T | C | Synonymous_Coding | ggT/ggC | G3982 | 1 | 0 | 0 | 0 | 0 | 0 | 0 | 0 | 0 | 0 | 0 | 0 | 0 | 0 | 0 | 1 | 0 | 0 | 0.5 | 0.5 | 1 |
| PP_0168 | Putative surface adhesion protein | 206,461 | G | C | Synonymous_Coding | gtG/gtC | V3989 | 0 | 1 | 0 | 0 | 1 | 0 | 0 | 0 | 0 | 0 | 0 | 0 | 1 | 0 | 0 | 2 | 0 | 1 | 0.2222 | 0.5833 | 0.5 |
| PP_0168 | Putative surface adhesion protein | 206,830 | T | C | Synonymous_Coding | gtT/gtC | V4112 | 0 | 0 | 0 | 0 | 0 | 0 | 0 | 0 | 0 | 0 | 0 | 0 | 0 | 1 | 0 | 0 | 0 | 1 | 1 | 0.5 | 0.5 |
| PP_0168 | Putative surface adhesion protein | 206,966 | T | C | Synonymous_Coding | Ttg/Ctg | L4158 | 0 | 0 | 0 | 0 | 1 | 0 | 0 | 0 | 0 | 0 | 0 | 0 | 0 | 0 | 1 | 1 | 0 | 1 | 0.5 | 1 | 0.5 |
| PP_0168 | Putative surface adhesion protein | 207,133 | T | C | Synonymous_Coding | acT/acC | T4213 | 0 | 0 | 0 | 0 | 1 | 0 | 0 | 0 | 0 | 0 | 1 | 1 | 0 | 0 | 1 | 1 | 0 | 3 | 0.5 | 0.2857 | 0.08333 |
| PP_0168 | Putative surface adhesion protein | 207,136 | C | G | Synonymous_Coding | gtC/gtG | V4214 | 0 | 0 | 0 | 0 | 0 | 0 | 0 | 0 | 0 | 0 | 0 | 1 | 0 | 0 | 0 | 0 | 0 | 1 | 1 | 0.5 | 0.5 |
| PP_0168 | Putative surface adhesion protein | 207,235 | G | A | Synonymous_Coding | acG/acA | T4247 | 0 | 0 | 1 | 0 | 0 | 0 | 0 | 0 | 0 | 0 | 0 | 1 | 0 | 0 | 0 | 1 | 0 | 1 | 0.5 | 1 | 0.5 |
| PP_0168 | Putative surface adhesion protein | 207,370 | G | T | Synonymous_Coding | gtG/gtT | V4292 | 0 | 1 | 0 | 0 | 0 | 0 | 0 | 0 | 0 | 0 | 0 | 0 | 0 | 0 | 0 | 1 | 0 | 0 | 0.5 | 0.5 | 1 |
| PP_0168 | Putative surface adhesion protein | 207,796 | G | C | Synonymous_Coding | gtG/gtC | V4434 | 1 | 0 | 0 | 0 | 0 | 0 | 0 | 0 | 0 | 0 | 0 | 0 | 0 | 0 | 0 | 1 | 0 | 0 | 0.5 | 0.5 | 1 |
| PP_0168 | Putative surface adhesion protein | 208,015 | A | G | Synonymous_Coding | aaA/aaG | K4507 | 0 | 0 | 1 | 0 | 0 | 0 | 0 | 0 | 0 | 0 | 0 | 0 | 0 | 0 | 0 | 1 | 0 | 0 | 0.5 | 0.5 | 1 |
| PP_0168 | Putative surface adhesion protein | 208,027 | T | C | Synonymous_Coding | acT/acC | T4511 | 0 | 0 | 1 | 0 | 0 | 0 | 0 | 0 | 0 | 0 | 0 | 0 | 0 | 0 | 0 | 1 | 0 | 0 | 0.5 | 0.5 | 1 |
| PP_0168 | Putative surface adhesion protein | 208,030 | T | G | Synonymous_Coding | gtT/gtG | V4512 | 0 | 0 | 1 | 0 | 0 | 0 | 0 | 0 | 0 | 0 | 0 | 0 | 0 | 0 | 0 | 1 | 0 | 0 | 0.5 | 0.5 | 1 |
| PP_0168 | Putative surface adhesion protein | 208,150 | T | C | Synonymous_Coding | gtT/gtC | V4552 | 0 | 0 | 1 | 1 | 0 | 0 | 0 | 0 | 0 | 0 | 0 | 0 | 0 | 1 | 0 | 2 | 0 | 1 | 0.2222 | 0.5833 | 0.5 |
| PP_0168 | Putative surface adhesion protein | 208,399 | T | C | Synonymous_Coding | agT/agC | S4635 | 0 | 0 | 0 | 0 | 0 | 0 | 0 | 0 | 0 | 0 | 0 | 0 | 1 | 0 | 0 | 0 | 0 | 1 | 1 | 0.5 | 0.5 |
| PP_0168 | Putative surface adhesion protein | 208,456 | C | G | Synonymous_Coding | gtC/gtG | V4654 | 0 | 0 | 1 | 0 | 0 | 0 | 0 | 0 | 0 | 0 | 0 | 0 | 0 | 0 | 0 | 1 | 0 | 0 | 0.5 | 0.5 | 1 |
| PP_0168 | Putative surface adhesion protein | 208,564 | A | G | Synonymous_Coding | aaA/aaG | K4690 | 0 | 0 | 0 | 0 | 0 | 1 | 0 | 0 | 0 | 0 | 0 | 0 | 0 | 0 | 0 | 0 | 1 | 0 | 0.5 | 1 | 0.5 |
| PP_0168 | Putative surface adhesion protein | 208,675 | G | A | Synonymous_Coding | aaG/aaA | K4727 | 0 | 0 | 1 | 0 | 0 | 0 | 0 | 0 | 0 | 0 | 0 | 0 | 0 | 0 | 0 | 1 | 0 | 0 | 0.5 | 0.5 | 1 |
| PP_0168 | Putative surface adhesion protein | 209,404 | T | C | Synonymous_Coding | gcT/gcC | A4970 | 0 | 1 | 0 | 0 | 0 | 1 | 0 | 0 | 0 | 0 | 0 | 0 | 0 | 0 | 1 | 1 | 1 | 1 | 1 | 1 | 1 |
| PP_0168 | Putative surface adhesion protein | 209,638 | T | C | Synonymous_Coding | ggT/ggC | G5048 | 0 | 0 | 0 | 0 | 0 | 1 | 0 | 0 | 0 | 0 | 0 | 0 | 0 | 0 | 0 | 0 | 1 | 0 | 0.5 | 1 | 0.5 |
| PP_0168 | Putative surface adhesion protein | 209,782 | T | C | Synonymous_Coding | gcT/gcC | A5096 | 0 | 0 | 0 | 0 | 0 | 0 | 0 | 0 | 1 | 0 | 0 | 0 | 0 | 0 | 0 | 0 | 1 | 0 | 0.5 | 1 | 0.5 |
| PP_0168 | Putative surface adhesion protein | 209,942 | T | C | Synonymous_Coding | Ttg/Ctg | L5150 | 0 | 0 | 0 | 1 | 0 | 0 | 0 | 0 | 0 | 0 | 0 | 0 | 0 | 0 | 0 | 1 | 0 | 0 | 0.5 | 0.5 | 1 |
| PP_0168 | Putative surface adhesion protein | 210,088 | A | G | Synonymous_Coding | gaA/gaG | E5198 | 0 | 0 | 0 | 0 | 1 | 0 | 0 | 0 | 0 | 0 | 0 | 0 | 0 | 0 | 0 | 1 | 0 | 0 | 0.5 | 0.5 | 1 |
| PP_0168 | Putative surface adhesion protein | 210,121 | T | C | Synonymous_Coding | ggT/ggC | G5209 | 0 | 0 | 0 | 0 | 0 | 0 | 0 | 0 | 0 | 0 | 0 | 0 | 0 | 0 | 1 | 0 | 0 | 1 | 1 | 0.5 | 0.5 |
| PP_0168 | Putative surface adhesion protein | 210,271 | T | C | Synonymous_Coding | acT/acC | T5259 | 0 | 1 | 0 | 0 | 0 | 0 | 0 | 0 | 0 | 0 | 0 | 0 | 0 | 0 | 0 | 1 | 0 | 0 | 0.5 | 0.5 | 1 |
| PP_0168 | Putative surface adhesion protein | 210,691 | T | C | Synonymous_Coding | gcT/gcC | A5399 | 0 | 0 | 0 | 0 | 0 | 0 | 0 | 0 | 0 | 0 | 0 | 0 | 1 | 0 | 0 | 0 | 0 | 1 | 1 | 0.5 | 0.5 |
| PP_0168 | Putative surface adhesion protein | 210,721 | T | C | Synonymous_Coding | aaT/aaC | N5409 | 1 | 0 | 1 | 0 | 0 | 0 | 0 | 0 | 0 | 0 | 0 | 0 | 1 | 0 | 0 | 2 | 0 | 1 | 0.2222 | 0.5833 | 0.5 |
| PP_0168 | Putative surface adhesion protein | 211,417 | T | C | Synonymous_Coding | gcT/gcC | A5641 | 0 | 0 | 0 | 1 | 0 | 0 | 0 | 0 | 0 | 0 | 0 | 0 | 0 | 0 | 0 | 1 | 0 | 0 | 0.5 | 0.5 | 1 |
| PP_0168 | Putative surface adhesion protein | 211,669 | T | C | Synonymous_Coding | gaT/gaC | D5725 | 0 | 0 | 0 | 0 | 0 | 0 | 0 | 0 | 0 | 0 | 0 | 1 | 0 | 0 | 0 | 0 | 0 | 1 | 1 | 0.5 | 0.5 |
| PP_0168 | Putative surface adhesion protein | 212,026 | T | C | Synonymous_Coding | gcT/gcC | A5844 | 0 | 0 | 0 | 0 | 0 | 1 | 0 | 0 | 0 | 0 | 0 | 0 | 0 | 0 | 0 | 0 | 1 | 0 | 0.5 | 1 | 0.5 |
| PP_0168 | Putative surface adhesion protein | 212,056 | T | C | Synonymous_Coding | aaT/aaC | N5854 | 1 | 0 | 0 | 0 | 0 | 0 | 0 | 0 | 0 | 0 | 0 | 0 | 0 | 0 | 0 | 1 | 0 | 0 | 0.5 | 0.5 | 1 |
| PP_0168 | Putative surface adhesion protein | 212,080 | T | C | Synonymous_Coding | ggT/ggC | G5862 | 0 | 0 | 0 | 0 | 0 | 1 | 0 | 0 | 0 | 0 | 0 | 0 | 1 | 0 | 0 | 0 | 1 | 1 | 0.5 | 0.5 | 1 |
| PP_0168 | Putative surface adhesion protein | 212,134 | T | C | Synonymous_Coding | gaT/gaC | D5880 | 0 | 0 | 0 | 0 | 0 | 0 | 0 | 0 | 0 | 0 | 0 | 0 | 0 | 0 | 1 | 0 | 0 | 1 | 1 | 0.5 | 0.5 |
| PP_0168 | Putative surface adhesion protein | 212,302 | T | C | Synonymous_Coding | acT/acC | T5936 | 0 | 0 | 0 | 0 | 0 | 0 | 0 | 0 | 1 | 0 | 0 | 0 | 0 | 0 | 0 | 0 | 1 | 0 | 0.5 | 1 | 0.5 |
| PP_0168 | Putative surface adhesion protein | 212,311 | T | C | Synonymous_Coding | atT/atC | I5939 | 0 | 0 | 0 | 0 | 0 | 0 | 0 | 0 | 0 | 1 | 0 | 0 | 0 | 0 | 0 | 0 | 1 | 0 | 0.5 | 1 | 0.5 |
| PP_0168 | Putative surface adhesion protein | 212,581 | G | A | Synonymous_Coding | ccG/ccA | P6029 | 0 | 0 | 0 | 0 | 0 | 0 | 0 | 0 | 0 | 0 | 0 | 1 | 0 | 0 | 0 | 0 | 0 | 1 | 1 | 0.5 | 0.5 |
| PP_0168 | Putative surface adhesion protein | 212,617 | C | G | Synonymous_Coding | acC/acG | T6041 | 0 | 0 | 0 | 0 | 0 | 0 | 0 | 0 | 0 | 0 | 0 | 1 | 0 | 0 | 0 | 0 | 0 | 1 | 1 | 0.5 | 0.5 |
| PP_0168 | Putative surface adhesion protein | 212,861 | G | C | Nonsynonymous_Coding | Gag/Cag | E6123Q | 0 | 0 | 1 | 0 | 0 | 0 | 0 | 0 | 0 | 0 | 0 | 0 | 0 | 0 | 0 | 1 | 0 | 0 | 0.5 | 0.5 | 1 |
| PP_0168 | Putative surface adhesion protein | 212,941 | C | G | Synonymous_Coding | gtC/gtG | V6149 | 0 | 0 | 0 | 0 | 0 | 1 | 0 | 0 | 0 | 0 | 0 | 0 | 0 | 0 | 0 | 0 | 1 | 0 | 0.5 | 1 | 0.5 |
| PP_0168 | Putative surface adhesion protein | 213,028 | G | C | Synonymous_Coding | acG/acC | T6178 | 0 | 0 | 0 | 0 | 0 | 0 | 0 | 0 | 0 | 0 | 0 | 0 | 0 | 0 | 1 | 0 | 0 | 1 | 1 | 0.5 | 0.5 |
| PP_0168 | Putative surface adhesion protein | 213,415 | A | G | Synonymous_Coding | ccA/ccG | P6307 | 0 | 0 | 0 | 0 | 0 | 0 | 0 | 0 | 0 | 0 | 0 | 1 | 0 | 0 | 0 | 0 | 0 | 1 | 1 | 0.5 | 0.5 |
| PP_0168 | Putative surface adhesion protein | 213,619 | T | C | Synonymous_Coding | atT/atC | I6375 | 0 | 0 | 0 | 0 | 1 | 0 | 0 | 0 | 0 | 0 | 0 | 0 | 0 | 0 | 0 | 1 | 0 | 0 | 0.5 | 0.5 | 1 |
| PP_0168 | Putative surface adhesion protein | 213,874 | T | G | Synonymous_Coding | acT/acG | T6460 | 0 | 0 | 0 | 0 | 0 | 1 | 0 | 0 | 0 | 0 | 0 | 0 | 1 | 0 | 0 | 0 | 1 | 1 | 0.5 | 0.5 | 1 |
| PP_0168 | Putative surface adhesion protein | 213,918 | C | T | Nonsynonymous_Coding | cCg/cTg | P6475L | 0 | 0 | 0 | 0 | 0 | 0 | 0 | 0 | 0 | 0 | 0 | 0 | 1 | 0 | 0 | 0 | 0 | 1 | 1 | 0.5 | 0.5 |
| PP_0168 | Putative surface adhesion protein | 214,699 | T | C | Synonymous_Coding | gcT/gcC | A6735 | 0 | 0 | 0 | 0 | 0 | 0 | 0 | 0 | 0 | 0 | 0 | 0 | 0 | 0 | 1 | 0 | 0 | 1 | 1 | 0.5 | 0.5 |
| PP_0168 | Putative surface adhesion protein | 214,777 | G | T | Synonymous_Coding | gcG/gcT | A6761 | 0 | 0 | 0 | 1 | 0 | 0 | 0 | 0 | 0 | 0 | 0 | 0 | 0 | 0 | 0 | 1 | 0 | 0 | 0.5 | 0.5 | 1 |
| PP_0168 | Putative surface adhesion protein | 215,035 | T | C | Synonymous_Coding | acT/acC | T6847 | 0 | 0 | 0 | 0 | 1 | 0 | 0 | 0 | 0 | 0 | 0 | 0 | 0 | 0 | 1 | 1 | 0 | 1 | 0.5 | 1 | 0.5 |
| PP_0168 | Putative surface adhesion protein | 215,047 | C | T | Synonymous_Coding | aaC/aaT | N6851 | 0 | 0 | 0 | 1 | 0 | 0 | 0 | 0 | 0 | 0 | 0 | 0 | 0 | 0 | 0 | 1 | 0 | 0 | 0.5 | 0.5 | 1 |
| PP_0168 | Putative surface adhesion protein | 215,077 | T | C | Synonymous_Coding | gcT/gcC | A6861 | 0 | 0 | 0 | 0 | 0 | 0 | 0 | 0 | 0 | 0 | 0 | 0 | 0 | 0 | 1 | 0 | 0 | 1 | 1 | 0.5 | 0.5 |
| PP_0168 | Putative surface adhesion protein | 215,383 | A | G | Synonymous_Coding | gaA/gaG | E6963 | 0 | 0 | 0 | 0 | 1 | 0 | 0 | 0 | 0 | 0 | 0 | 1 | 1 | 0 | 0 | 1 | 0 | 2 | 0.5 | 0.5833 | 0.2222 |
| PP_0168 | Putative surface adhesion protein | 215,897 | C | T | Synonymous_Coding | Ctg/Ttg | L7135 | 0 | 0 | 0 | 0 | 0 | 0 | 0 | 0 | 0 | 0 | 0 | 0 | 0 | 0 | 1 | 0 | 0 | 1 | 1 | 0.5 | 0.5 |
| PP_0168 | Putative surface adhesion protein | 215,929 | T | C | Synonymous_Coding | acT/acC | T7145 | 1 | 0 | 0 | 0 | 0 | 0 | 0 | 0 | 0 | 0 | 0 | 0 | 0 | 0 | 0 | 1 | 0 | 0 | 0.5 | 0.5 | 1 |
| PP_0168 | Putative surface adhesion protein | 215,950 | A | G | Synonymous_Coding | aaA/aaG | K7152 | 0 | 0 | 0 | 1 | 0 | 0 | 0 | 0 | 0 | 0 | 0 | 0 | 0 | 0 | 0 | 1 | 0 | 0 | 0.5 | 0.5 | 1 |
| PP_0168 | Putative surface adhesion protein | 215,962 | T | C | Synonymous_Coding | acT/acC | T7156 | 0 | 0 | 1 | 0 | 0 | 0 | 0 | 0 | 0 | 0 | 0 | 0 | 0 | 0 | 0 | 1 | 0 | 0 | 0.5 | 0.5 | 1 |
| PP_0168 | Putative surface adhesion protein | 216,016 | T | C | Synonymous_Coding | aaT/aaC | N7174 | 0 | 0 | 0 | 0 | 0 | 0 | 0 | 0 | 1 | 0 | 0 | 0 | 0 | 0 | 0 | 0 | 1 | 0 | 0.5 | 1 | 0.5 |
| PP_0168 | Putative surface adhesion protein | 216,085 | T | C | Synonymous_Coding | gtT/gtC | V7197 | 0 | 0 | 1 | 0 | 0 | 0 | 0 | 0 | 0 | 0 | 0 | 0 | 0 | 0 | 0 | 1 | 0 | 0 | 0.5 | 0.5 | 1 |
| PP_0168 | Putative surface adhesion protein | 216,334 | T | C | Synonymous_Coding | agT/agC | S7280 | 0 | 0 | 0 | 0 | 0 | 1 | 0 | 0 | 0 | 0 | 0 | 0 | 0 | 0 | 0 | 0 | 1 | 0 | 0.5 | 1 | 0.5 |
| PP_0168 | Putative surface adhesion protein | 216,499 | A | G | Synonymous_Coding | aaA/aaG | K7335 | 0 | 0 | 0 | 0 | 0 | 0 | 0 | 0 | 0 | 0 | 0 | 0 | 1 | 0 | 0 | 0 | 0 | 1 | 1 | 0.5 | 0.5 |
| PP_0168 | Putative surface adhesion protein | 216,715 | G | A | Synonymous_Coding | ccG/ccA | P7407 | 0 | 1 | 0 | 0 | 0 | 0 | 0 | 0 | 0 | 0 | 0 | 0 | 0 | 0 | 0 | 1 | 0 | 0 | 0.5 | 0.5 | 1 |
| PP_0174 | Hypothetical protein | 226,226 | T | A | Nonsynonymous_Coding | cTg/cAg | L151Q | 0 | 0 | 0 | 0 | 0 | 0 | 0 | 0 | 1 | 0 | 0 | 0 | 0 | 0 | 0 | 0 | 1 | 0 | 0.5 | 1 | 0.5 |
| Anonymous | | 235,390 | G | A | Intergenic | | | 0 | 0 | 0 | 1 | 0 | 0 | 0 | 0 | 0 | 0 | 0 | 0 | 0 | 0 | 0 | 1 | 0 | 0 | 0.5 | 0.5 | 1 |
| PP_0188 | methyltransferase | 204,531 | G | C | Nonsynonymous_Coding | Gcc/Ccc | A276P | 1 | 0 | 0 | 0 | 0 | 0 | 0 | 0 | 0 | 0 | 0 | 0 | 0 | 0 | 0 | 1 | 0 | 0 | 0.5 | 0.5 | 1 |
| PP_0188 | methyltransferase | 240,536 | G | C | Synonymous_Coding | ggG/ggC | G277 | 1 | 0 | 0 | 0 | 0 | 0 | 0 | 0 | 0 | 0 | 0 | 0 | 0 | 0 | 0 | 1 | 0 | 0 | 0.5 | 0.5 | 1 |
| Anonymous | | 278,742 | T | TA | Intergenic | | | 1 | 1 | 1 | 1 | 1 | 1 | 1 | 1 | 1 | 1 | 1 | 1 | 1 | 1 | 1 | 5 | 5 | 5 | 1 | 1 | 1 |
| tauB-II | Taurine ABC transporter ATP-binding protein | 287,067 | G | C | Nonsynonymous_Coding | cCg/cGg | P19R | 0 | 0 | 0 | 0 | 0 | 0 | 0 | 1 | 0 | 0 | 0 | 0 | 0 | 0 | 0 | 0 | 1 | 0 | 0.5 | 1 | 0.5 |
| tauB-II | Taurine ABC transporter ATP-binding protein | 287,068 | G | C | Nonsynonymous_Coding | Ccg/Gcg | P19A | 0 | 0 | 0 | 0 | 0 | 0 | 0 | 1 | 0 | 0 | 0 | 0 | 0 | 0 | 0 | 0 | 1 | 0 | 0.5 | 1 | 0.5 |
| envZ | Osmolarity sensor protein EnvZ | 302,935 | G | C | Nonsynonymous_Coding | gGt/gCt | G367A | 0 | 0 | 0 | 0 | 0 | 0 | 0 | 0 | 0 | 0 | 0 | 1 | 0 | 0 | 0 | 0 | 0 | 1 | 1 | 0.5 | 0.5 |
| envZ | Osmolarity sensor protein EnvZ | 303,121 | G | GGCT | Codon change and codon insertion | ccg/cTGCcg | P433LP | 0 | 0 | 0 | 0 | 0 | 0 | 0 | 0 | 0 | 0 | 1 | 1 | 1 | 0 | 0 | 0 | 0 | 3 | 1 | 0.08333 | 0.08333 |
| Anonymous | | 308,738 | C | T | Intergenic | | | 0 | 1 | 0 | 0 | 1 | 0 | 0 | 0 | 0 | 0 | 0 | 0 | 0 | 0 | 0 | 2 | 0 | 0 | 0.2222 | 0.2222 | 1 |
| Anonymous | | 308,739 | A | G | Intergenic | | | 0 | 1 | 0 | 0 | 1 | 0 | 0 | 0 | 0 | 0 | 0 | 0 | 0 | 0 | 0 | 2 | 0 | 0 | 0.2222 | 0.2222 | 1 |
| Anonymous | | 336,298 | T | A | Intergenic | | | 0 | 0 | 0 | 0 | 0 | 0 | 0 | 1 | 0 | 0 | 0 | 0 | 0 | 0 | 0 | 0 | 1 | 0 | 0.5 | 1 | 0.5 |

| | | | | | | | | | | | | | | | | | | | | | | | | | | | | |
|---|---|---|---|---|---|---|---|---|---|---|---|---|---|---|---|---|---|---|---|---|---|---|---|---|---|---|---|---|
| Anonymous | | 353,067 | A | G | Intergenic | | | 1 | 1 | 1 | 1 | 1 | 1 | 1 | 1 | 1 | 1 | 1 | 1 | 1 | 1 | 1 | 5 | 5 | 5 | 1 | 1 | 1 |
| Anonymous | | 364,577 | T | A | Intergenic | | | 0 | 0 | 0 | 0 | 0 | 0 | 0 | 0 | 1 | 0 | 0 | 0 | 0 | 0 | 0 | 0 | 1 | 0 | 0.5 | 1 | 0.5 |
| thiL | Thiamin monophosphate kinase | 605,121 | G | A | Nonsynonymous_Coding | cGc/cAc | R145H | 0 | 0 | 0 | 0 | 0 | 0 | 0 | 1 | 0 | 0 | 0 | 0 | 0 | 0 | 0 | 0 | 1 | 0 | 0.5 | 1 | 0.5 |
| thiL | Thiamin monophosphate kinase | 605,126 | A | T | Nonsynonymous_Coding | Agc/Tgc | S147C | 0 | 0 | 0 | 0 | 0 | 0 | 0 | 1 | 0 | 0 | 0 | 0 | 0 | 0 | 0 | 0 | 1 | 0 | 0.5 | 1 | 0.5 |
| thiL | Thiamin monophosphate kinase | 605,131 | T | C | Synonymous_Coding | ggT/ggC | G148 | 0 | 0 | 0 | 0 | 0 | 0 | 0 | 1 | 0 | 0 | 0 | 0 | 0 | 0 | 0 | 0 | 1 | 0 | 0.5 | 1 | 0.5 |
| Anonymous | | 661,588 | C | T | Intergenic | | | 0 | 0 | 0 | 0 | 0 | 0 | 1 | 0 | 0 | 0 | 0 | 0 | 0 | 0 | 0 | 0 | 1 | 0 | 0.5 | 1 | 0.5 |
| Anonymous | | 786,604 | T | C | Intergenic | | | 0 | 0 | 0 | 0 | 0 | 0 | 0 | 0 | 0 | 0 | 0 | 0 | 0 | 0 | 1 | 0 | 0 | 1 | 1 | 0.5 | 0.5 |
| hemH | Ferrochelatase | 863,376 | A | T | Synonymous_Coding | gcA/gcT | A170 | 0 | 0 | 1 | 1 | 0 | 1 | 0 | 1 | 0 | 0 | 0 | 0 | 1 | 0 | 0 | 2 | 2 | 1 | 1 | 0.5833 | 0.5833 |
| Anonymous | | 975,287 | A | T | Intergenic | | | 0 | 0 | 0 | 0 | 0 | 1 | 0 | 0 | 0 | 0 | 0 | 0 | 0 | 0 | 0 | 0 | 1 | 0 | 0.5 | 1 | 0.5 |
| Anonymous | | 975,288 | A | T | Intergenic | | | 0 | 0 | 0 | 0 | 0 | 1 | 0 | 0 | 0 | 0 | 0 | 0 | 0 | 0 | 0 | 0 | 1 | 0 | 0.5 | 1 | 0.5 |
| PP_0861 | Outer membrane ferric siderophore receptor | 999,625 | C | G | Nonsynonymous_Coding | gGc/gCc | G182A | 0 | 0 | 0 | 0 | 0 | 1 | 0 | 0 | 0 | 0 | 0 | 0 | 0 | 0 | 0 | 0 | 1 | 0 | 0.5 | 1 | 0.5 |
| PP_0904 | Protein InaA | 1,044,757 | CAGCACGGCTGCCTGTATGGCA | C | Codon change and codon deletion | ggctgcctgtatggcaagcacgta/gta | GCLYGKHV156V | 0 | 0 | 0 | 0 | 0 | 0 | 0 | 0 | 0 | 0 | 0 | 0 | 0 | 1 | 0 | 0 | 0 | 1 | 1 | 0.5 | 0.5 |
| Anonymous | | 1,159,239 | T | G | Intergenic | | | 0 | 0 | 0 | 0 | 0 | 0 | 0 | 1 | 0 | 0 | 1 | 0 | 0 | 0 | 0 | 0 | 1 | 1 | 0.5 | 0.5 | 1 |
| Anonymous | | 1,184,536 | GC | G | Intergenic | | | 0 | 0 | 0 | 0 | 0 | 1 | 0 | 0 | 0 | 0 | 0 | 0 | 0 | 0 | 0 | 0 | 1 | 0 | 0.5 | 1 | 0.5 |
| xcpY | Type II secretion pathway protein XcpY | 1,202,854 | A | C | Nonsynonymous_Coding | cAg/cCg | Q228P | 0 | 0 | 0 | 0 | 0 | 1 | 0 | 0 | 0 | 0 | 0 | 0 | 0 | 0 | 0 | 0 | 1 | 0 | 0.5 | 1 | 0.5 |
| dtcD-II | C4-dicarboxylate transport transcriptional regulator | 1,221,046 | C | A | Nonsynonymous_Coding | gGc/gTc | G87V | 1 | 0 | 0 | 0 | 0 | 0 | 0 | 0 | 0 | 0 | 0 | 0 | 0 | 0 | 0 | 1 | 0 | 0 | 0.5 | 0.5 | 1 |
| PP_16SE | 16S ribosomal RNA | 1,326,938 | G | A | Intragenic | | | 1 | 0 | 0 | 0 | 0 | 1 | 1 | 1 | 1 | 1 | 0 | 0 | 0 | 0 | 0 | 1 | 5 | 0 | 0.02381 | 0.5 | 0.003968 |
| PP_1175 | Hypothetical protein | 1,350,104 | C | A | Stop lost_splice site region | taG/taT | *32Y | 0 | 0 | 0 | 0 | 0 | 1 | 0 | 0 | 0 | 0 | 0 | 0 | 0 | 0 | 0 | 0 | 1 | 0 | 0.5 | 1 | 0.5 |
| PP_1175 | Hypothetical protein | 1,350,119 | A | G | Synonymous_Coding | agT/agC | S27 | 0 | 0 | 0 | 1 | 0 | 0 | 0 | 0 | 0 | 0 | 0 | 0 | 0 | 0 | 0 | 1 | 0 | 0 | 0.5 | 0.5 | 1 |
| PP_1195 | Hypothetical protein | 1,370,775 | GGGTCGAAAA | G | Codon change and codon deletion | gaaaaggtcgag/gag | EKVE535E | 0 | 0 | 0 | 0 | 0 | 0 | 0 | 0 | 0 | 0 | 0 | 0 | 0 | 0 | 1 | 0 | 0 | 1 | 1 | 0.5 | 0.5 |
| PP_1256 | Alpha-ketoglutarate semialdehyde dehydrogenase | 1,435,822 | A | T | Nonsynonymous_Coding | Tgc/Agc | C48S | 0 | 0 | 0 | 0 | 0 | 0 | 0 | 0 | 1 | 0 | 0 | 0 | 0 | 0 | 0 | 0 | 1 | 0 | 0.5 | 1 | 0.5 |
| Anonymous | | 1,499,497 | T | TC | Intergenic | | | 0 | 0 | 1 | 0 | 1 | 0 | 1 | 0 | 0 | 1 | 0 | 0 | 1 | 1 | 0 | 2 | 2 | 2 | 1 | 1 | 1 |
| PP_1325 | Putative lipoprotein | 1,511,070 | T | A | Nonsynonymous_Coding | cAg/cTg | Q166L | 0 | 0 | 0 | 0 | 0 | 0 | 0 | 0 | 0 | 0 | 0 | 0 | 1 | 0 | 0 | 0 | 0 | 1 | 1 | 0.5 | 0.5 |
| ftsZ/rna60/PP_mr19 | Cell division protein FtsZ/nucleotide motif:Rfam:RF02243 | 1,503,160 | C | T | Nonsynonymous_Coding | Cgt/Tgt | R395C | 0 | 0 | 0 | 0 | 0 | 0 | 1 | 0 | 0 | 0 | 1 | 1 | 1 | 1 | 1 | 0 | 1 | 5 | 0.5 | 0.003968 | 0.02381 |
| ftsZ/rna60/PP_1344 | Cell division protein FtsZ/nucleotide motif:Rfam:RF02243 | 1,530,173 | A | C | Stop lost_Splice site region | tAa/tCa | *399S | 1 | 1 | 1 | 1 | 1 | 1 | 1 | 1 | 1 | 1 | 0 | 0 | 0 | 0 | 0 | 5 | 5 | 0 | 1 | 0.003968 | 0.003968 |
| PP_1344 | Hypothetical protein | 1,531,780 | G | T | Non synonymous_Coding | Ccc/Acc | P36T | 0 | 0 | 0 | 0 | 0 | 0 | 0 | 0 | 0 | 0 | 0 | 0 | 1 | 0 | 0 | 0 | 0 | 1 | 1 | 0.5 | 0.5 |
| Anonymous | | 1,617,777 | C | T | Intergenic | | | 0 | 0 | 0 | 0 | 0 | 0 | 0 | 0 | 0 | 0 | 1 | 0 | 0 | 0 | 0 | 0 | 0 | 1 | 1 | 0.5 | 0.5 |
| Anonymous | | 1,617,782 | C | G | Intergenic | | | 0 | 0 | 0 | 0 | 0 | 0 | 0 | 0 | 0 | 0 | 1 | 0 | 0 | 0 | 0 | 0 | 0 | 1 | 1 | 0.5 | 0.5 |
| Anonymous | | 1,777,419 | G | A | Intergenic | | | 0 | 1 | 0 | 0 | 1 | 1 | 1 | 0 | 1 | 1 | 1 | 1 | 0 | 0 | 1 | 2 | 4 | 3 | 0.2857 | 0.6032 | 0.5833 |
| Anonymous | | 1,777,422 | C | A | Intergenic | | | 0 | 1 | 0 | 0 | 1 | 1 | 1 | 0 | 1 | 1 | 1 | 1 | 0 | 0 | 0 | 2 | 4 | 2 | 0.2857 | 1 | 0.2857 |
| Anonymous | | 1,777,428 | G | A | Intergenic | | | 0 | 1 | 0 | 0 | 1 | 1 | 1 | 0 | 1 | 1 | 1 | 0 | 0 | 0 | 0 | 2 | 4 | 1 | 0.2857 | 0.5833 | 0.1071 |
| Anonymous | | 1,780,067 | C | T | Intergenic | | | 0 | 1 | 0 | 0 | 0 | 1 | 1 | 0 | 0 | 0 | 0 | 0 | 1 | 0 | 0 | 1 | 2 | 1 | 0.5833 | 1 | 0.5833 |
| Anonymous | | 1,838,142 | G | A | Intergenic | | | 0 | 0 | 0 | 0 | 0 | 0 | 0 | 0 | 1 | 0 | 0 | 0 | 0 | 0 | 0 | 0 | 1 | 0 | 0.5 | 1 | 0.5 |
| gacS | Sensor protein GacS | 1,843,091 | A | C | Nonsynonymous_Coding | gTg/gGg | V568G | 1 | 1 | 1 | 0 | 1 | 1 | 1 | 1 | 1 | 1 | 0 | 0 | 0 | 0 | 1 | 4 | 5 | 1 | 1 | 0.1071 | 0.02381 |
| gacS | Sensor protein GacS | 1,843,901 | C | T | Nonsynonymous_Coding | cGc/cAc | R298H | | | | | | | | | | | | | | | | 0 | 0 | 0 | 1 | 1 | 1 |
| gacS | Sensor protein GacS | 1,844,166 | T | A | Stop gained | Aag/Tag | K210* | 0 | 0 | 0 | 0 | 0 | 0 | 1 | 0 | 0 | 0 | 1 | 1 | 1 | 1 | 0 | 0 | 1 | 4 | 0.5 | 0.02381 | 0.1071 |
| PP_1666 | Hypothetical protein | 1,863,240 | T | A | Nonsynonymous_Coding | cTg/cAg | L309Q | 0 | 0 | 0 | 0 | 0 | 0 | 0 | 0 | 0 | 0 | 0 | 0 | 1 | 0 | 0 | 0 | 0 | 1 | 1 | 0.5 | 0.5 |
| PP_1666 | Hypothetical protein | 1,863,278 | G | C | Nonsynonymous_Coding | Gcg/Ccg | A322P | 0 | 0 | 1 | 0 | 0 | 0 | 0 | 0 | 0 | 0 | 0 | 0 | 0 | 0 | 0 | 1 | 0 | 0 | 0.5 | 0.5 | 1 |
| PP_1703 | Assimilatory nitrate reductase/sulfite reductase | 1,901,761 | A | T | Nonsynonymous_Coding | gAc/gTc | D787V | 0 | 0 | 0 | 0 | 0 | 0 | 0 | 1 | 0 | 0 | 0 | 0 | 0 | 0 | 0 | 0 | 1 | 0 | 0.5 | 1 | 0.5 |
| Anonymous | | 2,051,065 | T | A | Intergenic | | | 1 | 0 | 0 | 0 | 0 | 0 | 1 | 1 | 1 | 0 | 0 | 0 | 0 | 0 | 0 | 1 | 3 | 0 | 0.2857 | 0.5 | 0.08333 |
| Anonymous | | 2,063,185 | C | G | Intergenic | | | 0 | 0 | 0 | 0 | 0 | 1 | 0 | 0 | 0 | 0 | 0 | 0 | 0 | 0 | 0 | 0 | 1 | 0 | 0.5 | 1 | 0.5 |
| Anonymous | | 2,068,151 | A | T | Intergenic | | | 0 | 0 | 0 | 0 | 0 | 0 | 0 | 0 | 1 | 0 | 0 | 0 | 0 | 0 | 0 | 0 | 1 | 0 | 0.5 | 1 | 0.5 |
| Anonymous | | 2,068,152 | A | T | Intergenic | | | 0 | 0 | 0 | 0 | 0 | 0 | 0 | 0 | 1 | 0 | 0 | 0 | 0 | 0 | 0 | 0 | 1 | 0 | 0.5 | 1 | 0.5 |
| Anonymous | | 2,069,557 | T | C | Intergenic | | | 1 | 0 | 0 | 0 | 0 | 1 | 1 | 1 | 0 | 1 | 1 | 1 | 0 | 0 | 0 | 1 | 4 | 2 | 0.1071 | 0.5833 | 0.2857 |
| Anonymous | | 2,087,422 | C | T | Intergenic | | | 0 | 0 | 0 | 0 | 0 | 0 | 1 | 0 | 0 | 0 | 0 | 0 | 0 | 1 | 0 | 0 | 1 | 1 | 0.5 | 0.5 | 1 |
| Anonymous | | 2,087,428 | G | A | Intergenic | | | 1 | 0 | 0 | 0 | 0 | 0 | 1 | 0 | 0 | 1 | 1 | 0 | 1 | 0 | 0 | 1 | 2 | 2 | 0.5833 | 0.5833 | 1 |
| Anonymous | | 2,087,439 | G | A | Intergenic | | | 0 | 0 | 0 | 0 | 0 | 0 | 1 | 0 | 0 | 0 | 0 | 0 | 0 | 0 | 0 | 0 | 1 | 0 | 0.5 | 1 | 0.5 |
| Anonymous | | 2,087,463 | G | A | Intergenic | | | 0 | 0 | 0 | 0 | 0 | 0 | 1 | 0 | 0 | 0 | 0 | 0 | 0 | 0 | 0 | 0 | 1 | 0 | 0.5 | 1 | 0.5 |
| Anonymous | | 2,087,635 | T | C | Intergenic | | | 0 | 0 | 0 | 0 | 1 | 1 | 0 | 1 | 0 | 0 | 0 | 0 | 1 | 0 | 0 | 1 | 2 | 1 | 0.5833 | 1 | 0.5833 |
| PP_5491 | SEC-C motif domain-containing protein | 2,187,722 | GT | G | Intergenic | | | 0 | 0 | 0 | 0 | 0 | 0 | 1 | 0 | 0 | 0 | 0 | 0 | 0 | 0 | 0 | 0 | 1 | 0 | 0.5 | 1 | 0.5 |
| Anonymous | | 2,256,267 | C | T | Intergenic | | | 0 | 0 | 0 | 0 | 0 | 0 | 0 | 0 | 0 | 0 | 0 | 0 | 1 | 0 | 0 | 0 | 0 | 1 | 1 | 0.5 | 0.5 |
| Anonymous | | 2,256,290 | C | T | Intergenic | | | 0 | 0 | 0 | 0 | 0 | 0 | 0 | 0 | 0 | 0 | 0 | 0 | 1 | 0 | 0 | 0 | 0 | 1 | 1 | 0.5 | 0.5 |
| PP_2068 | Multidrug MFS transporter membrane fusion | 2,354,379 | T | G | Nonsynonymous_Coding | aaA/aaC | K51N | 0 | 0 | 0 | 1 | 0 | 0 | 0 | 0 | 0 | 0 | 0 | 0 | 0 | 0 | 0 | 1 | 0 | 0 | 0.5 | 0.5 | 1 |
| Anonymous | | 2,414,802 | C | G | Intergenic | | | 0 | 0 | 0 | 0 | 0 | 0 | 0 | 0 | 0 | 0 | 0 | 0 | 1 | 0 | 0 | 0 | 0 | 1 | 1 | 0.5 | 0.5 |
| Anonymous | | 2,550,588 | A | G | Intergenic | | | 0 | 0 | 0 | 0 | 0 | 0 | 0 | 0 | 0 | 1 | 0 | 0 | 0 | 0 | 0 | 0 | 1 | 0 | 0.5 | 1 | 0.5 |
| PP_2397 | EF hand domain-containing protein/Ca2+ binding motif | 2,742,917 | C | T | Nonsynonymous_Coding | Cac/Tac | H5Y | 0 | 0 | 0 | 0 | 0 | 1 | 0 | 0 | 0 | 0 | 0 | 0 | 0 | 0 | 0 | 0 | 1 | 0 | 0.5 | 1 | 0.5 |
| PP_2397 | EF hand domain-containing protein/Ca2+ binding motif | 2,742,918 | A | G | Nonsynonymous_Coding | cAc/cGc | H5R | 0 | 0 | 0 | 0 | 0 | 1 | 0 | 0 | 0 | 0 | 0 | 0 | 0 | 0 | 0 | 0 | 1 | 0 | 0.5 | 1 | 0.5 |
| PP_2478 | Isoquinoline 1-oxidoreductase subunit beta | 2,824,518 | A | C | Nonsynonymous_Coding | gAg/gCg | E44A | 0 | 0 | 0 | 0 | 1 | 0 | 0 | 0 | 0 | 0 | 0 | 0 | 0 | 0 | 0 | 1 | 0 | 0 | 0.5 | 0.5 | 1 |
| Anonymous | | 2,935,858 | C | G | Intergenic | | | 0 | 0 | 0 | 0 | 0 | 0 | 0 | 0 | 0 | 0 | 0 | 0 | 1 | 0 | 0 | 0 | 0 | 1 | 1 | 0.5 | 0.5 |
| PP_2638 | Cellulose synthase operon protein C | 3,021,030 | A | T | Nonsynonymous_Coding | cAg/cTg | Q523L | 0 | 0 | 1 | 0 | 0 | 0 | 0 | 1 | 0 | 0 | 1 | 0 | 0 | 1 | 0 | 1 | 1 | 2 | 1 | 0.5833 | 0.5833 |
| pp_2757 | ABC-type transport system/sugar-binding protein | 3,141,492 | GT | G | Frame shift | gtc/ | V137 | 0 | 0 | 0 | 0 | 0 | 0 | 0 | 0 | 0 | 0 | 0 | 1 | 0 | 0 | 0 | 0 | 0 | 1 | 1 | 0.5 | 0.5 |

| | | | | | | | | | | | | | | | | | | | | | | | | | | | | |
|---|---|---|---|---|---|---|---|---|---|---|---|---|---|---|---|---|---|---|---|---|---|---|---|---|---|---|---|---|
| PP_2758 | Periplasmic binding protein domain/ribose ABC transpoter substrate-binding protein | 3,142,272 | C | T | Nonsynonymous_Coding | gCg/gTg | A74V | 0 | 0 | 0 | 0 | 0 | 0 | 0 | 0 | 0 | 0 | 1 | 1 | 1 | 1 | 0 | 0 | 0 | 4 | 1 | 0.02381 | 0.02381 |
| PP_3045 | ATP-dependent protease ClpP | 3,430,349 | G | C | Synonymous_Coding | tcG/tcC | S189 | 0 | 0 | 0 | 0 | 0 | 0 | 0 | 0 | 0 | 0 | 0 | 0 | 1 | 0 | 0 | 0 | 0 | 1 | 1 | 0.5 | 0.5 |
| PP_3221 | ABC transporter permease/nickel ABC transporter | 3,656,539 | T | G | Synonymous_Coding | ccA/ccC | P253 | 0 | 0 | 0 | 0 | 0 | 0 | 0 | 0 | 0 | 0 | 1 | 0 | 0 | 0 | 0 | 0 | 0 | 1 | 1 | 0.5 | 0.5 |
| PP_5743/PP_3334 | TonB-dependent receptor protein/pseudo gene | 3,774,247 | C | G | Intragenic | | | 1 | 0 | 0 | 0 | 0 | 1 | 0 | 0 | 0 | 0 | 0 | 0 | 0 | 0 | 0 | 1 | 1 | 0 | 1 | 0.5 | 0.5 |
| oprN | Multidrug RND transporter outer membrane protein | 3,881,829 | A | T | Nonsynonymous_Coding | cAg/cTg | Q163L | 0 | 1 | 0 | 0 | 0 | 0 | 0 | 0 | 0 | 0 | 0 | 0 | 0 | 0 | 0 | 1 | 0 | 0 | 0.5 | 0.5 | 1 |
| Anonymous | | 4,022,306 | G | C | Intergenic | | | 0 | 0 | 0 | 1 | 0 | 1 | 1 | 1 | 1 | 0 | 1 | 0 | 1 | 1 | 0 | 1 | 4 | 3 | 0.1071 | 0.2857 | 0.5833 |
| Anonymous | | 4,022,307 | G | C | Intergenic | | | 0 | 0 | 0 | 1 | 0 | 1 | 1 | 1 | 1 | 0 | 1 | 0 | 1 | 1 | 0 | 1 | 4 | 3 | 0.1071 | 0.2857 | 0.5833 |
| Anonymous | | 4,022,806 | C | G | Intergenic | | | 1 | 0 | 0 | 0 | 0 | 1 | 0 | 1 | 0 | 0 | 0 | 0 | 1 | 1 | 0 | 1 | 2 | 2 | 0.5833 | 0.5833 | 1 |
| Anonymous | | 4,022,807 | C | G | Intergenic | | | 1 | 0 | 0 | 0 | 0 | 1 | 0 | 1 | 0 | 0 | 1 | 1 | 1 | 1 | 0 | 1 | 2 | 4 | 0.5833 | 0.1071 | 0.2857 |
| PP_3563 | Hypothetical protein | 4,042,322 | T | G | Nonsynonymous_Coding | cTc/cGc | L137R | 0 | 0 | 0 | 0 | 0 | 0 | 1 | 0 | 0 | 0 | 0 | 0 | 0 | 0 | 0 | 0 | 1 | 0 | 0.5 | 1 | 0.5 |
| PP_3573 | Monooxygenase/flavin-dependent oxidoreductase | 4,053,476 | A | T | Nonsynonymous_Coding | cAg/cTg | Q17L | 0 | 1 | 0 | 0 | 1 | 0 | 0 | 1 | 0 | 0 | 0 | 0 | 1 | 0 | 0 | 2 | 1 | 1 | 0.5833 | 0.5833 | 1 |
| Anonymous | | 4,293,218 | G | C | Intergenic | | | 0 | 0 | 0 | 0 | 0 | 0 | 0 | 0 | 0 | 0 | 1 | 0 | 0 | 0 | 0 | 0 | 0 | 1 | 1 | 0.5 | 0.5 |
| yrpB | 2-Nitropropane dioxugenase/dixygenases related to 2-Nitropropane dioxugenase | 4,353,296 | C | G | Nonsynonymous_Coding | cCg/cGg | P141R | 0 | 0 | 0 | 0 | 0 | 0 | 0 | 1 | 0 | 0 | 1 | 0 | 0 | 0 | 0 | 0 | 1 | 1 | 0.5 | 0.5 | 1 |
| clpA | ATP-dependent serine protease | 4,518,633 | C | T | Nonsynonymous_Coding | gGc/gAc | G296D | 0 | 0 | 0 | 0 | 0 | 0 | 0 | 0 | 0 | 0 | 0 | 0 | 0 | 0 | 1 | 0 | 0 | 1 | 1 | 0.5 | 0.5 |
| idh | Isocitrate dehydrogenase | 4,522,716 | G | A | Nonsynonymous_Coding | Gca/Aca | A189T | 0 | 0 | 1 | 0 | 0 | 0 | 0 | 0 | 0 | 0 | 0 | 0 | 0 | 0 | 0 | 1 | 0 | 0 | 0.5 | 0.5 | 1 |
| malQ | Biosynthesis and degradation of polysaccharides, 4-alpha-glucanotransferase | 4,568,619 | C | G | Synonymous_Coding | gcC/gcG | A183 | 1 | 0 | 0 | 0 | 0 | 0 | 0 | 0 | 0 | 0 | 0 | 0 | 0 | 0 | 0 | 1 | 0 | 0 | 0.5 | 0.5 | 1 |
| mccB | Methylcrotonyl-CoA carboxylase biotin-containing subunit beta | 4,990,049 | T | C | Nonsynonymous_Coding | gTc/gCc | V244A | 0 | 0 | 0 | 0 | 0 | 0 | 0 | 0 | 0 | 0 | 1 | 0 | 0 | 0 | 0 | 0 | 0 | 1 | 1 | 0.5 | 0.5 |
| uvrY | DBarA/UvrY two-componenent | 4,635,590 | T | C | Nonsynonymous_Coding | Aaa/Gaa | K56E | 1 | 1 | 1 | 1 | 1 | 1 | 1 | 1 | 1 | 1 | 0 | 0 | 0 | 0 | 1 | 5 | 5 | 1 | 1 | 0.02381 | 0.02381 |
| PP_5662 | Protein of unknown function | 4,741,229 | A | C | Intergenic | | | 1 | 1 | 1 | 0 | 1 | 1 | 1 | 1 | 1 | 0 | 1 | 1 | 1 | 1 | 1 | 4 | 4 | 5 | 1 | 1 | 1 |
| PP_5662 | Protein of unknown function | 4,741,231 | C | G | Intergenic | | | 1 | 1 | 1 | 0 | 1 | 1 | 1 | 1 | 0 | 0 | 1 | 1 | 1 | 1 | 1 | 4 | 3 | 5 | 0.5833 | 1 | 0.2222 |
| PP_5662 | Protein of unknown function | 4,741,234 | C | T | Intergenic | | | 1 | 1 | 1 | 0 | 1 | 1 | 1 | 1 | 1 | 0 | 1 | 1 | 1 | 1 | 1 | 4 | 4 | 5 | 1 | 1 | 1 |
| PP_5662 | Protein of unknown function | 4,741,236 | G | T | Intergenic | | | 1 | 1 | 1 | 0 | 1 | 1 | 1 | 1 | 1 | 0 | 1 | 1 | 1 | 1 | 1 | 4 | 4 | 5 | 1 | 1 | 1 |
| PP_5662 | Protein of unknown function | 4,741,237 | C | G | Intergenic | | | 1 | 1 | 1 | 0 | 1 | 1 | 1 | 1 | 1 | 0 | 1 | 1 | 1 | 1 | 1 | 4 | 4 | 5 | 1 | 1 | 1 |
| PP_5662 | Protein of unknown function | 4,741,239 | A | G | Intergenic | | | 1 | 1 | 1 | 0 | 1 | 1 | 1 | 1 | 1 | 0 | 1 | 1 | 1 | 1 | 1 | 4 | 4 | 5 | 1 | 1 | 1 |
| PP_5662 | Protein of unknown function | 4,741,243 | G | T | Intergenic | | | 1 | 1 | 1 | 0 | 1 | 1 | 1 | 1 | 1 | 0 | 1 | 1 | 1 | 1 | 1 | 4 | 4 | 5 | 1 | 1 | 1 |
| PP_5662 | Protein of unknown function | 4,741,244 | C | G | Intergenic | | | 1 | 1 | 1 | 0 | 1 | 1 | 1 | 1 | 1 | 0 | 1 | 1 | 1 | 1 | 1 | 4 | 4 | 5 | 1 | 1 | 1 |
| PP_5662 | Protein of unknown function | 4,741,245 | A | C | Intergenic | | | 1 | 1 | 1 | 0 | 1 | 1 | 1 | 1 | 1 | 0 | 1 | 1 | 1 | 1 | 1 | 4 | 4 | 5 | 1 | 1 | 1 |
| PP_5662 | Protein of unknown function | 4,741,255 | C | T | Intergenic | | | 1 | 1 | 1 | 0 | 1 | 1 | 1 | 1 | 1 | 0 | 1 | 1 | 1 | 1 | 1 | 4 | 4 | 5 | 1 | 1 | 1 |
| PP_5662 | Protein of unknown function | 4,741,257 | A | C | Intergenic | | | 1 | 1 | 1 | 0 | 1 | 1 | 1 | 1 | 1 | 0 | 1 | 1 | 1 | 1 | 1 | 4 | 4 | 5 | 1 | 1 | 1 |
| Anonymous | | 4,741,260 | C | G | Intergenic | | | 1 | 1 | 1 | 0 | 1 | 1 | 1 | 1 | 1 | 0 | 1 | 1 | 1 | 1 | 1 | 4 | 4 | 5 | 1 | 1 | 1 |
| Anonymous | | 4,741,261 | A | C | Intergenic | | | 1 | 1 | 1 | 0 | 1 | 1 | 1 | 1 | 1 | 0 | 1 | 1 | 1 | 1 | 1 | 4 | 4 | 5 | 1 | 1 | 1 |
| Anonymous | | 4,741,263 | A | C | Intergenic | | | 1 | 1 | 1 | 0 | 1 | 1 | 1 | 1 | 1 | 0 | 1 | 1 | 1 | 1 | 1 | 4 | 4 | 5 | 1 | 1 | 1 |
| Anonymous | | 4,741,272 | A | G | Intergenic | | | 1 | 1 | 1 | 0 | 1 | 1 | 1 | 1 | 1 | 0 | 1 | 1 | 1 | 1 | 1 | 4 | 4 | 5 | 1 | 1 | 1 |
| Anonymous | | 4,741,274 | A | C | Intergenic | | | 1 | 1 | 1 | 0 | 1 | 1 | 1 | 1 | 1 | 0 | 1 | 1 | 1 | 1 | 1 | 4 | 4 | 5 | 1 | 1 | 1 |
| Anonymous | | 4,741,276 | G | C | Intergenic | | | 1 | 1 | 1 | 0 | 1 | 1 | 1 | 1 | 1 | 0 | 1 | 1 | 1 | 1 | 1 | 4 | 4 | 5 | 1 | 1 | 1 |
| Anonymous | | 4,741,278 | C | T | Intergenic | | | 1 | 1 | 1 | 0 | 1 | 1 | 1 | 1 | 1 | 0 | 1 | 1 | 1 | 1 | 1 | 4 | 4 | 5 | 1 | 1 | 1 |
| Anonymous | | 4,741,283 | C | CT | Intergenic | | | 1 | 1 | 1 | 0 | 1 | 1 | 1 | 1 | 1 | 0 | 1 | 1 | 1 | 1 | 1 | 4 | 4 | 5 | 1 | 1 | 1 |
| Anonymous | | 4,741,289 | C | G | Intergenic | | | 1 | 1 | 1 | 0 | 1 | 1 | 1 | 1 | 1 | 0 | 1 | 1 | 1 | 1 | 1 | 4 | 4 | 5 | 1 | 1 | 1 |
| Anonymous | | 4,741,292 | C | T | Intergenic | | | 1 | 1 | 1 | 0 | 1 | 1 | 1 | 1 | 1 | 0 | 1 | 1 | 1 | 1 | 1 | 4 | 4 | 5 | 1 | 1 | 1 |
| Anonymous | | 4,741,294 | C | G | Intergenic | | | 1 | 1 | 1 | 0 | 1 | 1 | 1 | 1 | 1 | 0 | 1 | 1 | 1 | 1 | 1 | 4 | 4 | 5 | 1 | 1 | 1 |
| Anonymous | | 4,741,296 | A | C | Intergenic | | | 1 | 1 | 1 | 0 | 1 | 1 | 1 | 1 | 1 | 0 | 1 | 1 | 1 | 1 | 1 | 4 | 4 | 5 | 1 | 1 | 1 |
| Anonymous | | 4,741,298 | C | T | Intergenic | | | 1 | 1 | 1 | 0 | 1 | 1 | 1 | 1 | 1 | 0 | 1 | 1 | 1 | 1 | 1 | 4 | 4 | 5 | 1 | 1 | 1 |
| Anonymous | | 4,808,166 | GAGCGCCGCCCGCGA | G | Intergenic | | | 0 | 0 | 0 | 0 | 0 | 0 | 0 | 1 | 0 | 0 | 0 | 0 | 0 | 0 | 0 | 0 | 1 | 0 | 0.5 | 1 | 0.5 |
| Anonymous | | 4,861,127 | A | G | Intergenic | | | 0 | 0 | 0 | 0 | 0 | 0 | 0 | 0 | 0 | 0 | 0 | 0 | 1 | 0 | 0 | 0 | 0 | 1 | 1 | 0.5 | 0.5 |
| Anonymous | | 4,861,219 | C | T | Intergenic | | | 0 | 0 | 0 | 0 | 0 | 1 | 1 | 0 | 0 | 0 | 1 | 0 | 0 | 1 | 0 | 0 | 2 | 2 | 0.2222 | 0.2222 | 1 |
| ccmE | Cytochrome c-type biogenesis protein CcmE | 4,914,726 | T | A | Nonsynonymous_Coding | cAg/cTg | Q142L | 0 | 0 | 0 | 0 | 0 | 0 | 0 | 0 | 0 | 0 | 1 | 0 | 0 | 0 | 0 | 0 | 0 | 1 | 1 | 0.5 | 0.5 |
| Anonymous | | 4,945,198 | C | G | Intergenic | | | 0 | 0 | 1 | 0 | 0 | 1 | 0 | 0 | 0 | 0 | 1 | 0 | 0 | 0 | 1 | 1 | 1 | 2 | 1 | 0.5833 | 0.5833 |
| Anonymous | | 4,945,201 | C | G | Intergenic | | | 0 | 0 | 1 | 0 | 0 | 1 | 0 | 0 | 0 | 0 | 1 | 0 | 0 | 0 | 0 | 1 | 1 | 1 | 1 | 1 | 1 |
| Anonymous | | 4,945,389 | G | C | Intergenic | | | 1 | 0 | 0 | 0 | 0 | 0 | 1 | 0 | 0 | 0 | 0 | 1 | 0 | 0 | 0 | 1 | 1 | 1 | 1 | 1 | 1 |
| Anonymous | | 4,945,392 | G | C | Intergenic | | | 1 | 0 | 0 | 0 | 0 | 0 | 1 | 0 | 0 | 0 | 0 | 1 | 0 | 0 | 0 | 1 | 1 | 1 | 1 | 1 | 1 |
| fliF | Flagellar M-ring protein | 4,960,263 | C | A | Nonsynonymous_Coding | Ggc/Tgc | G70C | 0 | 0 | 0 | 0 | 0 | 0 | 0 | 0 | 0 | 0 | 0 | 0 | 0 | 0 | 1 | 0 | 0 | 1 | 1 | 0.5 | 0.5 |
| flgK | Flagellar hook-associated protein FlgK | 4,974,437 | CAG | AG | Frame shift | ctg/ | L389 | | | | | | | | | | | | | | | | 0 | 0 | 0 | 1 | 1 | 1 |
| astB | N-succinylarginine dihydrolase | 5,087,105 | A | T | Nonsynonymous_Coding | cTg/cAg | L73Q | 0 | 0 | 0 | 0 | 0 | 0 | 0 | 0 | 0 | 0 | 0 | 1 | 0 | 1 | 0 | 0 | 0 | 2 | 1 | 0.2222 | 0.2222 |
| Anonymous | | 5,119,601 | A | G | Intergenic | | | 0 | 0 | 0 | 0 | 0 | 0 | 0 | 0 | 0 | 0 | 0 | 1 | 0 | 0 | 0 | 0 | 0 | 1 | 1 | 0.5 | 0.5 |
| Anonymous | | 5,257,856 | T | C | Intergenic | | | 0 | 0 | 0 | 0 | 0 | 1 | 0 | 0 | 0 | 0 | 0 | 0 | 0 | 0 | 0 | 0 | 1 | 0 | 0.5 | 1 | 0.5 |
| Anonymous | | 5,459,560 | T | G | Intergenic | | | 0 | 0 | 0 | 0 | 0 | 0 | 1 | 0 | 0 | 0 | 0 | 0 | 0 | 0 | 0 | 0 | 1 | 0 | 0.5 | 1 | 0.5 |
| Anonymous | | 5,459,561 | C | A | Intergenic | | | 0 | 0 | 0 | 0 | 0 | 0 | 1 | 0 | 0 | 0 | 0 | 0 | 0 | 0 | 0 | 0 | 1 | 0 | 0.5 | 1 | 0.5 |
| Anonymous | | 5,509,759 | C | T | Intergenic | | | 0 | 0 | 0 | 0 | 0 | 0 | 0 | 0 | 0 | 0 | 0 | 0 | 0 | 1 | 0 | 0 | 0 | 1 | 1 | 0.5 | 0.5 |
| hisZ | ATP phosphoribosyltransferase regulatory subuint | 5,559,943 | T | A | Nonsynonymous_Coding | cAg/cTg | Q185L | 0 | 0 | 0 | 0 | 0 | 1 | 0 | 0 | 0 | 0 | 0 | 0 | 0 | 0 | 0 | 0 | 1 | 0 | 0.5 | 1 | 0.5 |
| PP_4941 | Hypothetical protein | 5,623,261 | A | T | Nonsynonymous_Coding | caT/caA | H159Q | 0 | 0 | 0 | 0 | 0 | 0 | 0 | 0 | 0 | 0 | 1 | 0 | 0 | 0 | 0 | 0 | 0 | 1 | 1 | 0.5 | 0.5 |
| Anonymous | | 5,692,563 | C | T | Intergenic | | | 0 | 0 | 0 | 0 | 0 | 0 | 0 | 0 | 0 | 0 | 1 | 0 | 0 | 0 | 0 | 0 | 0 | 1 | 1 | 0.5 | 0.5 |
| Anonymous | | 5,743,331 | G | A | Intergenic | | | 0 | 0 | 0 | 0 | 0 | 0 | 0 | 0 | 0 | 0 | 0 | 1 | 0 | 0 | 0 | 0 | 0 | 1 | 1 | 0.5 | 0.5 |

| | | | | | | | | | | | | | | | | | | | | | | | | | | | | |
|---|---|---|---|---|---|---|---|---|---|---|---|---|---|---|---|---|---|---|---|---|---|---|---|---|---|---|---|---|
| Anonymous | | 5,743,335 | T | C | Intergenic | | | 0 | 0 | 0 | 0 | 0 | 0 | 0 | 0 | 0 | 0 | 0 | 1 | 0 | 0 | 0 | 0 | 0 | 1 | 1 | 0.5 | 0.5 |
| Anonymous | | 5,743,390 | G | A | Intergenic | | | 0 | 0 | 0 | 0 | 0 | 1 | 0 | 0 | 0 | 0 | 0 | 0 | 0 | 0 | 0 | 0 | 1 | 0 | 0.5 | 1 | 0.5 |
| Anonymous | | 5,743,394 | T | C | Intergenic | | | 0 | 0 | 0 | 0 | 0 | 1 | 0 | 0 | 0 | 0 | 0 | 0 | 0 | 0 | 0 | 0 | 1 | 0 | 0.5 | 1 | 0.5 |
| metW | Methionine biosynthesis protein MetW | 5,821,897 | T | A | Nonsynonymous_Coding | cTg/cAg | L173Q | 1 | 0 | 0 | 0 | 0 | 0 | 0 | 0 | 0 | 0 | 0 | 0 | 0 | 0 | 0 | 1 | 0 | 0 | 0.5 | 0.5 | 1 |
| ubiH | 2-0ctaprenyl--6-methoxyphenyl hydroxylase/ubiquinone biosynthesis hydroxylase | 5,931,283 | C | G | Nonsynonymous_Coding | Gtt/Ctt | V263L | 0 | 0 | 0 | 0 | 0 | 0 | 0 | 0 | 0 | 0 | 0 | 0 | 0 | 0 | 1 | 0 | 0 | 1 | 1 | 0.5 | 0.5 |
| Anonymous | | 5,988,760 | C | G | Intergenic | | | 0 | 0 | 0 | 0 | 0 | 0 | 0 | 0 | 0 | 0 | 0 | 1 | 0 | 1 | 0 | 0 | 0 | 2 | 1 | 0.2222 | 0.2222 |
| Anonymous | | 5,988,763 | C | G | Intergenic | | | 0 | 0 | 0 | 0 | 0 | 0 | 0 | 0 | 0 | 0 | 0 | 1 | 0 | 1 | 0 | 0 | 0 | 2 | 1 | 0.2222 | 0.2222 |
| Anonymous | | 5,988,803 | G | T | Intergenic | | | 0 | 0 | 0 | 0 | 0 | 1 | 1 | 0 | 0 | 0 | 0 | 0 | 0 | 0 | 0 | 0 | 2 | 0 | 0.2222 | 1 | 0.2222 |
| Anonymous | | 5,988,889 | C | A | Intergenic | | | 0 | 0 | 0 | 1 | 1 | 1 | 0 | 0 | 0 | 0 | 1 | 0 | 1 | 0 | 1 | 2 | 1 | 3 | 0.5833 | 0.6032 | 0.2857 |
| Anonymous | | 5,988,894 | C | T | Intergenic | | | 0 | 0 | 0 | 1 | 1 | 1 | 0 | 0 | 0 | 0 | 1 | 0 | 1 | 0 | 1 | 2 | 1 | 3 | 0.5833 | 0.6032 | 0.2857 |
| Anonymous | | 5,988,895 | A | G | Intergenic | | | 0 | 0 | 0 | 1 | 1 | 1 | 0 | 0 | 0 | 0 | 1 | 0 | 1 | 0 | 1 | 2 | 1 | 3 | 0.5833 | 0.6032 | 0.2857 |
| Anonymous | | 5,988,905 | T | C | Intergenic | | | 1 | 0 | 0 | 1 | 0 | 1 | 0 | 1 | 0 | 1 | 0 | 1 | 1 | 0 | 1 | 2 | 3 | 3 | 0.6032 | 0.6032 | 1 |
| Anonymous | | 5,988,910 | CA | C | Intergenic | | | 1 | 0 | 0 | 1 | 0 | 1 | 0 | 1 | 0 | 1 | 0 | 1 | 1 | 0 | 1 | 2 | 3 | 3 | 0.6032 | 0.6032 | 1 |
| Anonymous | | 5,988,951 | G | C | Intergenic | | | 0 | 1 | 1 | 1 | 1 | 1 | 1 | 1 | 0 | 0 | 1 | 1 | 0 | 1 | 1 | 4 | 3 | 4 | 0.5833 | 1 | 0.5833 |
| Anonymous | | 5,988,954 | G | C | Intergenic | | | 0 | 1 | 1 | 1 | 1 | 1 | 1 | 1 | 0 | 0 | 1 | 1 | 1 | 1 | 1 | 4 | 3 | 5 | 0.5833 | 1 | 0.2222 |
| copA | Copper resistance protein A | 6,132,511 | C | T | Nonsynonymous_Coding | Ggc/Agc | G448S | 0 | 0 | 0 | 0 | 1 | 1 | 0 | 0 | 0 | 1 | 0 | 0 | 0 | 1 | 0 | 1 | 2 | 1 | 0.5833 | 1 | 0.5833 |
| copA | Copper resistance protein A | 6,131,525 | T | C | Nonsynonymous_Coding | gAg/gGg | E443G | 0 | 0 | 0 | 0 | 0 | 1 | 0 | 0 | 1 | 0 | 1 | 0 | 1 | 1 | 0 | 0 | 2 | 3 | 0.2222 | 0.08333 | 0.6032 |
| copA | Copper resistance protein A | 6,132,542 | A | G | Synonymous_Coding | gaT/gaC | D437 | 0 | 0 | 0 | 0 | 0 | 0 | 0 | 0 | 0 | 0 | 1 | 0 | 0 | 0 | 0 | 0 | 0 | 1 | 1 | 0.5 | 0.5 |
