## Supplementary Data S3 for "Experimental Evolution of *Pseudomonas putida* under Silver Ion versus Nanoparticle Stress"

SI for Feng Dong, Ana C. Quevedo, Xiang Wang, Eugenia Valsami-Jones & Jan-Ulrich Kreft (2020).Experimental Evolution of Pseudomonas putida under Silver Ion versus Nanoparticle Stress.

List of mutations in the coding regions for the 15 populations from the main evolution experiment and corresponding gene products. The gene name, codon substrate and amino acid change were obtained by aligning the reads from the evolved populations to the *Pseudomonas putida* KT2440 reference genome. The annotations of product type, EC number and product were extracted from the MicroScope platform. The capitalized bases in the codon substrate are the changed nucleotides. A modified Fisher's exact test was used to calculate the mid-p-values for association of mutation with treatment, p-values ≤0.05 are marked in red.

| Gene name | Gene position | Mutation Position | Product type | EC Number | Genome function and product | Reference genome | Mutation | Effect | Codon substrate | Amino acid change | Amino acid length | Mutation call: Control | | | | | $Ag^+$-population | | | | | AgNP-population | | | | | Total number of parallel mutations | | | p-values for association | | |
|---|---|---|---|---|---|---|---|---|---|---|---|---|---|---|---|---|---|---|---|---|---|---|---|---|---|---|---|---|---|---|---|---|
| | | | | | | | | | | | | C1 | C2 | C3 | C4 | C5 | C1 | C2 | C3 | C4 | C5 | C1 | C2 | C3 | C4 | C5 | Control | $Ag^+$-population | AgNP-population | $Ag^+$-population vs Control | AgNP-population vs Control | $Ag^+$-population vs AgNP-population |
| PP_0174 | 225775-226593 | 226226 | N/A | N/A | Conserved protein of unknown function | T | A | Nonsynonymous | cTg/cAg | L151Q | 272 | 0 | 0 | 0 | 0 | 0 | 0 | 0 | 0 | 1 | 0 | 0 | 0 | 0 | 0 | 0 | 0 | 1 | 0 | 0.5 | 1 | 0.5 |
| PP_0188 | 239706-240833 | 240531 | Putative enzyme | N/A | Putative Uroporphyrin-III C-methyltransferase | G | C | Nonsynonymous | Gcc/Ccc | A276P | 375 | 1 | 0 | 0 | 0 | 0 | 0 | 0 | 0 | 0 | 0 | 0 | 0 | 0 | 0 | 0 | 1 | 0 | 0 | 0.5 | 0.5 | 1 |
| PP_0188 | 239706-240833 | 240536 | Putative enzyme | N/A | Putative Uroporphyrin-III C-methyltransferase | G | C | Synonymous | ggG/ggC | G277 | 375 | 1 | 0 | 0 | 0 | 0 | 0 | 0 | 0 | 0 | 0 | 0 | 0 | 0 | 0 | 0 | 1 | 0 | 0 | 0.5 | 0.5 | 1 |
| tauB-II | 286334-287122 | 287067 | Transporter | 3.6.3.36 | Taurine transporter subunit; ATP-binding component of ABC superfamily | G | C | Nonsynonymous | cCg/cGg | P19R | 262 | 0 | 0 | 0 | 0 | 0 | 0 | 0 | 1 | 0 | 0 | 0 | 0 | 0 | 0 | 0 | 0 | 1 | 0 | 0.5 | 1 | 0.5 |
| tauB-II | 286334-287122 | 287068 | Transporter | 3.6.3.36 | Taurine transporter subunit; ATP-binding component of ABC superfamily | G | C | Nonsynonymous | Ccg/Gcg | P19A | 262 | 0 | 0 | 0 | 0 | 0 | 0 | 0 | 1 | 0 | 0 | 0 | 0 | 0 | 0 | 0 | 0 | 1 | 0 | 0.5 | 1 | 0.5 |
| envZ | 301836-303149 | 302935 | Regulator | N/A | Osmolarity sensor protein EnvZ, sensory histidine kinase in two-component regulatory system with OmpR | G | C | Nonsynonymous | gGt/gCt | G367A | 437 | 0 | 0 | 0 | 0 | 0 | 0 | 0 | 0 | 0 | 0 | 0 | 1 | 0 | 0 | 0 | 0 | 0 | 1 | 1 | 0.5 | 0.5 |
| envZ | 301836-303149 | 303121 | Regulator | N/A | Osmolarity sensor protein EnvZ, sensory histidine kinase in two-component regulatory system with OmpR | G | GGCT | Codon change/Codon insertion | ccg/cTGCcg | P433LP | 437 | 0 | 0 | 0 | 0 | 0 | 0 | 0 | 0 | 0 | 0 | 1 | 1 | 1 | 0 | 0 | 0 | 0 | 3 | 1 | 0.08333 | 0.08333 |
| thiL | 604688-605656 | 605121 | Enzyme | 2.7.4.16 | Thiamin monophosphate kinase | G | A | Nonsynonymous | cGc/cAc | R145H | 322 | 0 | 0 | 0 | 0 | 0 | 0 | 0 | 1 | 0 | 0 | 0 | 0 | 0 | 0 | 0 | 0 | 1 | 0 | 0.5 | 1 | 0.5 |
| thiL | 604688-605656 | 605126 | Enzyme | 2.7.4.16 | Thiamin monophosphate kinase | A | T | Nonsynonymous | Agc/Tgc | S147C | 322 | 0 | 0 | 0 | 0 | 0 | 0 | 0 | 1 | 0 | 0 | 0 | 0 | 0 | 0 | 0 | 0 | 1 | 0 | 0.5 | 1 | 0.5 |
| thiL | 604688-605656 | 605131 | Enzyme | 2.7.4.16 | Thiamin monophosphate kinase | T | C | Synonymous | ggT/ggC | G148 | 322 | 0 | 0 | 0 | 0 | 0 | 0 | 0 | 1 | 0 | 0 | 0 | 0 | 0 | 0 | 0 | 0 | 1 | 0 | 0.5 | 1 | 0.5 |
| hemH | 862867-863883 | 863376 | Enzyme | 4.99.1.1 | Ferrochelatase | A | T | Synonymous | gcA/gcT | A170 | 338 | 0 | 0 | 1 | 1 | 0 | 1 | 0 | 1 | 0 | 0 | 0 | 0 | 1 | 0 | 0 | 2 | 2 | 1 | 1 | 0.5833 | 0.5833 |
| PP_0861 | 997827-1000118 | 999625 | Receptor/putative transporter | N/A | Outer membrane ferric siderophore receptor/TonB-dependent siderophore receptor | C | G | Nonsynonymous | gGc/gCc | G182A | 780 | 0 | 0 | 0 | 0 | 0 | 1 | 0 | 0 | 0 | 0 | 0 | 0 | 0 | 0 | 0 | 0 | 1 | 0 | 0.5 | 1 | 0.5 |
| PP_0904 | 1044293-1044994 | 1044757 | ORF of unknown function | N/A | InaA protein | CAGCACGGCTGCCTGTATGGCA | C | Codon change/Codon deletion | ggctgcctgtatggcaagcacgta/gta | GCLYGKHV156V | 233 | 0 | 0 | 0 | 0 | 0 | 0 | 0 | 0 | 0 | 0 | 0 | 0 | 0 | 1 | 0 | 0 | 0 | 1 | 1 | 0.5 | 0.5 |
| xcpY | 1202172-1203260 | 1202854 | N/A | N/A | Type II secretion pathway protein XcpY | A | C | Nonsynonymous | cAg/cCg | Q228P | 362 | 0 | 0 | 0 | 0 | 0 | 1 | 0 | 0 | 0 | 0 | 0 | 0 | 0 | 0 | 0 | 0 | 1 | 0 | 0.5 | 1 | 0.5 |
| dtcD-II | 1219977-1221305 | 1221046 | Regulator | N/A | C4-dicarboxylate transport transcriptional regulatory protein | C | A | Nonsynonymous | gGc/gTc | G87V | 442 | 1 | 0 | 0 | 0 | 0 | 0 | 0 | 0 | 0 | 0 | 0 | 0 | 0 | 0 | 0 | 1 | 0 | 0 | 0.5 | 0.5 | 1 |
| PP_16SE | 1325503-1327024 | 1326938 | N/A | N/A | 16S ribosomal RNA | | | Intergenic | | | | 1 | 0 | 0 | 0 | 0 | 1 | 1 | 1 | 1 | 1 | 0 | 0 | 0 | 0 | 0 | 1 | 5 | 0 | 0.02381 | 0.5 | 0.003968 |
| PP_1175 | 1350104-1350199 | 1350104 | ORF or unknown function | N/A | Conserved protein of unknown function (fragment) | C | A | Stop lost/Splice site region | taG/taT | *32Y | 31 | 0 | 0 | 0 | 0 | 0 | 1 | 0 | 0 | 0 | 0 | 0 | 0 | 0 | 0 | 0 | 0 | 1 | 0 | 0.5 | 1 | 0.5 |
| PP_1175 | 1350104-1350199 | 1350119 | ORF or unknown function | N/A | Conserved protein of unknown function (fragment) | A | G | Synonymous | agT/agC | S27 | 31 | 0 | 0 | 0 | 1 | 0 | 0 | 0 | 0 | 0 | 0 | 0 | 0 | 0 | 0 | 0 | 1 | 0 | 0 | 0.5 | 0.5 | 1 |
| PP_1195 | 1369205-1370923 | 1370775 | ORF of unknown function | N/A | Conserved exported protein of unknown function | GGGTCGAAAA | G | Codon change/Codon deletion | gaaaaggtcgag/gag | EKVE535E | 572 | 0 | 0 | 0 | 0 | 0 | 0 | 0 | 0 | 0 | 0 | 0 | 0 | 0 | 0 | 1 | 0 | 0 | 1 | 1 | 0.5 | 0.5 |
| PP_1256 | 1434386-1435963 | 1435822 | Putative enzyme | 1.2.1.26 | Putative alpha-ketoglutarate semialdehyde dehydrogenase | A | T | Nonsynonymous | Tgc/Agc | C48S | 525 | 0 | 0 | 0 | 0 | 0 | 0 | 0 | 0 | 1 | 0 | 0 | 0 | 0 | 0 | 0 | 0 | 1 | 0 | 0.5 | 1 | 0.5 |
| PP_1325 | 1509749-1511566 | 1511070 | Lipoprotein | N/A | Putative lipoprotein | T | A | Nonsynonymous | cAg/cTg | Q166L | 605 | 0 | 0 | 0 | 0 | 0 | 0 | 0 | 0 | 0 | 0 | 0 | 0 | 1 | 0 | 0 | 0 | 0 | 1 | 1 | 0.5 | 0.5 |
| ftsZ/rna60/PP_mr19 | 1528978-1530174 | 1530160 | Cell process | N/A | GTP-binding tubulin-like cell division protein | C | T | Nonsynonymous | Cgt/Tgt | R395C | 398 | 0 | 0 | 0 | 0 | 0 | 0 | 1 | 0 | 0 | 0 | 1 | 1 | 1 | 1 | 1 | 0 | 1 | 5 | 0.5 | 0.003968 | 0.02381 |
| ftsZ/rna60/PP_1344 | 1528978-1530174 | 1530173 | Cell process | N/A | GTP-binding tubulin-like cell division protein | A | C | Stop lost/Splice site region | tAa/tCa | *399S | 398 | 1 | 1 | 1 | 1 | 1 | 1 | 1 | 1 | 1 | 1 | 0 | 0 | 0 | 0 | 0 | 5 | 5 | 0 | 1 | 0.003968 | 0.003968 |
| PP_1344 | 1531246-1531875 | 1531780 | ORF of unknown function | N/A | conserved protein of unknown function | G | T | Nonsynonymous | Ccc/Acc | P36T | 209 | 0 | 0 | 0 | 0 | 0 | 0 | 0 | 0 | 0 | 0 | 0 | 0 | 1 | 0 | 0 | 0 | 0 | 1 | 1 | 0.5 | 0.5 |
| gacS | 1842040-1844793 | 1843091 | N/A | 2.7.13.3 | Sensor protein GacS | A | C | Nonsynonymous | gTg/gGg | V568G | 917 | 1 | 1 | 1 | 0 | 1 | 1 | 1 | 1 | 1 | 1 | 0 | 0 | 0 | 0 | 1 | 4 | 5 | 1 | 1 | 0.1071 | 0.02381 |
| gacS | 1842040-1844793 | 1844166 | N/A | 2.7.13.3 | Sensor protein GacS | T | A | Stop gain | Aag/Tag | K210* | 917 | 0 | 0 | 0 | 0 | 0 | 0 | 1 | 0 | 0 | 0 | 1 | 1 | 1 | 1 | 0 | 0 | 1 | 4 | 0.5 | 0.02381 | 0.1071 |
| PP_1666 | 1862315-1863382 | 1863240 | ORF of unknown function | N/A | Conserved exported protein of unknown function | T | A | Nonsynonymous | cTg/cAg | L309Q | 355 | 0 | 0 | 0 | 0 | 0 | 0 | 0 | 0 | 0 | 0 | 0 | 0 | 1 | 0 | 0 | 0 | 0 | 1 | 1 | 0.5 | 0.5 |
| PP_1666 | 1862315-1863382 | 1863278 | ORF of unknown function | N/A | Conserved exported protein of unknown function | G | C | Nonsynonymous | Gcg/Ccg | A322P | 355 | 0 | 0 | 1 | 0 | 0 | 0 | 0 | 0 | 0 | 0 | 0 | 0 | 0 | 0 | 0 | 1 | 0 | 0 | 0.5 | 0.5 | 1 |
| PP_1703 | 1899450-1903475 | 1901761 | Enzyme | 1.7.99.4 | Assimilatory nitrate reductase/sulfite reductase | A | T | Nonsynonymous | gAc/gTc | D787V | 1357 | 0 | 0 | 0 | 0 | 0 | 0 | 0 | 1 | 0 | 0 | 0 | 0 | 0 | 0 | 0 | 0 | 1 | 0 | 0.5 | 1 | 0.5 |
| PP_5491 | 2187611-2188525 | 2187722 | ORF of unknown function | N/A | Conserved protein of unknown function with SEC-C motif domain | GT | G | Intergenic | | | | 0 | 0 | 0 | 0 | 0 | 0 | 1 | 0 | 0 | 0 | 0 | 0 | 0 | 0 | 0 | 0 | 1 | 0 | 0.5 | 1 | 0.5 |
| PP_2068 | 2353254-2354531 | 2354379 | N/A | N/A | Putative Multidrug efflux MFS membrane fusion protein | T | G | Nonsynonymous | aaA/aaC | K51N | 425 | 0 | 0 | 0 | 1 | 0 | 0 | 0 | 0 | 0 | 0 | 0 | 0 | 0 | 0 | 0 | 1 | 0 | 0 | 0.5 | 0.5 | 1 |
| PP_2397 | 2742905-2743561 | 2742917 | N/A | N/A | EF hand domain protein | C | T | Nonsynonymous | Cac/Tac | H5Y | 218 | 0 | 0 | 0 | 0 | 0 | 1 | 0 | 0 | 0 | 0 | 0 | 0 | 0 | 0 | 0 | 0 | 1 | 0 | 0.5 | 1 | 0.5 |
| PP_2397 | 2742905-2743561 | 2742918 | N/A | N/A | EF hand domain protein | A | G | Nonsynonymous | cAc/cGc | H5R | 218 | 0 | 0 | 0 | 0 | 0 | 1 | 0 | 0 | 0 | 0 | 0 | 0 | 0 | 0 | 0 | 0 | 1 | 0 | 0.5 | 1 | 0.5 |
| PP_2478 | 2824388-2826580 | 2824518 | N/A | N/A | Putative Isoquinoline 1-oxidoreductase, beta subunit | A | C | Nonsynonymous | gAg/gCg | E44A | 730 | 0 | 0 | 0 | 0 | 1 | 0 | 0 | 0 | 0 | 0 | 0 | 0 | 0 | 0 | 0 | 1 | 0 | 0 | 0.5 | 0.5 | 1 |
| PP_2638 | 3019463-3022981 | 3021030 | N/A | N/A | Putative Cellulose synthase operon C protein | A | T | Nonsynonymous | cAg/cTg | Q523L | 1172 | 0 | 0 | 1 | 0 | 0 | 0 | 0 | 1 | 0 | 0 | 1 | 0 | 0 | 1 | 0 | 1 | 1 | 2 | 1 | 0.5833 | 0.5833 |
| PP_2757 | 3141084-3142055 | 3141492 | N/A | N/A | Putative Sugar-binding protein,imilar to Periplasmic binding protein/LacI transcriptional regulator | GT | G | Frame shift | gtc/ | V137 | 323 | 0 | 0 | 0 | 0 | 0 | 0 | 0 | 0 | 0 | 0 | 0 | 1 | 0 | 0 | 0 | 0 | 0 | 1 | 1 | 0.5 | 0.5 |
| PP_2758 | 3142052-3143080 | 3142272 | N/A | N/A | Putative Ribose ABC transporter, periplasmic ribose-binding protein | C | T | Nonsynonymous | gCg/gTg | A74V | 342 | 0 | 0 | 0 | 0 | 0 | 0 | 0 | 0 | 0 | 0 | 1 | 1 | 1 | 1 | 0 | 0 | 0 | 4 | 1 | 0.02381 | 0.02381 |
| PP_3045 | 3429783-3430946 | 3430349 | N/A | N/A | Putative ClpP protease | G | C | Synonymous | tcG/tcC | S189 | 387 | 0 | 0 | 0 | 0 | 0 | 0 | 0 | 0 | 0 | 0 | 0 | 0 | 1 | 0 | 0 | 0 | 0 | 1 | 1 | 0.5 | 0.5 |
| PP_3221 | 3656389-3657297 | 3656539 | N/A | N/A | ABC transporter, permease protein | T | G | Synonymous | ccA/ccC | P253 | 302 | 0 | 0 | 0 | 0 | 0 | 0 | 0 | 0 | 0 | 0 | 1 | 0 | 0 | 0 | 0 | 0 | 0 | 1 | 1 | 0.5 | 0.5 |
| PP_5743/PP_3334 | 3773298-3774305/3774012-3774296 | 3774247 | putative membrane component/putative regulator | N/A | Putative TonB-dependent receptor protein/putative lysR family transcriptional regulator (fragment) | C | G | Intragenic | | | | 1 | 0 | 0 | 0 | 0 | 1 | 0 | 0 | 0 | 0 | 0 | 0 | 0 | 0 | 0 | 1 | 1 | 0 | 1 | 0.5 | 0.5 |
| oprN | 3881342-3882757 | 3881829 | N/A | N/A | Multidrug efflux RND outer membrane protein OprN | A | T | Nonsynonymous | cAg/cTg | Q163L | 471 | 0 | 1 | 0 | 0 | 0 | 0 | 0 | 0 | 0 | 0 | 0 | 0 | 0 | 0 | 0 | 1 | 0 | 0 | 0.5 | 0.5 | 1 |
| PP_3563 | 4041913-4042764 | 4042322 | ORF of unknown function | N/A | Conserved exported protein of unknown function | T | G | Nonsynonymous | cTc/cGc | L137R | 283 | 0 | 0 | 0 | 0 | 0 | 0 | 1 | 0 | 0 | 0 | 0 | 0 | 0 | 0 | 0 | 0 | 1 | 0 | 0.5 | 1 | 0.5 |
| PP_3573 | 4053427-4054746 | 4053476 | N/A | N/A | Putative Monooxygenase/ similar to FAD-dependent oxidoreductase | A | T | Nonsynonymous | cAg/cTg | Q17L | 439 | 0 | 1 | 0 | 0 | 1 | 0 | 0 | 1 | 0 | 0 | 0 | 0 | 1 | 0 | 0 | 2 | 1 | 1 | 0.5833 | 0.5833 | 1 |
| yrpB | 4352875-4353960 | 4353296 | Enzyme | 1.13.12.16 | 2-nitropropane dioxygenase | C | G | Nonsynonymous | cCg/cGg | P141R | 361 | 0 | 0 | 0 | 0 | 0 | 0 | 0 | 1 | 0 | 0 | 1 | 0 | 0 | 0 | 0 | 0 | 1 | 1 | 0.5 | 0.5 | 1 |
| clpA | 4517249-4519519 | 4518633 | Enzyme | 3.4.21.92 | ATP-dependent serine protease,chaperone activity/class III stress response-related ATPase, AAA+ superfamily | C | T | Nonsynonymous | gGc/gAc | G296D | 756 | 0 | 0 | 0 | 0 | 0 | 0 | 0 | 0 | 0 | 0 | 0 | 0 | 0 | 0 | 1 | 0 | 0 | 1 | 1 | 0.5 | 0.5 |
| idh | 4522152-4524377 | 4522716 | Enzyme | 1.1.1.- | Isocitrate dehydrogenase | G | A | Nonsynonymous | Gca/Aca | A189T | 741 | 0 | 0 | 1 | 0 | 0 | 0 | 0 | 0 | 0 | 0 | 0 | 0 | 0 | 0 | 0 | 1 | 0 | 0 | 0.5 | 0.5 | 1 |
| malQ | 4568071-4570140 | 4568619 | Enzyme | 2.4.1.25 | 4-alpha-glucanotransferase | C | G | Synonymous | gcC/gcG | A183 | 689 | 1 | 0 | 0 | 0 | 0 | 0 | 0 | 0 | 0 | 0 | 0 | 0 | 0 | 0 | 0 | 1 | 0 | 0 | 0.5 | 0.5 | 1 |
| mccB | 4589319-4590926 | 4590049 | Enzyme | 6.4.1.4 | Methylcrotonyl-CoA carboxylase biotin-containing subunit beta | T | C | Nonsynonymous | gTc/gCc | V244A | 535 | 0 | 0 | 0 | 0 | 0 | 0 | 0 | 0 | 0 | 0 | 1 | 0 | 0 | 0 | 0 | 0 | 0 | 1 | 1 | 0.5 | 0.5 |
| uvrY | 4635117-4635755 | 4635590 | Regulator | N/A | Two-componenent system BarA/UvrY - regulatory subunit | T | C | Nonsynonymous | Aaa/Gaa | K56E | 212 | 1 | 1 | 1 | 1 | 1 | 1 | 1 | 1 | 1 | 1 | 0 | 0 | 0 | 0 | 1 | 5 | 5 | 1 | 1 | 0.02381 | 0.02381 |
| ccmE | 4914695-4915150 | 4914726 | Factor | N/A | Cytochrome c-type biogenesis protein CcmE | T | A | Nonsynonymous | cAg/cTg | Q142L | 151 | 0 | 0 | 0 | 0 | 0 | 0 | 0 | 0 | 0 | 0 | 1 | 0 | 0 | 0 | 0 | 0 | 0 | 1 | 1 | 0.5 | 0.5 |
| fliF | 4958692-4960470 | 4960263 | Structure | N/A | Flagellar M-ring protein | C | A | Nonsynonymous | Ggc/Tgc | G70C | 592 | 0 | 0 | 0 | 0 | 0 | 0 | 0 | 0 | 0 | 0 | 0 | 0 | 0 | 0 | 1 | 0 | 0 | 1 | 1 | 0.5 | 0.5 |
| astB | 5085973-5087322 | 5087105 | Enzyme | 3.5.3.23 | N-succinylarginine dihydrolase | A | T | Nonsynonymous | cTg/cAg | L73Q | 449 | 0 | 0 | 0 | 0 | 0 | 0 | 0 | 0 | 0 | 0 | 0 | 1 | 0 | 1 | 0 | 0 | 0 | 2 | 1 | 0.2222 | 0.2222 |
| hisZ | 5559309-5560496 | 5559946 | Enzyme | 2.4.2.17 | Histidyl-tRNA synthetase-like component of ATP phophoribosyltransferase | T | A | Nonsynonymous | cAg/cTg | Q185L | 395 | 0 | 0 | 0 | 0 | 0 | 1 | 0 | 0 | 0 | 0 | 0 | 0 | 0 | 0 | 0 | 0 | 1 | 0 | 0.5 | 1 | 0.5 |
| PP_4941 | 5622334-5623737 | 5623261 | ORF or unknown function | N/A | Conserved exported protein of unknown function | A | T | Nonsynonymous | caT/caA | H159Q | 467 | 0 | 0 | 0 | 0 | 0 | 0 | 0 | 0 | 0 | 0 | 1 | 0 | 0 | 0 | 0 | 0 | 0 | 1 | 1 | 0.5 | 0.5 |
| metW | 5821380-5822018 | 5821897 | Enzyme | N/A | Methionine biosynthesis protein MetW | T | A | Nonsynonymous | cTg/cAg | L173Q | 212 | 1 | 0 | 0 | 0 | 0 | 0 | 0 | 0 | 0 | 0 | 0 | 0 | 0 | 0 | 0 | 1 | 0 | 0 | 0.5 | 0.5 | 1 |
| ubiH | 5930870-59320692 | 5931283 | Enzyme | 1.14.-.- | 2-octaprenyl-6-methoxyphenol hydroxylase, FAD/NAD(P)-binding | C | G | Nonsynonymous | Gtt/Ctt | V263L | 399 | 0 | 0 | 0 | 0 | 0 | 0 | 0 | 0 | 0 | 0 | 0 | 0 | 0 | 0 | 1 | 0 | 0 | 1 | 1 | 0.5 | 0.5 |
| copA | 6131843-6133852 | 6132511 | Transporter | N/A | Copper resistance protein A | C | T | Nonsynonymous | Ggc/Agc | G448S | 669 | 0 | 0 | 0 | 0 | 1 | 1 | 0 | 0 | 0 | 1 | 0 | 0 | 0 | 1 | 0 | 1 | 2 | 1 | 0.5833 | 1 | 0.5833 |
| copA | 6131843-6133852 | 6131525 | Transporter | N/A | Copper resistance protein A | T | C | Nonsynonymous | gAg/gGg | E443G | 669 | 0 | 0 | 0 | 0 | 0 | 1 | 0 | 0 | 1 | 0 | 1 | 0 | 1 | 1 | 0 | 0 | 2 | 3 | 0.2222 | 0.08333 | 0.6032 |
| copA | 6131843-6133852 | 6132542 | Transporter | N/A | Copper resistance protein A | A | G | Synonymous | gaT/gaC | D437 | 669 | 0 | 0 | 0 | 0 | 0 | 0 | 0 | 0 | 0 | 0 | 1 | 0 | 0 | 0 | 0 | 0 | 0 | 1 | 1 | 0.5 | 0.5 |
